# Haplotype tagging reveals parallel formation of hybrid races in two butterfly species

**DOI:** 10.1101/2020.05.25.113688

**Authors:** Joana I. Meier, Patricio A. Salazar, Marek Kučka, Robert William Davies, Andreea Dréau, Ismael Aldás, Olivia Box Power, Nicola J. Nadeau, Jon R. Bridle, Campbell Rolian, Nicholas H. Barton, W. Owen McMillan, Chris D. Jiggins, Yingguang Frank Chan

## Abstract

Genetic variation segregates as linked sets of variants, or haplotypes. Haplotypes and linkage are central to genetics and underpin virtually all genetic and selection analysis. And yet, genomic data often lack haplotype information, due to constraints in sequencing technologies. Here we present “haplotagging”, a simple, low-cost linked-read sequencing technique that allows sequencing of hundreds of individuals while retaining linkage information. We apply haplotagging to construct megabase-size haplotypes for over 600 individual butterflies (*Heliconius erato* and *H. melpomene*), which form overlapping hybrid zones across an elevational gradient in Ecuador. Haplotagging identifies loci controlling distinctive high- and lowland wing color patterns. Divergent haplotypes are found at the same major loci in both species, while chromosome rearrangements show no parallelism. Remarkably, in both species the geographic clines for the major wing pattern loci are displaced by 18 km, leading to the rise of a novel hybrid morph in the centre of the hybrid zone. We propose that shared warning signalling (Müllerian mimicry) may couple the cline shifts seen in both species, and facilitate the parallel co-emergence of a novel hybrid morph in both co-mimetic species. Our results show the power of efficient haplotyping methods when combined with large-scale sequencing data from natural populations.

**One-sentence summary:** Haplotagging, a novel linked-read sequencing technique that enables whole genome haplotyping in large populations, reveals the formation of a novel hybrid race in parallel hybrid zones of two co-mimicking *Heliconius* butterfly species through strikingly parallel divergences in their genomes.

## Introduction

Understanding how changes in DNA sequence affect traits and shape the evolution of populations and species has been a defining goal in genetics and evolution (*1*-*3*). DNA is naturally organized in the genome as long molecules consisting of linked chromosome segments. Linkage is a core concept in genetics: in genetic mapping, geneticists map causal variants not by tracking the actual mutation, but through many otherwise neutral and unremarkable linked variants. Likewise, the detection of selection relies on observing hitchhiking of linked variants, rather than seeing the mutation itself. This recognition makes it all the more paradoxical that haplotype information is routinely omitted from most genomic studies, as a technical compromise. Lacking haplotype information not only complicates analysis, but also precludes accurate ancestry reconstruction, detection of allele-specific expression (*4*) and chromosome rearrangements, and reduces power to detect selective sweeps, even entirely missing them when multiple haplotypes sweep together (*5*). Instead of sequencing genomes as haplotypes, short-read sequencing produces 150 bp reads. Until affordable long-read platforms become sufficiently reliable, this lack of haplotype context will continue to impact mapping and genomic studies, particularly those in non-model organisms.

One way to simplify haplotype reconstruction and inference from sequencing data is to avoid discarding haplotype information in the first place. A promising emerging technique is thus linked-read (LR) sequencing (*6*-*9*), which preserves long-range information via molecular barcoding of long DNA molecules before sequencing. Individual short reads can then be linked via a shared barcode to reconstruct the original haplotype. However, existing options all suffer from high cost, poor scalability and/or require custom sequencing primers or settings that have thus far prevented them from being applied as the default sequencing platform (Tables S1–2). If linked-read sequencing can become scalable and affordable, it would significantly advance genetics by enabling, for the first time, the “*haplotyping*” of entire populations, i.e., the sequencing and systematic discovery of genomic variants as haplotypes in hundreds or even thousands of samples in model and non-model organisms alike.

Here we describe a novel solution called “haplotagging”, a simple and rapid protocol for linked-read (LR) sequencing. Importantly, haplotagging maintains full compatibility with standard Illumina sequencing and can easily scale to large populations with no extra costs. We demonstrate this in three steps. First, we show that direct haplotyping using haplotagging is robust in single human and mouse samples with known haplotypes (“phases”). Next, we show the feasibility of population haplotyping in 245 mice, even with very low-coverage LR sequencing. Finally, we apply haplotagging to investigate the emergence of a novel hybrid morph in a hybrid zone system in Ecuador featuring 670 individuals of two species of *Heliconius* butterflies.

### Direct haplotype tagging

Haplotagging is a bead-based protocol for the production of linked-read DNA sequencing libraries. Haplotagging works by molecular barcoding of long, kilobase-spanning DNA molecules to generate short fragments for sequencing. In solution, DNA molecules tend to wrap around a single bead, a property that can be exploited for constructing linked-read libraries (*8, 9*). Each haplotagging bead is tethered with Tn5 transposase carrying one of 85 million molecular barcodes directly integrated into an otherwise standard Nextera Tn5 transposon adaptor (Fig. 1A; fig. S1; Table S3). In a single transposition reaction, microbead-tethered Tn5 transposase transfers the barcoded sequencing adaptors into the long DNA molecule. A tube of beads carrying millions of unique molecular barcodes can be used to tag a pool of DNA molecules, each carrying its bead-specific barcode. Following sequencing, unique long-range haplotypes can be reconstructed from each DNA molecule (Fig. 1A).

**Fig. 1.**
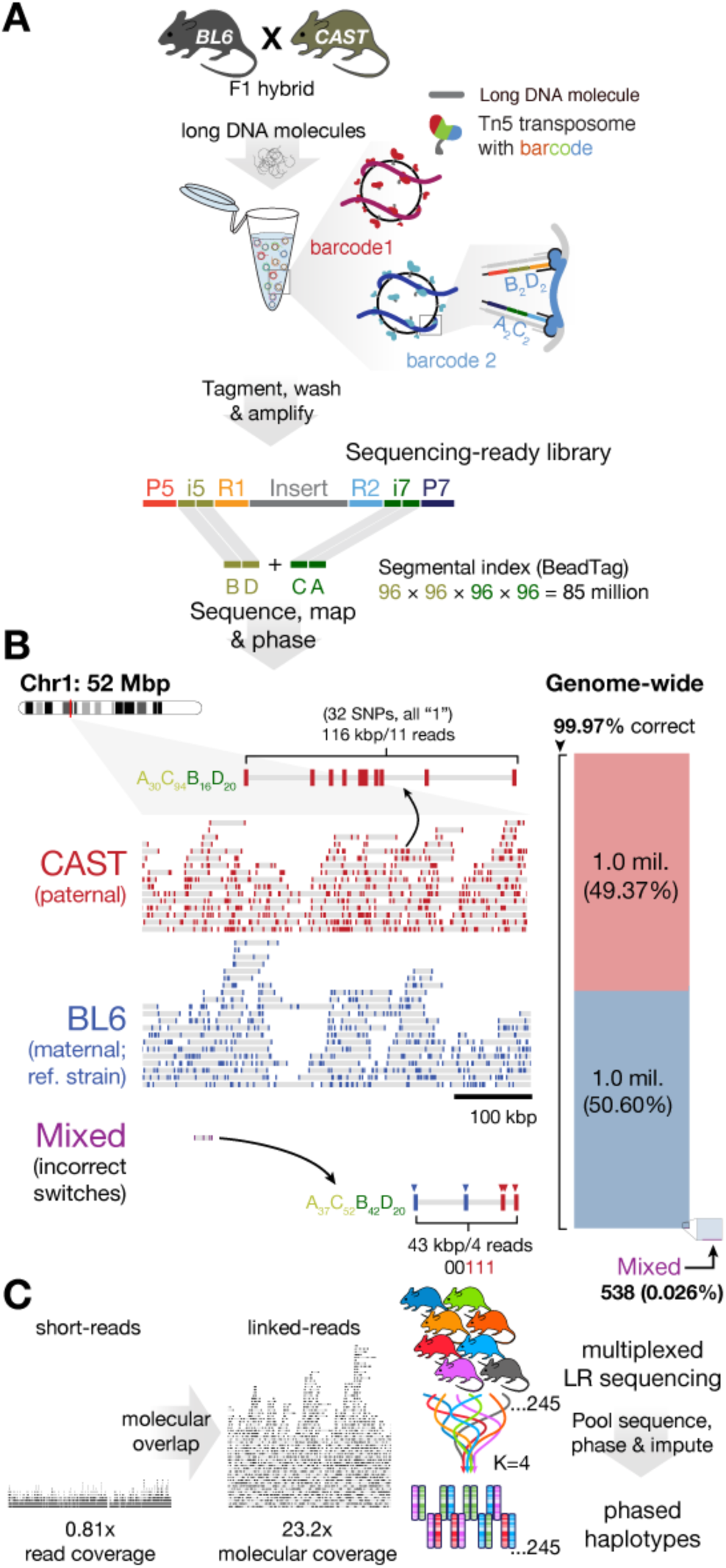
Haplotagging enables population-scale linked-read sequencing. (**A**) Principles of haplotagging. Microbeads coated with barcoded transposon adaptors enable simultaneous molecular barcoding and Tn5-mediated fragmentation of long DNA molecules into sequencing-ready libraries after PCR amplification, all in a single tube. This technique takes advantage of the tendency of DNA to interact only with a single bead in solution (inset). A key feature of haplotagging is that each bead is uniformly coated with a single segmental barcode combination (“beadTag”) made up of 4 segments of 96 barcodes each (designated “B”, “D”, olive; and “C”, “A”, green, at the standard i5/7 index positions of the Illumina Nextera design). Across beads, the four segments represent up to 96^4^, or 85 million beadTags. Thus, DNA wrapped around a single bead can be reconstructed from individual short reads that share the same beadTag. (**B**) Haplotagging in an F1 hybrid mouse between the reference strain C57BL/6 (BL6) and CAST/EiJ (CAST), with detailed view at Chr1: 52-52.5 megabase (Mbp). Each molecule is represented by a grey bar connecting short-reads (colored bars for CAST, red; or BL6, blue) sharing a single beadTag, e.g., A_30_C_94_B_16_D_20_ tags a 116 kilobase molecule carrying a CAST allele. All but one molecule in this window match perfectly to CAST or BL6 alleles. Genome-wide, 99.97% of all reconstructed molecules correspond to CAST or BL6 haplotypes (2 million correct vs. 538 incorrect molecules). (**C**) Vast expansion in molecular vs. read coverage for whole population haplotyping. Linked-read (LR) molecules typically span tens of kilobases, compared to ∼500 bp of short reads. The increased overlap among molecules often lead to >10-fold increase in molecular coverage (Table S4). In a large population, LR data allow both accurate haplotype reconstruction using pooled read depths and accurate imputation by leveraging linkage information, even with input read coverage reduced to 0.07x (fig. S2B). Bead and Tn5 image modified with permission from Zinkia Entertainment, S.A./Pocoyo.

Haplotagging features three main design improvements over other linked-read (LR) alternatives (Table S1). Firstly, it avoids specialized instrumentation (c.f., microfluidics chips and controller for 10X Genomics’ discontinued Chromium platform). Haplotagging is, in essence, a one-step, 10-minute transposition reaction, followed by PCR. It requires only a magnet and standard molecular biology equipment available in most laboratories. A haplotagging library, in our hands, costs less than 1% of a 10X Genomics Chromium library, and despite featuring long-range haplotype information, costs about 1/20^th^ as much as a Nextera DNAFlex short-read library (Table S2). Secondly, we designed the segmental beadTag barcode and the protocol with scalability and high-order multiplexing in mind. A single person can thus prepare and sequence hundreds of uniquely barcoded libraries within weeks. Last but not least, one of the major design challenges we have solved in haplotagging is encoding 85 million barcodes and maintaining full compatibility with standard Illumina TruSeq sequencing that is available at most sequencing facilities, even when pooled with other library types (Fig. 1A).

To test the recovery of molecular haplotypes, we performed haplotagging on high molecular weight DNA from an F1 hybrid mouse between two inbred lab strains with known sequence differences, CAST/EiJ (CAST) and C57BL/6N (BL6 is the genome reference strain). We could assign 94.3% of 201 million read-pairs to a beadTag and inferred molecules based on beadTag sharing (Fig. 1B). Across the genome, we found that 99.97% of phase-informative molecules accurately capture the parental haplotypes with exclusively BL6 (reference, maternal) or CAST (alternate, paternal) alleles at multiple SNPs (Fig. 1B), with even representation of both alleles (50.6% and 49.4%, respectively; about 1.0 million molecules each on autosomes). Many of these molecules span many kilobases (kbp), up to as much as 415 kbp (N_50_ = 42.1 kbp; Table S4). These results provide strong evidence that haplotagging can accurately capture and reconstruct haplotypes.

Using these data, we phased nearly all heterozygous SNPs (99.74%) using HAPCUT2 (*10*) into large, megabase-spanning phased haplotype blocks (N_50_ = 15.5 Mbp; fig. S2A; maximum: 61.46 Mbp; Table S4; see Supplementary Notes for phasing performances in additional human and mouse samples, including comparison against other LR platforms).

### Whole population haplotyping

We next tackled haplotype phasing using LR data in large populations. Unlike phasing in single individuals, population phasing can be challenging, because neither the number of haplotypes nor their frequencies are known in advance. To our knowledge this is the first study to apply population phasing using LR data, presumably because such large datasets have not been feasible before.

Our strategy involves leveraging naturally occurring haplotype blocks in populations and trading off linkage against coverage: first we reconstruct the set of haplotypes present in the study population, exploiting the fact that most segregating haplotypes are common such that we can pool reads from many individuals for maximum coverage. Then, we impute across the entire genome in every sample, using linkage from the expanded “molecular coverage” to boost accuracy. For example, there was a ten-fold increase from 12.6× read- to 165.6× molecular coverage in the F1 hybrid mouse, (i.e., each parental haplotype was sampled more than 80 times; Fig. 1C; fig. S2B; Table S4). This strategy dovetails neatly with STITCH, an algorithm for (short) read-based statistical phasing (*11*), which we have adopted to incorporate LR information. The implication of this principle of tandem molecular and statistical phasing is profound: with LR data, we can sequence populations at a fraction of current coverage (and costs), yet still obtain accurate haplotypes for the entire population.

To test this concept, we performed haplotagging on 245 “Longshanks” mice from a 20-generation selective breeding experiment for long tibia length (*12, 13*). We sequenced these mice to an average depth of 0.24×, identified molecules (giving 2.23× molecular coverage), and phased the data using STITCH. We tested the accuracy of genotype imputation by comparing against higher-coverage conventional short-read data for 32 of these mice (2.9× coverage) (*13*). Our results show that genotype imputation using data from haplotagging is remarkably robust (>96% accurate) and remains so even when read coverage is reduced to 0.07× (vs. 0.15× without using LR information; fig. S2B; see the supplementary materials). Compared to short read sequencing without linked-read data, there was a 100-fold expansion in the ability to assign SNPs into linked sets of phased blocks, with an average of 24.1 kbp using LR data (vs. 283 bp otherwise). We achieved these robust results despite having sequenced only about one-tenth as deeply (cf. typically 20– 40x under classical phasing (*14*), or 2–5x when imputing using an external reference haplotype panel (*11, 15*). Importantly, we are no longer dependent on a reference haplotype panel, which does not exist for most study populations. By multiplexing haplotagging libraries of hundreds of samples, users can now perform (ultra)low-coverage Illumina seqencing, achieve 10- to 100-fold deeper LR molecular coverage and thus accurately reconstruct haplotypes for each individual.

### *Parallel* Heliconius *hybrid zones*

We next applied haplotagging to address key evolutionary questions in two *Heliconius* butterfly species in Ecuador. Local collectors have previously noticed the abundance of a morph of putative hybrid origin in Eastern Ecuador. Here, we investigate the patterns of phenotypic and genetic variation across the hybrid zone where this novel morph is found and test the hypothesis that the morph has arisen and spread in parallel in two species that mimic each other.

*Heliconius* butterflies have diversified into many species and subspecies (or “races”) across South and Central America and represent a classic example of adaptive radiation (16). They are toxic and advertise their unpalatability with bright warning coloration. Predators (mainly birds) learn to avoid the warning signal (*17, 18*) and selection favors the locally common pattern (*19*). *Heliconius* species often converge on the same color patterns to reinforce the advertising effect, a phenomenon known as Müllerian mimicry (Fig. 2A; (*17*)). The striking variety notwithstanding, the genetics of these color patterns is remarkably simple, involving only a few loci of large effect (16, *20*-*25*).

**Fig. 2.**
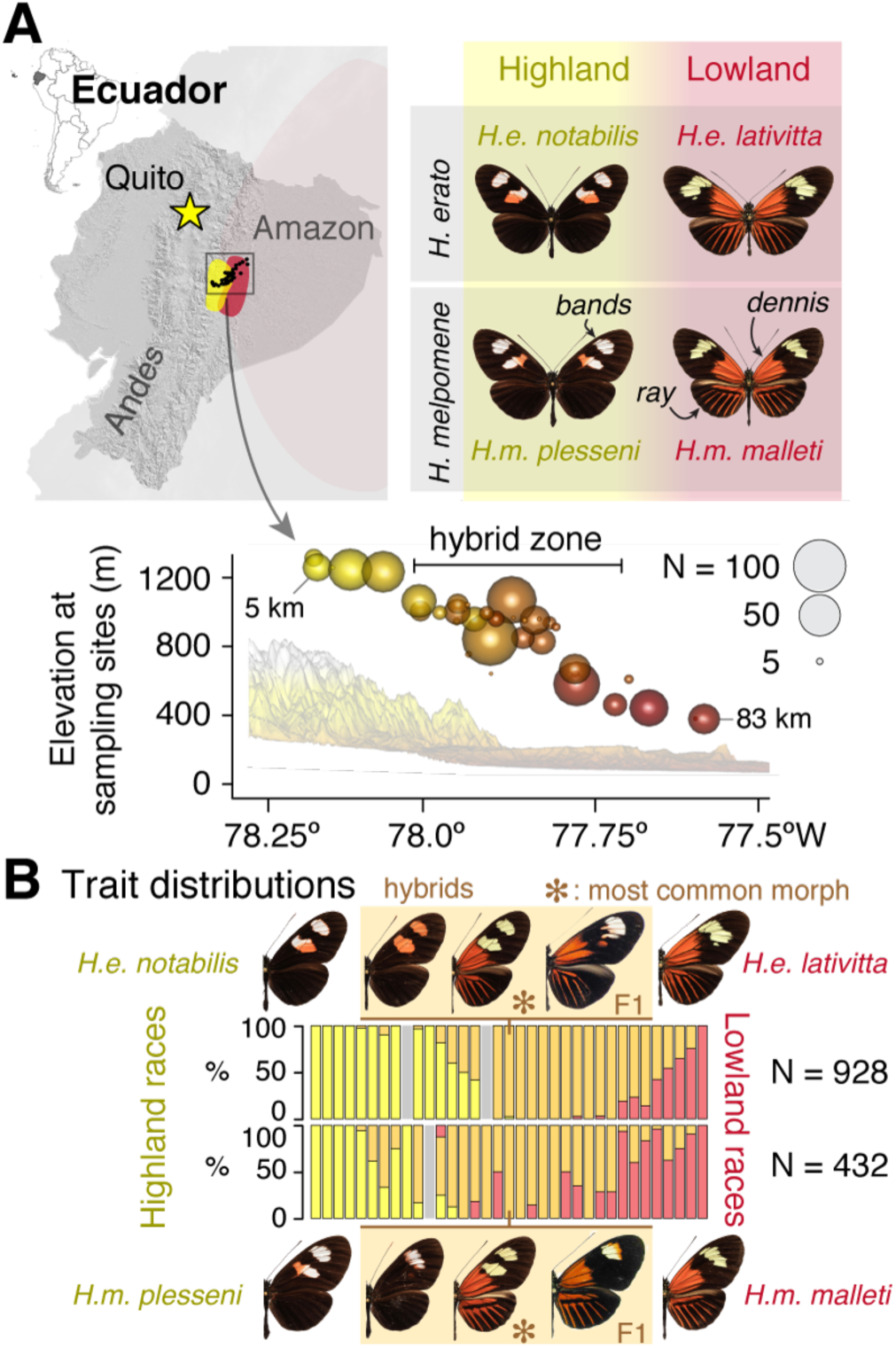
Parallel hybrid zones in a pair of Müllerian co-mimicking *Heliconius* butterflies. (**A**) In eastern Ecuador, butterflies of the species *H. erato* and *H. melpomene* occur in the transition zone between the Andes (up to 1307 m elevation, “Highland”) and the Amazon basin (376 m, “Lowland”) as distinctive races with major wing color pattern differences (labelled as “bands”, “*dennis*” and “*ray*”). *Heliconius* butterflies are unpalatable and share warning wing patterns (Müllerian co-mimicry) (*17*). We sampled a total of 1,360 butterflies of both species along an 83 km transect consisting of 35 sampling sites across the double hybrid zones (km 19 – 59; symbols scaled to sample size and colors indicate elevation) and 12 additional off-transect sites (Table S5). (**B**) Proportions of butterflies displaying the highland double-band phenotype (*H. erato notabilis* and *H. melpomene plesseni*: yellow) and lowland *dennis*-*ray* patterns (*H. erato lativitta* and *H. melpomene malleti*: red) as well as hybrid patterns (F1 and beyond: orange; *, most common morph; fig. S3B; grey: sites with no specimen in one species) at sampling sites along the transect.

Here we focus on two distantly related *Heliconius* species, *H. erato* and *H. melpomene* (diverged 12 million years ago) (*26*), which feature many distinct color patterns and mimic each other (and other species) whenever they overlap (16). In the Pastaza valley of eastern Ecuador, a highland race of each species meets and forms a hybrid zone with a distinct lowland race (Fig. 2A; fig. S3A; (*27, 28*)). The hybrid zones range from around 1,300 m to 400 m from the Andean mountains into the Amazon basin (highland race: *H. erato notabilis* and *H. melpomene plesseni*; and lowland race *H. erato lativitta* and *H. melpomene malleti*; Fig. 2). To survey the hybrid zone, we collected 975 *H. erato* and 394 *H. melpomene* butterflies (928 and 343 at 35 transect sites; Fig. 2; fig. S3A; Tables S6–7) and scored their color traits as informed by controlled laboratory crosses (fig. S3C–F) (*27*). Figure 2b shows that hybrid butterflies (both F1 and beyond) are observed in all but five highland and one lowland sites, with the core transition zone between 1,000–900 m of elevation (km 36 – 45 along the transect).

### Divergence, selection and trait mapping

Using haplotagging, we sequenced 484 *H. erato* and 187 *H. melpomene* butterflies from the transect in 96-plex batches to a median read coverage of 1.29× for *H. erato*; and 2.72× for *H. melpomene* (samples both individually *and* molecularly barcoded; see Material and Methods; Tables S3, S5–8). Following phasing and imputation, we retained a set of 49.2 M single nucleotide polymorphisms (SNPs) in *H. erato* and 26.3 M SNPs in *H. melpomene*, most of which were polymorphic throughout the zone (*H. erato*: 69.4%; *H. melpomene*: 81.1%), consistent with high gene flow. By contrast, only 232 SNPs were completely fixed for opposite alleles between *H. e. notabilis* and *H. e. lativitta*; and none between *H. m. plesseni* and *H. m. malleti.* Sequence diversity was high (131 and 97 SNPs / kbp in *H. erato* and *H. melpomene*, respectively), which helped to produce long average phased block sizes of 3.6 and 3.3 Mbp in *H. erato* and *H. melpomene* respectively (maximum: 20.7 Mbp, effectively spanning a whole chromosome; fig. S4). This dataset represents a qualitative jump in quality and resolution over the state-of-the-art in a natural population study.

Across the genome, there was little background genomic differentiation between highland and lowland races in both *H. erato* and *H. melpomene* (mean genetic distance F_ST_ in *H.* erato: 0.0261 and in *H. melpomene*: 0.0189; Fig. 3). This is consistent with free introgression of neutral and globally adaptive variants in hybrid zones (*29, 30*).

**Fig. 3.**
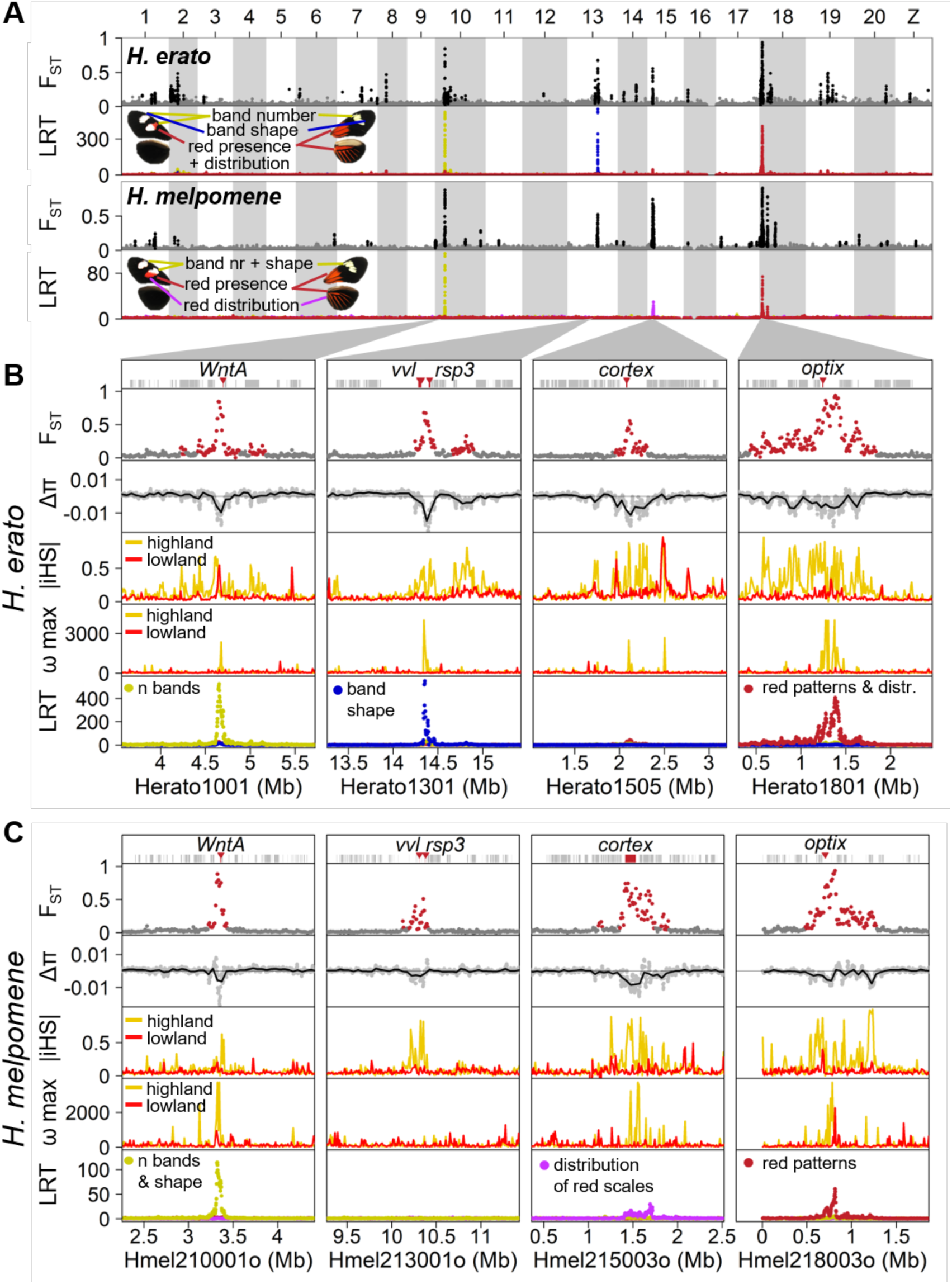
Highly parallel patterns of differentiation at genomic regions underlying wing color patterns. (**A**) Major peaks of differentiation are shared across *H. erato* and *H. melpomene* (as indicated by F_ST_; *H. melpomene* data is plotted at its homologous *H. erato* coordinates). F_ST_ values of windows assigned to the high differentiation state by the HMM analysis, are shown in black, others in grey. The three most strongly differentiated regions in each pair of subspecies all show strong association with color pattern differences (LRT: likelihood ratio test). (**B** and **C**) Detailed view at the four loci with strongest differentiation in *H. erato* (**B**) and *H. melpomene* (**C**). At all four major loci, the races also differ in nucleotide diversity (π; Δπ=π_*highland*_ – π_*lowland*_), whereby the highland races (*H. e. notabilis* and *H. m. plesseni*) consistently show greater reduction in diversity than the lowland races (*H. e. lativitta* and *H. m. malleti*), indicative of strongest selection in the highland races in both species. Compared to the Δπ values of all genomic 50 kb windows, the four major loci are among the most negative 1% in both species (fig. S5). Stronger selection among highland races than lowland races is also supported by haplotype-based selection statistics such as absolute normalized iHS (integrated haplotype score) and the ω-statistic. Three of the four major loci in each species are associated with major color patterns and all fall into the vicinity of the genes *WntA* (forewing band number), *Ro* (likely *vvl* and/or *rsp3 (24)*), *cortex* (distribution of red scales) and *optix* (presence of red either as forewing patch and hindwing bar and rays (*dennis-ray*) or in forewing band) in *H. erato* (**C**) and in *H. melpomene* (**C**). (for details see fig. S9 and Data S4–5).

Against this backdrop, peaks in genomic differentiation stand out in stark contrast in each species. Using a Hidden Markov Model (HMM), we identified 24 and 52 regions of high differentiation in *H. melpomene* and *H. erato*, respectively (Fig 3; Supplementary Methods). The strongest divergence peaks are found at four major color loci, namely *WntA* (Chromosome 10) (*22*), *Ro* (Chr13), *cortex* (Chr15) (*31, 32*) and *optix* (Chr18) (16, *21, 33*)(Fig. 3B–C). The improved resolution from haplotagging reveals novel loci and greater parallelism than previously described (*28*), with F_ST_ peaks at *cortex* in *H. erato* and *vvl* in *H. melpomene* (the putative homologue of *Ro* in *H. erato*) being the most surprising, because they have not previously been hypothesized to play a role in phenotypic divergence in these races (see fig. S6 for a direct comparison; (*28*)). The windows with the highest F_ST_ values are all located in these four major color loci and are highly correlated between the two species highlighting the fine-scale parallelism at these loci (fig. S7).

All four loci show reduced nucleotide diversity and elevated linkage disequilibrium (LD), characteristic signatures of selective sweeps (captured by the *π, ω*, and iHS statistics; Tables S9–10, Data S4 & S5 (*34, 35*)). Here, the LR data allows us to track the breakdown of haplotypes across the hybrid zone (fig. S8) and greatly increase the power of the haplotype-based *ω*-statistic (especially compared to haplotypes reconstructed from short-read only data, fig. S6C). The resolution in these tests is high enough to separate the *dennis* and *ray cis-*regulatory regions from the target gene *optix* in *H. erato* (fig. S6C). The data can reveal unsuspected molecular details, as evidenced by the detection of rare *H. melpomene* recombinants at the tightly linked *band, dennis* and *ray* elements at *optix*. These recombinants show that the presence of red scales in the forewing bands is controlled solely by *dennis* but not *band*, as in other *H. melpomene* races (fig. S9A; (*55, 85*)). In addition, one particularly informative recombinant helps to refine the *ray* element from 14.1 kbp to 8.0 kbp (fig. S9B). Together, our data underscore the precision and power of population haplotyping.

### Chromosome inversions and other structural rearrangements

In local adaptation and speciation, chromosome rearrangements, and inversions in particular, are thought to play a major role in holding together adaptive variants (*36*-*39*). However, they are hard to detect using short-read techniques. By contrast, longer LR molecules that span rearranged junctions can systematically reveal insertions, deletions, inversions and additional chromosome rearrangements. We therefore analyzed beadTag sharing across adjacent 10 kbp windows to detect differences between the physical molecules and the reference assembly (Fig. 4; see Methods for details). We detected 685 and 415 indels; and 14 and 19 major inversion/translocation events in *H. erato* and *H. melpomene*, respectively (Data S2 & S3).

**Fig. 4.**
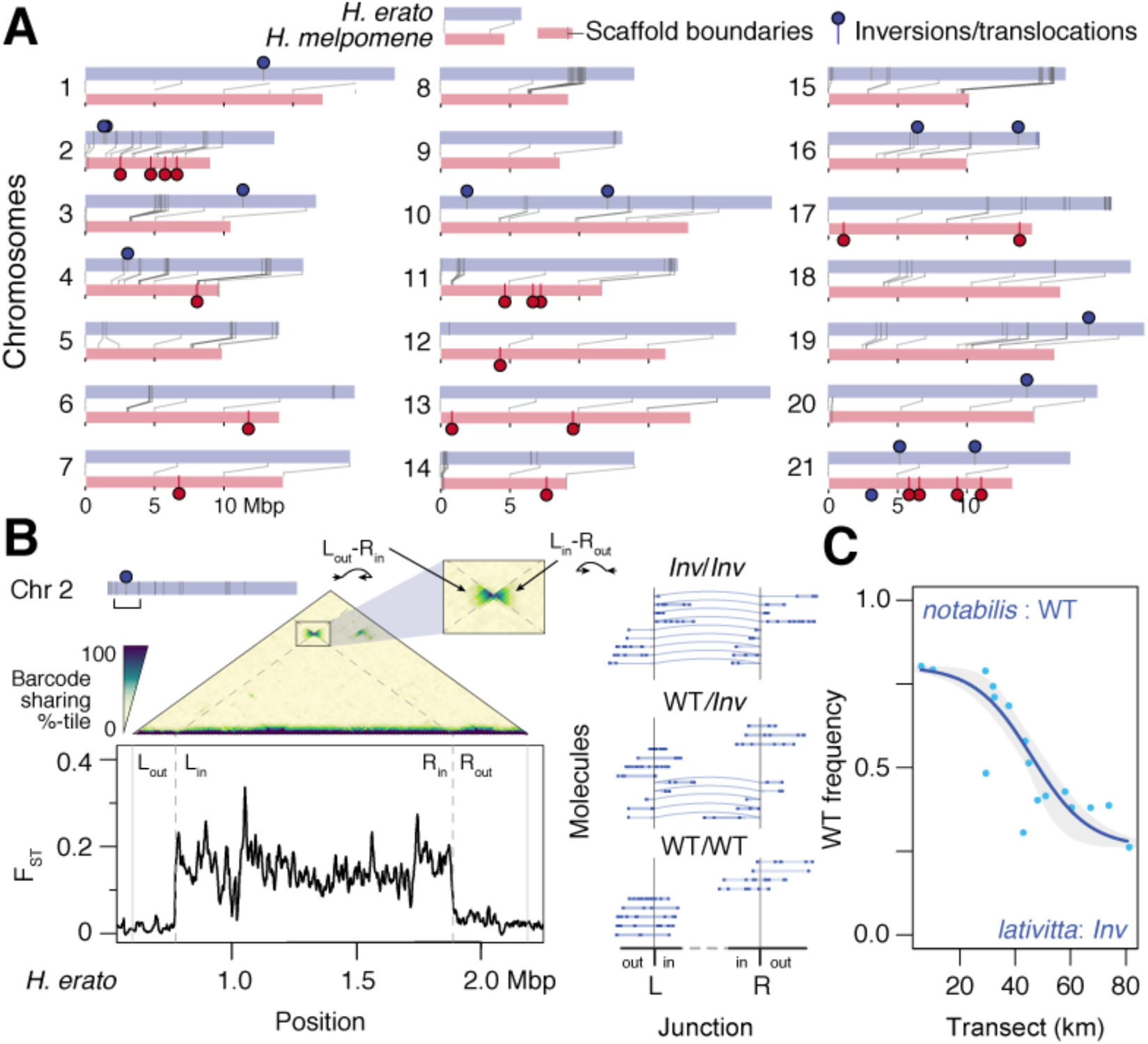
Distinct structural rearrangements across the parallel hybrid zones. (**A**) Locations of major structural rearrangements in the two *Heliconius* hybrid zones. Chromosome homologues are shown in pairs, with lines connecting syntenic positions between *H. erato* (grey) and *H. mel.* (red; lines; dark grey bars mark scaffold boundaries; circles mark major inversions or translocations). In contrast to the parallelism at divergent peaks shown in Fig. 3, major structural rearrangements tend to be unique for each species. (**B**) Detection of a major inversion on *H. erato* Chr2. The average linked-read molecule spans multiple 10 kbp windows. Thus, the extent of beadTag sharing across windows (10 kbp here) can reveal discrepancies between the physical molecules and the reference assembly as well as across populations. The triangular matrix shows a heatmap of barcode sharing (color indicates genome-wide percentile) juxtaposed against genetic distance (F_ST_) across the pure *notabilis* and *lativitta* races. Inversions appear as a “bow-tie” shaped pattern across the inverted junction boundaries (L, left boundary of the inversion; R, right boundary of the inversion; out/in, outside or inside of the inversion; Left_in_/Right_out_ and Left_out_/Right_in_, zoomed inset). This inversion coincides with a plateau of elevated genetic distance across the *notabilis* and *lativitta* races. Dotted lines mark the inferred inversion boundaries at Herato0204:172503– 1290057. Molecules from three individuals representing the three inversion collinear vs. hetero-karyotypes are shown (inferred inversion indicated with curved arrows). (**C**) The Chr2 inversion shows a clinal distribution across the *notabilis*–*lativitta* hybrid zone (frequency of WT karyotype: blue dots; fitted cline: blue line; confidence interval: grey envelope).

Although structural rearrangements occasionally overlap divergent peaks or signatures of selection in single high- or lowland populations in either species, generally speaking they differ only very little between highland and lowland populations. There is also no sign of parallel rearrangements between *H. erato* and *H. melpomene* (Fig. 4A). However, a specific rearrangement on *H. erato* Chromosome 2 (Chr2) stands out. Here, among all *H. erato* samples we observed unusually high beadTag sharing between windows 1.1 Mbp apart, in a manner indicative of an inversion (Fig. 4B; inferred junctions at “Left”: Herato0204:172503 and “Right” Herato0204:1290057). This elevated signal is especially strong among lowland *H. e. lativitta* butterflies and is distinct from previously reported inversions on chromosome 2 (*24, 40, 41*), suggesting that the inversion may have originated in *H. e. lativitta* or its close relatives. The two junctions bracket a region of elevated F_ST_ (Fig. 4B), suggesting reduced gene flow in this genomic region between the highland *H. e. notabilis* and lowland *H. e. lativitta* races. Using LR data, we have the opportunity to directly detect molecules that span the inverted Left and Right junctions, i.e., Left_out_–Right_in_ and Left_in_–Right_out_ (Fig. 4B, left, “bow-tie” pattern). This we find in 113 *Inv/Inv* individuals homozygous for the inverted allele (as well as 163 WT/*Inv* and 152 WT/WT individuals; Fig. 4B, middle panel), and show that the inversion is segregating across the *H. erato* hybrid zone.

### Haplotype frequencies across the hybrid zone

In both species, migration and gene flow between the high- and lowland forms generated clines across the genome, i.e. gradient in gene frequencies along the transect (fig. S10). For example, at the Chr2 inversion in *H. erato*, the wild type orientation decreases from 80.2% in the highland *notabilis* race to 26.3% in the lowland *lativitta* one (or 73.7% inverted; estimated centre of zone: 46.6 ± 3.2 km; width: 53.44 ± 23.7 km; Fig. 4C). This inversion contains 50 genes, and a spike in F_ST_ and *ω* within 25 kbp of a putative homeodomain transcription factor and a previous reported heat shock protein (*28*). In cline analysis, the steepness indicates selection: neutral loci introgress freely and produce wide and shallow clines, whereas strongly selected loci remain distinct between the races and produce sharp and narrow clines. Accordingly, the major color loci are among the narrowest clines in the genome (fig. S11). Plotting of both phenotypic and haplotype clines at *optix* (Chr18) and *WntA* (Chr10) in the two co-mimetic species shows a striking pattern: in each species, the *WntA* cline centre is shifted east towards the lowlands (at a large drop in elevation between km 45 – 50) relative to the *optix* cline, or indeed much of the genome (Fig. 5A; 15.28 and 20.87 km in *H. erato* and *H. melpomene* respectively; fig. S11). However, at these two color loci both the positions and widths of the clines closely mirror each other between *H. erato* and *H. melpomene* (*optix*, centres: 31.9 vs. 28.9 km; widths: 15.7 vs. 15.2 km; *WntA*, centres: 47.1 vs. 49.8 km; widths: 19.0 vs. 24.8 km; Fig. 5A & B; Table S11). Interestingly, the minor color loci (*Ro* and *cortex*) track with a different major color locus in each species: *Ro* (Chr13) tracks with *WntA* in *H. erato*; and *cortex* (Chr15) tracks with *optix* in *H. melpomene* (figs. S11–12). Both show broader cline widths, likely due to dominance and reduced phenotypic effects and thus weaker selection (Table S11; (*27*)). In fact, genetic crosses suggest that the displacement of the clines at these minor loci may reflect the underlying genetics in refining the match across the co-mimics: *Ro* acts as a modifier for *WntA* in shaping the forewing band in *H. erato* butterflies and its cline is shifted relative to other loci only in *H. erato* but not in *H. melpomene.* Likewise, *cortex* modifies *optix* to generate the fully *dennis*-*ray* phenotype only in *H. melpomene* and its cline shows a closer match with *optix* there than in *H. erato* (fig. S3) (*27*).

**Fig. 5.**
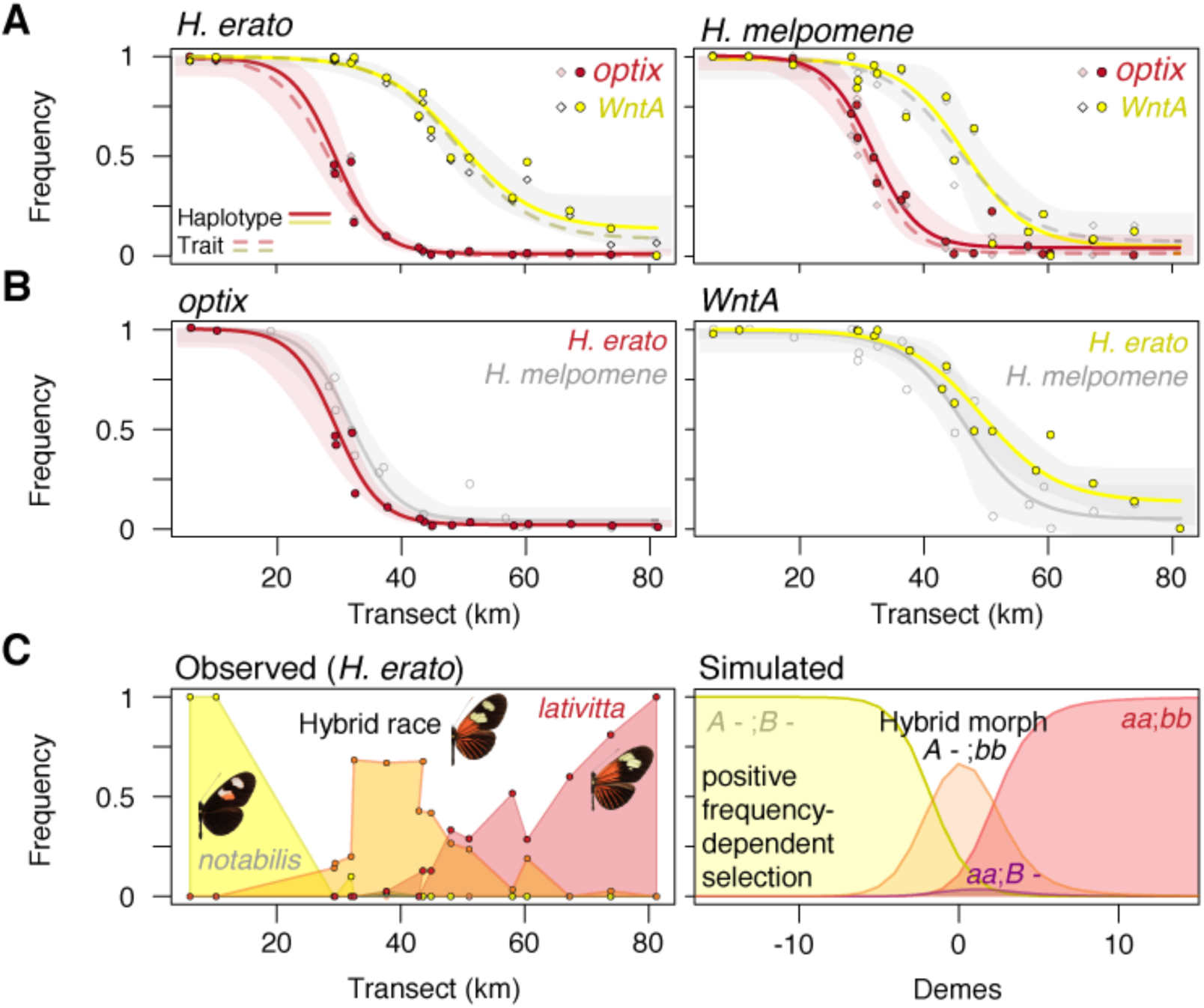
Müllerian co-mimicry and the emergence of a hybrid race due to mirrored cline displacement of color traits. (**A**) Major color traits segregating across the Ecuadorian hybrid zones show a clinal distribution of haplotype frequencies along the transect in both *H. erato* (left) and *H. melpomene* (right). There is strong agreement in cline fits between haplotype frequencies (filled circles; cline: colored lines with 95% confidence envelope) and phenotype frequencies (diamonds and dashed lines). *Optix* (red) controls the red color pattern (see Fig. 3 and fig. S3E– F) and shows a steeper and west-shifted cline compared to *WntA* (yellow), which controls the number of forewing bands (Fig. 3). (**B**) Clines are mirrored at both *optix* (left) and *WntA* (right) loci between *H. erato* (filled circles and colored lines) and its *H. melpomene* co-mimic (empty circles and grey lines). **c**, Emergence of a novel hybrid morph in the middle of the hybrid zone. Due to the displaced clines, hybrid *H. erato* butterflies (left panel; orange symbols and lines; middle wings) can display the highland *notabilis* double-band (left; yellow) along with the lowland *lativitta dennis* and *rays* (right; red). This hybrid morph carries homozygous *WntA*^H/H^ and *optix*^L/L^ genotypes and is therefore true-breeding. Simulation results show the frequencies of the four morphs, assuming complete dominance at two loci. Morph *i* has fitness 1 + *s*_*i*_ (*P*_*i*_ *-Q*_*I*_), which increases linearly with its own frequency, *P*_*i*_. Even when clines at the two loci start fully coincident, they can shift apart and produce displaced clines over time (here, generation 1000), if there is a fitness advantage to one of the hybrid genotypes, here *s*_*A -; bb*_ = 0.25, and the rest having *s*_*i*_ = 0.1.

The finding of displaced clines at the major color loci makes this Ecuadorean hybrid zone unique among *Heliconius* hybrid zones. In all other *Heliconius* hybrid zones both *H. erato* or *H. melpomene* clines across the genome coincide regardless of mimicry (e.g., Peru (*42, 43*) and Panama and Colombia (*44*)). This may be because whenever clines overlap, they tend to be pulled together into coincidence due to increased LD (*45*).

To estimate the strength of selection, we analyzed the shape and correlation between clines at unlinked loci, and contrasted these results against previous estimates from the coincident zone in Peru (*19*). Whenever clines overlap, linkage disequilibrium (LD) is generated through the admixture of distinct populations, even between unlinked loci; the correlation between unlinked alleles is expected to have a maximum approximately equal to selection [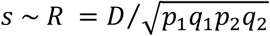; for linear frequency-dependent selection (FDS) at two loci with complete dominance, *R* = 0.73*s*; under linear FDS, selection favors the most common morph, as in Müllerian mimicry]. Previous studies in Peru gave *R* ∼ 0.35 in *H. erato*, and *R* ∼ 0.5 in *H. melpomene (19, 43)*. Here, we do not expect LD between shifted clines but neither do we see any significant association between coincident clines (Tables S14 and S15). The maximum *R* consistent with our data is ∼0.054 in *H. erato*, and ∼0.154 in *H. melpomene* (Table S15). Thus, we estimate selection to be substantially weaker (< 4% in *H. erato* and 11% in *H. melpomene*).

We can set upper limits to the dispersal rates (σ) that would be consistent with observed cline widths (*w*) and weak LD; assuming linear FDS with complete dominance 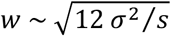 (*42*) so that 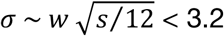 km in *H. erato* and 6.2 km in *H. melpomene* (Table S15). These upper limits to dispersal are slightly larger than, and so consistent with, previous estimates in Peru (*42*). Thus, our data suggest that the clines here are maintained by substantially weaker selection, but dispersal rates may be similar. This weaker selection may in part be because the coincident clines involve loci with minor effects.

### Emergence of a novel hybrid morph

One consequence of the displaced *WntA* and *optix* clines is that a novel, hybrid morph combining the highland double forewing band with the lowland *dennis*-*ray* pattern (*WntA*^H/H^;*optix*^L/L^) has become common in the middle of both *Heliconius* hybrid zones. Indeed, from 33 km – 45 km along the transect, this novel *WntA*^H/H^;*optix*^L/L^ genotype is the most common morph in both species (Fig. 2B & 5C, left; Tables S6–7). We used deterministic two-locus simulations to test whether positive FDS could maintain a novel hybrid morph. Our data with the shifted cline centres are largely consistent with dominance of the lowland over the highland allele at *WntA* and *optix*, and fit with the higher similarity of F1 individuals to lowland rather than highland individuals. This implies introgression of highland alleles into the lowland populations with four possible morphs (*WntA*^*H/H*^;*optix*^*H/H*^, *WntA*^*H/H*^;*optix*^*L/-*^, *WntA*^*L/-*^;*optix*^*H/H*^, *WntA*^*L/-*^;*optix*^*L/-*^; Table S16). Over time, the cline centres are expected to shift west towards the highlands (*46*). All else being equal for the four morphs, clines that are well-displaced to start with can remain separate, because each cline moves at the same speed. But if they overlap, either initially or because the leading cline stops moving due to other factors, LD pulls them together (fig.S13).

The above scenario will produce a hybrid morph that can persist, perhaps indefinitely; but its distribution will shift west over time, given suitable habitats. Since the distribution of the hybrid morph appear to be stable (*47*), there must be additional factors that cause clines (including ones that at first coincide) to shift and remain apart. These could include genetic incompatibilities or a selective advantage of one hybrid morph over the other (*48, 49*), but we favor a model in which the hybrid morph is favored or experiences stronger FDS, perhaps due to a more memorable phenotype for predators, which can maintain stable shifted clines as observed in the empirical data (fig. S13, bottom). Unlike hybrid morphs with heterozygous genotypes, this novel *WntA*^H/H^;*optix*^L/L^ hybrid morph breeds true and has risen to appreciable frequencies, perhaps representing establishment of a novel hybrid race.

## Discussion and Conclusions

The discovery and characterization of natural variation in the genome is a key first step in genetics and evolution. Such information can help us understand the genetic basis of trait variation and speciation. However, until now it has not been easy to capture this variation as haplotypes in large population samples. Haplotagging solves this problem by generating linked-read data from hundreds of samples efficiently and affordably. These data are far richer in information and permit the simultaneous characterization of both nucleotide and structural variation.

More broadly, this work highlights the advantage of combining broad population sampling with linkage information in large-scale LR data. Together, they allow efficient and accurate genome-wide haplotyping, as opposed to genotyping. We hope that this work will spur development of improved algorithms and experimental designs, such that future researchers may be able to perform (meta)genome assembly, phasing, imputation and mapping in a single experiment. We anticipate that haplotagging or similar approaches (and eventually long-read sequencing) may help drive the next phase of discoveries in model and non-model organisms alike.

We have used these data to demonstrate the early stages in establishment of a novel hybrid race, through the parallel displacement of clines in two species. In addition, we have also discovered >300 novel candidates under local or divergent selection, which opens up additional dimensions beyond wing color patterns to investigate this double hybrid zone. Somewhat to our surprise, our survey for structural rearrangements such as inversions were not consistently associated with population differentiation in either species, suggesting that they do not play an early role in mediating divergence in the face of gene flow, despite widespread support in the literature (*36, 38, 50, 51*), (reviewed in (*52*) but see (*40*)). More broadly, our results highlight further the evolutionary potential of hybridisation in local adaptation and the early stages of speciation (*53*-*55*).

## Material and Methods

See Material and Methods in Supplementary Materials

## Acknowledgements

We thank Felicity Jones for input into experimental design, helpful discussion and improving the manuscript. We thank the Rolian, Jiggins, Chan and Jones Labs members for support, insightful scientific discussion and improving the manuscript. We thank the Rolian lab members, the Animal Resource Centre staff at the University of Calgary, and Caroline Schmid and Ann-Katrin Geysel at the Friedrich Miescher Laboratory for animal husbandry. We thank Christa Lanz, Rebecca Schwab and Ilja Bezrukov for assistance with high-throughput sequencing and associated data processing; Andre Noll and the MPI Tübingen IT team for computational support. We thank Ben Haller and Richard Durbin for helpful discussions. We thank David M. Kingsley for thoughtful input that has greatly improved our manuscript. J.I.M. is supported by a Research Fellowship from St. John’s College, Cambridge. A.D. was supported by a European Research Council Consolidator Grant (No. 617279 “EvolRecombAdapt”, P/I Felicity Jones). C.R. is supported by Discovery Grant #4181932 from the Natural Sciences and Engineering Research Council of Canada and by the Faculty of Veterinary Medicine at the University of Calgary. C.D.J. is supported by a BBSRC grant BB/R007500 and a European Research Council Advanced Grant (No. 339873 “SpeciationGenetics”). M.K. and Y.F.C. are supported by the Max Planck Society and a European Research Council Starting Grant (No. 639096 “HybridMiX”).

## Funding statement

The authors declare competing financial interests in the form of patent and employment by the Max Planck Society. The European Research Council provides funding for the research but no other competing interests.

## Notes

### Summary of Updates

Reformatted, and fig. S9 updated for improved presentation of interval mapping.

