## Supplementary Materials for "Haplotype tagging reveals parallel formation of hybrid races in two butterfly species"

**Supplementary Notes**
**Supplementary Methods**
**Supplementary Figures**
**Supplementary Tables**

Table S1. Linked-read sequencing technique comparisons
Table S2. Example sequencing library preparation costs
Table S3. Sequencing Summary
Table S4. Phasing performance in single individuals and comparisons with other platforms
Table S5. Sampling locations
Table S6. *H. erato* individuals collected by site and genotype.
Table S7. *H. melpomene* individuals collected by site and genotype.
Table S8. Validation results for STITCH at known color loci
Table S9. Top loci based on ω, sweeD and iHS in *H. erato*
Table S10. Top loci based on ω, sweeD and iHS in *H. melomene*
Table S11. Cline analysis at five different loci
Table S12. Loci with narrow clines in *H. erato*
Table S13. Loci with narrow clines in *H. melpomene*
Table S14. Maximum likelihood estimates for heterozygote deficit, $\hat{F}_{IS}$
Table S15. Maximum likelihood estimates for the correlation between loci ( $R = D/\sqrt{p_1q_1p_2q_2}$ ) , together with the difference in log(likelihood) relative to $R = 0$
Table S16. Testing for asymmetry of single-locus clines

**Supplementary Data**
Data S1. Oligonucleotide used for assembling haplotagging beads
Data S2. Structural rearrangements
Data S3. Insertions and deletions
Data S4. Detailed plots of $F_{ST}$, selection statistics and association by chromosome
Data S5. Locations of highly differentiated $F_{ST}$ regions and signatures of selection

**Supplementary References**

**Supplementary Notes**

Segmental barcode design to maximize diversity within strict length limitations
Due to the current limits of a maximum of 25 indexing cycles in the sequencing recipe design and reagent amounts in standard Illumina sequencing flow cell kits, the configuration supporting the highest barcode diversity would be achieved by partitioning a total of 25 indexing cycles into segments, each of which of 4nt to up to 9nt long, as shown below.

| Length | Single segment (with error correction) |
|---|---|
| 3nt | 4 |
| 4nt | 12 |
| 5nt | 48 |
| 6nt | 85* |
| 7nt | 278 |
| 8nt | 727 |
| 9nt | 2620 |

* current chosen option

For example, a 13nt i5 index read can be split into segments of 6nt + 7nt, yielding 23,630 combinations; alternatively, it can be 5nt + 8nt, yielding 34,896 combinations. Combined with the costs of synthesizing the required oligonucleotide duplexes, then the slightly lower diversity in a 6nt + 7nt barcode combination becomes favorable, because it only requires a total of 363 unique sets of duplexes, whereas 5nt + 8nt would require nearly double the number of duplexes (775). This latter factor also has downstream effect on the amount of reagents used for the assembly and synthesis of beads.

Therefore, the practical solution for the lowest cost configuration would be to synthesize 4 segments of 6nt or 7nt barcodes, such that together they make up 12 and 13 cycles of i5 and i7 index reads.

Demultiplexing
The characteristics of robust barcode design at 6nt are shown below:

| | 6 nt barcodes (full set of 96) | | | 6 nt barcodes (first 84) | | |
|---|---|---|---|---|---|---|
| Description | Hamming | SeqLev | Levenshtein | Hamming | SeqLev | Levenshtein |
| Mean distance | 4.54 | 3.19 | 4.08 | 4.54 | 3.19 | 4.08 |
| Median distance | 5 | 3 | 4 | 5 | 3 | 4 |
| Minimum distance | 2 | 1 | 2 | 3 | 1 | 2 |
| Maximum distance | 6 | 6 | 6 | 6 | 6 | 6 |
| Guaranteed error correction | 0 | 0 | 0 | 1 | 0 | 0 |
| Guaranteed error detection | 1 | 0 | 1 | 2 | 0 | 1 |

The table above shows the complexity statistics for a set of 84 barcodes and the effect of adding 12 additional barcodes to make it up to 96, such that the entire split-and-pool assembly reaction can be performed on standard 96-well plate formats, with minimal effects on the possibility to detect or where possible, correct errors. With 96 barcodes per segment and a total of 4 segments or 24nt plus 2 – 4nt for overhangs, a set of barcodes with up to approximately 85 million combinations can be encoded among the beadTags.

Given the strict limit on the combined length of the i5 and i7 index reads (25nt) under standard running conditions, the feasibility of efficient ligation with an 1nt overhang was evaluated. In addition, to avoid having the higher costs associated with synthesizing multiple attaching biotinylated primer in order to vary the overhang for ligation, 5' overhangs on the short, complementary strand (instead of the more stable and common 3' overhang) were designed. We found that we were able to use the Blunt/TA ligase kit from New England Biolabs to achieve robust ligation, despite having only 1 bp 5' overhang. With this result the possible combinations of barcodes were extended to over 85 million with four sets of duplexes of 96 each.

Differences from other LR platforms

The CPT-seq method, as disclosed by Amini et al. 2014, involves two or more separate step (*6*). The first step of CPT-seq introduces a first set of barcodes through Tn5 transposition, followed by splitting of the bulk samples into separate pools for subsequent amplification or ligation of a second set of barcodes. Having these two steps is cumbersome and the required additional handling involving PCR amplification increases the chance of introducing undesired nucleic acid exchanges that can decrease sequencing accuracy. Most crucially, the method does not lend itself to high-throughput, highly multiplexed applications, which would be necessary if CPT-seq were to be performed on a large number of samples simultaneously.

CPTv2-seq as described by Zhang et al. 2017 uses a slightly different strategy involving beads (*8*). Their single-tube variant involves tagmenting target DNA with pre-assembled transposome complexes and hybridization thereof to beads comprising bead-specific oligonucleotides comprising two barcode sequences separated by a splint 1 and splint 2 sequence. While the method avoids an amplification step, the hybridization of the beads and the transposome complexes adds additional complexity to the protocol that may introduce errors. They also require an aldehyde modification to the primers. In their study, this was carried out using custom processes.

Another major limitation of CPTv2-seq of Zhang et al. is that this method only employs 147,456 different barcode combinations. As described in more detail below, a set of only 147,456 unique barcodes falls far short of the number required to avoid barcode re-use (a form of "barcode collisions") due to the high number of DNA molecules present in a typical reaction volume.

Importantly, CPTv2-seq has the disadvantage of producing sequencing libraries that are not compatible with standard Illumina Nextera sequencing reagents and thus require customized sequencing primers and run protocols for both sequencing the barcodes and the target sequence. As it is presently configured, it precludes the ability to run samples generated through CPTv2-seq together with Nextera or TruSeq protocols in the same Illumina flow cell. This is a major drawback that greatly limits the reach of CPTv2-seq. Due to the design of the beads used in CPTv2-seq, it is highly inconvenient, if not impossible, to operate the libraries generated with this method on an Illumina HiSeq or NovaSeq sequencing instrument with standard sequencing primers and settings. Instead, whole sequencing runs featuring exclusively CPTv2-seq libraries may have to be scheduled to enable access to the CPTv2-seq technology. This significantly reduces the multiplexing capability and leads to additional costs and unnecessary delays.

Yet another method, single-tube long fragment reads (stLFR) described in

Wang et al., 2019 (*9*), also uses a hybridization step following tagmentation in solution. This configuration bears many similarities to CPTv2-seq, and is optimized for BGI's proprietary sequencers. Although they provide an option for sequencing on Illumina's platform, their barcode design also involves long splint segments, extended index read-lengths, as well as custom primers.

Phasing in single individuals and large populations

We evaluated phasing performance by haplotagging DNA from an additional recombinant mouse and the human cell line GM12878 that is widely used in genome assembly and phasing comparisons (*56*). Consistently, we achieved robust diploid phasing success (98.59% to 99.91% heterozygous single nucleotide polymorphisms or SNPs phased, with molecule $N_{50}$ ranging from 40.87 to 63.47 kbp; the largest molecule spans 573 kbp). Phased block size was 1.08 Mbp in GM12878 and around 15 Mb in mice (maximum: 61.46 Mbp). Phasing errors, both as short switches (single mismatch positions, 0.95% in human and <0.18% in mice) and long switches, e.g., 0001111, were minimal (<0.04% in all cases, Table S4).

These performances are comparable to benchmarks set previously by CPTseqv2 and 10X Chromium (Table S4). For example, haplotagging produces on average only one to two molecules per barcode due to its greater barcode diversity, well below the average of 6 from CPTseqv2 and 10X Genomics's Chromium platform (*7*, *8*). It should be noted, however, that direct comparisons are not always possible due to underlying technical differences, e.g., the power to detect long switch errors increases with longer molecules and greater number of reads per molecule.

### Supplementary Methods

### Material and Methods

Animal Care and Use
All experimental procedures described in this study have been approved by the applicable University institutional ethics committee for animal welfare at the University of Calgary (HSACC Protocols M08146 and AC13-0077); the local competent authority: Regierungspräsidium Tübingen, Germany, permit and notice numbers 35/9185.46-5 and 35/9185.82-5. The study and collection of *Heliconius* butterflies has been approved by the Ecuadorian government with permits to C.D.J and P.A.S.: Collection permits: 033-10-IC-FAU/FLO-DPN/MA, research permits: 0007-IC-FAU/FLO-DPPZ/MA and 013-09 IC-FAU-DNB/MA. Export permits: 01-2011-FAU-DPAP-MA and 002-EXP-CIEN-FAU-DPN/MA.

Data and code availability
Analysis codes are available at the following repositories: https://github.com/joanam; https://github.com/evolgenomics; and https://github.com/rwdavies. Sequence data have been deposited at the NCBI Short Read Archive under accession numbers: [PENDING]. Phenotype data are available at Dryad.

Reference genome assemblies
All co-ordinates in the human genome refer to `GRCh38`, specifically the `hg38` assembly as part of the Resource Bundle hosted by the Broad Institute (ftp://gsapubftp-anonymous@ftp.broadinstitute.org/bundle/hg38/). Mouse genome co-ordinates refer to *Mus musculus* reference `mm10`, which is derived from `GRCm38`. Butterfly genomic data are placed against the *Heliconius erato demophoon* version 1 assembly (`helera1_demo`; http://download.lepbase.org/v4/sequence/Heliconius_erato_demophoon_v1_-_scaffolds.fa.gz) and the *Heliconius melpomene melpomene* version 2.5 assembly (`Hmel2.5`; (http://download.lepbase.org/v4/sequence/Heliconius_melpomene_melpomene_Hmel2.5.scaffolds.fa.gz) according to each individual.

Tn5 transposase
Sequencing libraries for high-throughput sequencing were generated using Tn5 transposase expressed in-house as previously described (*57*). Briefly the bacterial expression plasmid pTXBX1-Tn5 (Addgene plasmid #60240) containing the hyperactive Tn5 transposase (carrying the E54K, L372P mutations) fused to an intein chitin-binding domain was transformed into the C3013 competent cells (New England BioLabs, Frankfurt am Main, Germany). Expression was induced under addition of isopropyl β-D-1-thiogalactopyranoside (IPTG) and cells were lysed using an Emulsiflex c3 (Avestin, Mannheim, Germany). The lysate was applied to a chitin resin column (New England BioLabs). The Tn5 transposase domain was cleaved and eluted using 1,4-dithiothreitol (DTT, Sigma Aldrich, Taufkirchen, Germany). Concentration of the eluted protein and DTT removal was achieved through a concentration column with a cut-off of 10 kilodaltons (Amicon Ultra-15, 10kDA, Merck-Millipore, Darmstadt, Germany).

Oligonucleotide design
Custom oligonucleotides were synthesized by Integrated DNA Technologies (Leuven, Belgium) at ready-to-use 10 μM concentration in a 96-well plate format. A list of the oligonucleotides can be found in Data S1.

Haplotagging bead assembly
Haplotagging beads are individually barcoded "Dynabeads M-280 Streptavidin magnetic beads" (Thermo Fisher Scientific) that are capable of DNA tagmentation through active Tn5 transposase that is immobilized on the surface (fig. S1). Briefly, the barcode is made up of four segments ("A", "B", "C" and "D"), which are added to each bead through a split-and-pool procedure and together form bead-bound and barcoded Tn5 transposon heterodimers (fig. S1B). Briefly, each of the 10 μM 96 A, B, C and D segment oligonucleotides was annealed with its reverse-complement counterpart to form a double stranded segment with one base pair overhang. Single A and B double-stranded segments were immobilized to the surface of a bead (1.5 μl of each 5 μM segments per 40 μl of M-280 beads) via the streptavidin–biotin binding. Using Blunt/TA Ligase Master Mix (New England BioLabs) segments A and C, and B and D, were ligated together to form the complete and barcoded Tn5ME-B and Tn5ME-A heterodimers, respectively. Tn5 heterodimers were then single-stranded using 0.15M NaOH induced chemical lysis and re-annealed with the 19-bp Tn5MErev oligonucleotide. Haplotagging beads are activated by loading of Tn5 transposase prior to use, and can be stably stored at 4°C for at least a year.
The final "96-well Haplotagging bead plate" contains beads carrying 84,934,656 unique barcodes split into 96 wells, with all barcodes of a well identified by the C-segment, or alternatively D-segment in the initial F1(CASTxBL6) mouse experiment. Each bead of any well thus carries many copies of one of 884,736 well-specific barcodes. There are ~3.25 million beads in 5 μl M-280 Dynabeads; but only 884,736 unique barcodes in a well. Therefore, when tagmenting small quantities of samples with 5 μl of beads, pooling haplotagging beads from several wells will increase number of barcodes per sample up to the maximum of ~3.25 million barcodes per sample (e.g. pooling beads from 24 wells of the "96-well Haplotaging bead plate" makes up 21.2 million possible barcodes, so in this case, 5 μl of haplotagging beads contain approximately 3.25 million beads/barcodes from a pool of 21.2 million possible barcodes).

Sequencing library construction
Haplotag library generation involves many of the same steps of the standard Nextera/DNAflex tagmentation procedure widely used for constructing Illumina sequencing libraries. Depending on the number of samples, haplotagging beads can be added to either single DNA sample or up to 96 samples in a plate format, mixed and incubated at 55 °C for 10 minutes to tagment the DNA. Tn5 is then stripped from the DNA using 0.3% SDS. The barcoded DNA libraries are then directly amplified off of the beads with PCR using universal forward and reverse primers TruSeq-F AATGATACGGCGACCACCGAGATCTACAC and TruSeq-R CAAGCAGAAGACGGCATACGAGAT.

Single-plex haplotagging, in human and F1(CASTxBL6) mouse:
To prepare haplotagging library from a single DNA sample, 5 μl of pooled haplotagging beads (e.g. wells A1-H3 or C-segments 1-24, corresponding to a pool of 21.2 million barcodes) were transferred into one tube of an 8-tube-PCR-strip. In the next tube, tagmentation mix was prepared by adding 110 μl of WASH buffer (20 mM Tris pH8, 50 mM NaCl, 0.1% Triton X-100), 10 μl of 0.15 ng/μl HMW DNA and

30 μl of 5x TAPS-Mg-DMF buffer (50 mM TAPS pH 8.5 with NaOH, 25 mM $MgCl_2$, 50% N,N-dimethylformamide). Next, while on a magnetic stand, storage buffer was removed from the beads and the tagmentation mix was carefully transferred onto the beads with a wide-orifice pipette tip and mixed 5 times or until complete re-suspension of the beads. Sample was incubated at 55°C for 10 min to tagment the DNA. 8 μl of 6% SDS was added into the sample immediately after tagmentation; sample was mixed by inverting the tube and incubated at 55°C for 10 minutes to inactivate and strip Tn5 from the DNA. Sample was then pulse spun-down for 10 seconds and placed on a magnetic stand. Supernatant was removed and beads were washed twice with WASH buffer and kept in the second wash until further use.

With sample on magnetic stand, the buffer was removed and 30 μl of 1x Lambda Exonuclease buffer, supplemented with 10 units U of Exonuclease I and 5 units U of Lambda Exonuclease (New England BioLabs), was added to the sample to remove excess of unused barcoded Tn5 heterodimers. Sample was incubated at 37 °C for 30 minutes, and then washed twice for 5 minutes with 150 μl of WASH buffer. Bead bound haplotagged DNA was PCR amplified with Q5 High-Fidelity DNA Polymerase (New England BioLabs) in a 50 μl reaction according to manufacturer's instructions, using 4 μl of 10 μM TruSeq-F and TruSeq-R primers, with the following cycling conditions: 10 min at 72°C, 30 sec 98°C and 12 cycles of: 98°C for 15 sec, 65°C for 20 sec and 72°C for 60 sec. Library was purified and size selected using Ampure magnetic beads (Beckman Coulter) to 300–700 bp fragment size and quantified using Qubit (Thermo Fisher Scientific). The purified size-selected library was again cleaned with Ampure magnetic bead at an 1:1 ratio and adjusted with 10 mM Tris, pH8, 0.1 mM EDTA to 2.5 nM concentration.

<u>96-plex haplotagging, in Longshanks mice and butterflies:</u>
To prepare haplotagging library from 96 HMW DNA samples, 30 ng of each HMW DNA samples was diluted to 0.15 ng/μl with 10 mM Tris, pH8 in a 96-well plate and quantified with Quant-iT PicoGreen dsDNA Assay Kit (Thermo Fisher Scientific). 2.5 μl of haplotagging beads from the stock-96-well plate (~1.6 million, each carrying one of 884,736 well-specific barcodes) were transferred into twelve 8-tube-PCR-strips and closed with strip lids. On magnetic stand, the storage buffer was removed and replaced with 110 μl of WASH buffer (20 mM Tris pH8, 50 mM NaCl, 0.1% Triton X-100). Next, 10 μl of 0.15 ng/μl DNA was transferred with a multichannel pipette and 200 μl wide-orifice pipette tips strip-by-strip, and mixed 5 to -10 times to re-suspend the beads. Next, 30 μl of 5x tagmentation buffer (50 mM TAPS pH 8.5 with NaOH, 25 mM $MgCl_2$, 50% N,N-dimethylformamide) was pipetted into each strip, closed with the lid, mixed by inverting the tubes 3 to 5 times, and incubated at 55°C for 10 min. During the incubation, strip-tubes were mixed by inverting 3 to 5 times every 3 minutes. After the incubation, samples were placed on ice for 1 minute, pulse spun-down, and 8 μl of 6% SDS was pipetted into each sample to inactivate and strip Tn5 from the DNA. Samples were incubated at 55°C for 10 min, then pulse spun-down, and placed on magnetic stand. All liquid was removed and beads were washed twice with 150 μl of WASH buffer, while not disturbing the beads. Beads were kept in the second wash until further use.

Efficient linked read sequencing benefits from keeping the number of total molecules in a sequencing lane to within a range of approximately 50 – 250 million read-pairs per 1 million barcodes. This was achieved by subsampling a fraction of each sample's beads after tagmentation. For the 96-plex experiment of butterfly DNA, only 1/24$^{th}$ of each tagmentation reaction was used during PCR to decrease the relative plexity of 96-plex to a 4-plex. Thus, 1/24th of each sample's beads was transferred with a multichannel pipette, strip-by-strip, into one 8-tube-PCR-strip. This corresponds to approximately ~68.000 beads (barcodes) and ~62.5 pg DNA per sample, or ~135 haploid copies of the butterfly genome. For Longshanks mice, due to the larger genome size, only 1/10th of the beads was taken from each sample. With only 8 samples on the magnetic stand, the buffer was removed, and 30 $\mu$l of 1x Lambda Exonuclease buffer, supplemented with 10 units of Exonuclease I and 5 units of Lambda Exonuclease (New England BioLabs), was added to each sample. Samples were incubated at 37 °C for 30 minutes, and then washed twice for 5 minutes with 150 $\mu$l of WASH buffer.

DNA library was then amplified using Q5 High-Fidelity DNA Polymerase (New England BioLabs) in a 50 $\mu$l PCR reaction according to manufacturer's instructions, using 4 $\mu$l of 10 $\mu$M TruSeq-F and TruSeq-R primers, with the following cycling conditions: 10 min at 72°C followed by 30 sec 98°C and 10 cycles of: 98°C for 15 sec, 65°C for 30 sec and 72°C for 60 sec. Libraries were pooled after PCR into a single library pool, size selected using Ampure magnetic beads (Beckman Coulter) Qubit quantified, and adjusted with 10 mM Tris, pH8, 0.1 mM EDTA to 2.5 nM concentration for sequencing.

Sequencing and demultiplexing

Pooled libraries were sequenced by a HiSeq 3000 (Illumina) instrument at the Genome Core Facility at the MPI Tübingen Campus with a 150+13+12+150 cycle run setting, such that the run produced 13 and 12nt in the i7 and i5 index reads, respectively. Sequence data were first converted into `fastq` format using `bcl2fastq v2.17.1.14` with the following parameters `--use-bases-mask=Y150,I13,I12,Y150 --minimum-trimmed-read-length=1 --mask-short-adapter-reads=1 --create-fastq-for-index-reads`, or if using Illumina's sample demultiplexing feature with a 150+12+13+150 run configuration: `--use-bases-mask=Y150,Y12,I7Y6,Y150 --minimum-trimmed-read-length=1 --mask-short-adapter-reads=1 --create-fastq-for-index-reads --barcode-mismatches=0` (Illumina; and where applicable, demultiplexed samples by the "C" or "D" segments of the beadTag barcode).

Then we performed beadTag demultiplexing to generate the modified `fastq` files using a custom programme `filterFastq_by_bc`, resulting in a `fastq` file supplemented with beadTag information. This programme is available at https://github.com/evolgenomics/haplotagging.

Analysis and phasing of molecules

The beadTag-annotated `fastq` files were then mapped onto reference genome assemblies using `bwa v0.7.10-r789` (*58*) and processed using `samtools v1.2` (*59*), marking and ignoring PCR and optical duplicate reads in subsequent analyses. For the human GM12878 sample, a set of "gold standard" variant positions were examined, made available by the Genome-in-a-bottle consortium website (*56*). For mouse samples, We examined a set of 6,620,436 biallelic SNP positions in the genome positions known to be different between the C57BL/6NJ and CAST/EiJ strains, made available by the Wellcome Trust Sanger Centre (Mouse Genomes Project version 5 dbSNP v142 release (*60*).
For all files, we processed them following the same dual-pronged pipeline: first, we determined molecules from sets of reads sharing the same beadTag encoded with the `BX` tag with a maximum gap size of 50 kbp using custom `Perl` and `bash` scripts.

These provided basic statistics on the molecule size, molecular coverage and reads per molecule. Second, we extracted reads overlapping phase or haplotype-informative positions following the pipeline as outlined by `HAPCUT2` (*10*). These phase-informative molecules were used to determine the number of phase-informative reads per molecule, informative molecule size, haplotype phasing, phase blocks, as well as short and long switch error rates for molecules spanning at least 4 phase-informative SNPs. The results are summarized in Table S4.
We then parsed the beadTag output to identify "molecules". We also followed the definition used by `longranger` and defined each molecule as a cluster of reads sharing the same beadTag within 50 kbp of each other. We then analyzed the molecules for the SNP alleles and classify them as "concordant" if a given position belongs to the majority allele and otherwise as "discordant" positions. We discarded molecules overlapping 2 or fewer SNPs, and assigned phasing of each molecule to Haplotype 0 or Haplotype 1 if they carry one or no discordant positions. We classified molecules carrying 2 or more discordant positions as "mixed molecules".

Comparison to 10X Chromium-generated data
Sequencing data was generated from the same DNA extraction as used in haplotagging. We subsampled the mapped `BAM` file to match the sequencing depth from the haplotagging dataset while retaining the paired-end structure by using `samtools` view module with the `-s` option. We then applied the exact same analysis and phasing pipeline as described above, in order to allow direct comparison of the performance of the two techniques in recovering phasing information from the F1(BL6xCAST) hybrid mice.

Heliconius *butterfly field sampling*
In all sampling sites, butterflies were captured using entomological nets, their wings preserved in glassine envelopes and their bodies in DMSO buffer solution (*61*) to preserve their DNA for future molecular genetic analyses. At each sampling site along the transect, we aimed to collect at least 30 individuals of the more abundant species *H. erato* individuals, the most abundant of the two species, and as many *H. melpomene* as could be collected in the same time. Field sampling was performed in several trips to the study area: May 2009, December 2009 – March 2010, July – September 2010 and February 2011.
Subsequent to sampling, genetic analysis demonstrated that cryptic individuals of the morphologically indistinguishable *H. timareta* exist among the lowland samples of *H. melpomene (28)*. All putative *H. melpomene* samples from the lowland were thus tested for their mitochondrial *COI* genotype following (*62*). In addition, it turned out later that three individuals from the hybrid zone centre clustered together with *H. timareta* in a PCA analysis that was performed with `pcangsd` (*63*) on the haplotagging data of all *H. melpomene* samples and three *H. timareta* samples. These three *H. timareta* samples were excluded from all analyses.

Population phasing and imputation
Three linked read dataset from large set of population samples were generated: mice from generations F11, F16 and F17 from the Longshanks selection experiment (N=245) (*12*, *13*) and butterflies from the two species *Heliconius erato* (N=484) and *Heliconius melpomene* (N=187) from an overlapping hybrid zone in Ecuador (see section below for details). In all three datasets, the initial pipeline involved beadTag demultiplexing, read trimming, placement (against `mm10`, `Hmel2.5` and `heleral_demo` respectively), duplicate marking, molecule identification and initial SNP calling using `bcftools call` with the multiallelic calling algorithm (`-m`) ([NO

STYLE for: ]). The set of BAM files and the raw SNP set were then used as input for the statistical phasing program STITCH (*11*). Initial parameter tuning were performed to maximize call rate and genotype concordance at focal loci with known genotypes or major color pattern loci: Chr5:415,36,431 (rs33219710) and 41,536,498 (rs33600994) for the Longshanks mice (*13*); *WntA*, *optix* and *Ro* for *H. erato*; and *WntA*, *optix* and *cortex* for *H. melpomene* (Table S8). The color loci are co-dominant except for *Ro* in *H. erato* and *cortex* in *H. melpomene (27)*.Then these parameters were refined to maximize informativeness (INFO_SCORE) and SNP diagnostic statistics such as the transition/transversion (TsTv) ratio. Phasing parameters for the different datasets are shown below in their own sections.

Evaluation of phasing performance under subsampling

In the Longshanks dataset, the chosen non-default key parameters were: `method=diploid, nGen = 20, K = 4, iSizeUpperLimit = 500000, niterations = 60, readAware = TRUE, shuffleHaplotypeIterations = c(5, 10, 15, 20, 30, 40), reference_iterations = 40, reference_shuffleHaplotypeIterations = c(4, 8, 12, 16), maxDifferenceBetweenReads = 1e10`. Here, our main objective was to determine if linked read information would improve phasing performance. STITCH features a read-aware mode in standard paired-end reads using the shared read name. To incorporate linked read information, we substituted the read names with the BX tag. We compared the average INFO_SCORE and the imputed allele frequencies. For imputation runs using linked reads, an additional parameter `splitReadIterations = c(20, 40)` was used to split the linked reads where appropriate. We also subsampled each input BAM in two different ways to simulate progressively high multiplexing: by randomly picking read pairs or by molecules. Molecular subsampling is appropriate for haplotagging, because during the multiplexing of a large number of samples, only a small subset of beads, and thus molecules, from each sample is used (see section “Sequencing library construction” for details). Subsampling by random read pairs would have disproportionately diminished the information content of linked reads. The results are shown in fig. S2.

Population genomics analysis of the parallel *Heliconius* hybrid zones

For the *Heliconius* datasets, the chosen non-default parameters were: `method = diploid, nGen = 500, readAware = TRUE, --shuffle_bin_radius=500 --expRate=5 --iSizeUpperLimit=500000, niterations = 40`. A K value of 30 was used for *H. erato* and 40 was used for *melpomene*. These parameters were applied genome-wide in windows of 1 and 0.5 Mbp (with an overlap of 25 kbp between adjacent windows) for *H. erato* and *H. melpomene* respectively. The resulting imputed variant call files (VCF) were merged and annotated with allelic depth and coverage from the original BAM files. We also removed positions with poor information content (`INFO_SCORE <= 0.2`). This call set contained 49.2 million SNP positions for *H. erato* (131 SNPs / kbp) and 26.3 million for *H. melpomene* (97 SNPs / kbp).

For each individual, molecular phasing using HAPCUT2 was performed at actual or likely heterozygous positions in the “10X” or linked read aware mode with a maximum gap of 50 kbp, a phase block PHRED-scaled threshold of 30 and the `call_homozygous` option. The resulting VCF file thus integrate both STITCH-imputed genotypes and phasing information from HAPCUT2. For population genomics analyses using NGSadmix and angsd (*65*, *66*), more stringent INFO_SCORE filter at 0.5 was used.

Population structure

In order to assess the genetic variation across the hybrid zone, we inferred admixture proportions for each species separately. We used the genotype likelihoods computed by STITCH to generate the input files with angsd v0.931 (*66*). angsd is well-suited for low-coverage sequencing data as it takes into account genotype uncertainty by using genotype likelihoods in a Bayesian framework. To infer admixture proportions, we used NGSadmix (*65*), a maximum-likelihood method which uses genotype likelihoods to infer genomic clusters and assign ancestry proportions to each individual. We ran NGSadmix on sites with a minor allele frequency of at least 0.05 and specified two clusters (K=2) or three clusters (K=3). In order to infer the optimal number of clusters, we ran 10 replicates each with K=1 to K=4 and assessed the optimal number of clusters with the method by (*67*) implemented in Clumpak (*68*).

Signatures of selection

We computed genetic differentiation ($F_{ST}$) between the subspecies using phenotypically pure individuals from the two sampling sites at either end of the transect (32 *H. e. notabilis* and 28 *H. e. lativitta* and 41 *H. m. plesseni* and 40 *H. m. malleti*). We used the HAPCUT2-emitted genotype likelihoods in angsd to calculate the site frequency spectrum (SFS) for each population using the method by (*66*). Then, we computed folded 2D-SFS for both populations together which we used as a prior for the joint allele frequency probabilities at each site to compute per-site $F_{ST}$ values using the Weir–Cockerham correction (*69*) as implemented in angsd. Finally, we computed weighted $F_{ST}$ averages across adjacent 10 kbp windows, excluding windows with less than 100 SNPs. In order to infer divergence peaks, we ran a Hidden Markov Model (HMM) analysis with two states (high and normal differentiation) on normalized $F_{ST}$ values (z-scores=number of standard deviations from the mean). Following (*70-72*), we optimized transition and emission probabilities with the Baum–Welch algorithm (6 runs with 1,000 iterations each (*73*)) and inferred the most likely sequence of states given the observed z-scores with the Viterbi algorithm (*74*). To best compare the high differentiation peaks between the two species, we fixed the means of the state distributions to 0 (normal) and 3 (high differentiation) and the standard deviations to 1.

Next, we identified signatures of selection by computing the nucleotide diversity (π) for each race with the same individuals as above in windows of 10 kbp and 50 kbp using VCFtools v0.1.14 (*75*). We expect decreased nucleotide diversity in the race where a selective sweep occurred. We computed the difference in nucleotide diversity between highland and lowland races (Δπ, specifically $\pi_{notabilis} - \pi_{lativitta}$ and $\pi_{plesseni} - \pi_{malleti}$).

We calculated integrated haplotype score (iHS) using selscan v.1.2.0a (*35*, *76*), excluding low-frequency variants and based on a genetic map interpolated onto our SNP sets. Raw genome-wide values were then normalized using the associated norm v.1.2.1a utility, with 100 frequency bins over 10 kbp windows. The genetic maps and the code for interpolation is available at our Dryad data repository.

We calculated ω using OmegaPlus v3.0.3 (*77*) and the composite likelihood ratio according to Nielsen et al, 2005 using sweeD (*78*). It implements the ω statistic proposed by Kim and Nielsen, 2004 (*34*). To further minimize phasing errors, we coded all low-confidence sites as unknowns, and ran OmegaPlus under binary mode with imputation. For each scaffold we performed exhaustive searches with a grid spacing of 10 bp, and report the $\omega_{MAX}$ found in a 10 kbp window. For SweeD we used a grid spacing of 1 kbp.

For comparison between the species we mapped the *H. melpomene* windows to the *H. erato* reference genome with the `liftOver` utility (*79*) using a chain file. See section *Cross-species alignments* on details on the chain file.
If entire windows are lifted over between species, the windows of the two species start at different positions and are thus not directly comparable. In order to assess the interspecific correlation in genome-wide patterns of differentiation, we thus defined 10 kbp windows for *H. melpomene* directly on *H. erato* reference genome positions. First, we extracted alpha and beta values for each *H. melpomene* SNP with `realSFS fst print`. Next, we mapped the position of each *H. melpomene* SNP to the *H. erato* reference genome with `liftOver`. To increase the mapping success, we added 5 kbp flanking regions on either side of the SNP. We combined multiple mapping locations with `bedtools groupby` and used the middle position as the *H. erato* position of the *H. melpomene* SNP. We computed weighted $F_{ST}$ averages for 10 kbp windows as the ratio of average alpha and average beta values (*80*).

Identifying the genomic regions underlying color pattern differences

We performed association mapping to identify genomic regions that control wing coloration. Again, we used `angsd` (*66*) to account for genotype uncertainty. We used the admixture proportion inferred with `NGSadmix` as a co-variate to account for population substructure in our dataset. Elements of color patterns of all individuals were scored using standardized photos, informed by trait segregations in controlled lab crosses (*27*). We performed likelihood ratio test (*81*) and classified the phenotypes as quantitative traits (`-yQuant` option) with an additive model (`-model 1`). For phenotypes where heterozygotes could not be identified visually, we used a binary classification (`-yBin option`) with a dominant model (`-model 2`). Sites with a minimum allele frequency of 0.05 were used to compute 10 kbp sliding window averages with a step size of 2.5 kbp.

Cross-species alignment

Genome assemblies from *H. erato demophoon* (`heleral_demo`) and *H. melpomene melpomene* (`Hmel2.5`) were first masked for repeats using `trf v4.07b (82)` and then aligned to each other using `lastz v.1.03.54 (83)`, with the following parameters: `C=0 E=150 H=0 K=4500 L=3000 M=254 O=600 Q=human_chimp.v2.q T=2 Y=15000` optimized for closely-related species. The resulting all-to-all pairwise alignments were grouped and alignment hits chained using the `axtChain` and `chainMergeSort` programmes. The distribution of scores were examined and only hits above 14000 were retained.

Comparison of haplotagging data to RAD data

An alternative to sequencing many whole genomes at low coverage is to sequence many individuals at only a fraction of the genome, e.g. by using restriction site associated DNA (RAD) sequencing (*84*). In order to compare the two techniques directly, we reanalyzed previously published RAD sequencing data of the populations at the edges of the Ecuadorian hybrid zone of *Heliconius erato (28)*. This RAD dataset consists of ten individuals each of *H. e. lativitta* and *H. e. notabilis* (130,332,160 mapping reads) and uses a frequent restriction enzyme (*PstI*, cutting 5′-CTGCAG-3′). To follow the same procedure as in the haplotagging data, we aligned the reads to the *H. erato* reference genome with `bwa mem v0.7.12`. We called variants and genotypes with `bcftools mpileup v1.9`. Using `vcftools v. 0.1.15`, we filtered out genotypes with less than 5 reads or genotype quality below 20. Additionally, we excluded sites with more than 25% missing data, with

SNP quality below 30 or with less than two alternative allele counts. We computed $F_{ST}$ in 10 kbp sliding windows (2.5 kbp step size) using `vcftools`. In order to get a comparable haplotagging dataset, we computed 10 kbp window $F_{ST}$ averages with `angsd` as explained above but with ten individuals per subspecies of the same populations as those used for RAD sequencing (127,204,006 mapping reads). To match the number of reads, we selected the 10 individuals with lowest number of reads for each subspecies, but excluding the *H. e. lativitta* sample with the lowest number of reads.

Haplotype length at major color loci

We are interested here in identifying the key switch events that switch a haplotype from highland to lowland type or vice versa around the major loci *optix* and *WntA*. To do so we first identified and polarized ancestry-informative SNPs within 0.5 Mbp of the focal position, generally defined as those showing a frequency difference ≥ 0.5, or 0.8 at *optix* in *H. erato*, which shows greater differentiation. Next, we used a set of heuristics to screen and remove potential switch errors from the raw `STITCH/HAPCUT2` output, e.g., by filling in low-confidence calls or non-calls by minimizing crossovers and removing double-switch events where both alleles switch between high- and lowland haplotypes. These switch errors occur due to low local molecule support, genuine phasing errors, or trivially between adjacent SNPs that are assigned into different phase blocks. We then identified long switches over 4 SNPs that transitions between high- and lowland types. The span of the haplotype between the closest flanking breakpoints were defined as the “haplotype length” around the selected loci and visualized in fig. S8.

Interval mapping at the *H. melpomene optix* region

We compared the phenotypes of rare recombinants at the *optix* region to narrow down the regulatory elements controlling the absence and presence of red scales in the forewing bands and the “dennis patch” in the forewing and the “dennis bar” and “rays” in the hindwing (see Fig. 2 and fig. S3). We filtered out genotypes with quality below 10 and sites with an `INFO_SCORE` below 0.7 or more than 25% missing data. Next, we extracted all sites that showed a difference in homozygote genotype frequencies of above 0.9 between individuals grouped by *plesseni*-like (“highland”) or *malleti*-like (“lowland”) red patterns. For ease of visualization of haplotypes, we corrected genotyping errors and replaced missing genotypes the following way. For each individual, we replaced runs of up to 15 consecutive genotypes that differed from the flanking 20 genotypes by the genotypes of the flanking region given that the flanking region was clearly assignable to H/H (=1, homozygous for the highland allele) or H/L (=0.5, heterozygous) or L/L (=0, homozygous for the lowland allele). If the mean genotype value of the flanking region deviated more than 0.1 from 0, 0.5 or 1, the region was defined as not clearly assignable. Each individual was “cleaned” three times to remove likely erroneous genotypes which are expected to be visible as single genotypes or short runs of genotypes that differ from the surrounding regions. The genotypes were visualized as yellow for the highland homozygotes, as orange for heterozygotes and as red for the lowland homozygotes. Rare recombinants between the different regulatory regions of *optix* previously found by (*55*, *85*) were then compared to the photos to determine the genetic basis of each red pattern.

Structural rearrangements and analyses

Structural rearrangements were detected based on beadTag between adjacent windows following (*7*). Briefly, a bash pipeline was used to summarize beadTag found in any given 10 kbp window across the genome as well as between any pair of

10 kbp windows on the same chromosome. On average, we observed 25.8K ± 22.7K molecules per window, of which 5314 ± 1658 molecules are shared across adjacent windows. By contrast, we observed an average of 9.2 ± 31.3 shared beadTags between windows that are more than 100 kbp apart. These patterns are summarized across the genome (Fig. 4). Signals of potential inversions and insertion/deletions (indels) were manually inspected and scored. The results are summarized in Data S2 & S3.

To estimate the frequency of the *H. erato* Chr2 inversion, first we determined the non-adjacent windows with the strongest beadTag sharing. Then among these molecules we inferred the inversion junctions using the positions with the lowest base-wise molecular coverage, because the junction itself is likely to fall within gaps between linked reads from either direction, and to avoid inflating the count of collinear molecules. For the Chr2 inversion we determined the junctions to be at Herato0204:172500 and 1290057. Then each of the 484 individuals were scored for beadTags that span either junction, or ones that are shared between $L_{out}$ – $R_{in}$ or $L_{in}$ – $R_{out}$ windows. We next determined the average inversion frequency at each sampling site and fitted cline models to them (see below).

<u>Cline analysis</u>

One-dimensional analyses of cline position and shape were carried out using a subset of the sampled locations; these included all the best-sampled sites along the straight line where sampling was concentrated. The sequential location of sites along this transect was defined by projecting their position perpendicularly to a regression line fitted to their geographic coordinates and expressed in km (Fig. 2). For genotypic clines, the allele frequencies at SNPs that separate the highland–lowland divergent haplotypes at the peak $F_{ST}$ windows were used to construct the clines. At the major color pattern loci, the best model (according to the Akaike Information Criterion, AIC) was found to be a 4-parameter model with free start and end frequencies. Maximum likelihood estimates of cline centre, width and start and end frequencies were then obtained for the allele frequency clines of the two co-dominant and homologous loci, *optix* and *WntA*, as well as the two dominant loci, *Ro* in *H. erato* and *cortex* in *H. melpomene* using the `R` software package `HZAR` (*86*). Cline coincidence and concordance was evaluated by comparing the overlap of likelihood limits between loci.

To assess the generality of the observed cline widths and centres, we additionally inferred clines for the site with the highest $F_{ST}$ value for each 100 kb window using the same populations for $F_{ST}$ calculation as used for previous $F_{ST}$ calculations. We excluded windows in which none of the sites reached an $F_{ST}$ value of 0.5 in *H. erato* or 0.4 in *H. melpomene*. We then inferred clines for the remaining 137 sites in *H. melpomene* and 132 sites in *H. erato* with `HZAR`. For each site, we tested six models (no tails, symmetric tails or two different tails each combined with either free or fixed minimum and maximum allele frequencies). Following the example by (*86*), we ran 3 MCMC chains each with default parameters and randomized starting values. We identified the best model of the six and a null model of allele frequency independent of the transect position with `AICc` as implemented in `HZAR`.

### Supplementary Figures

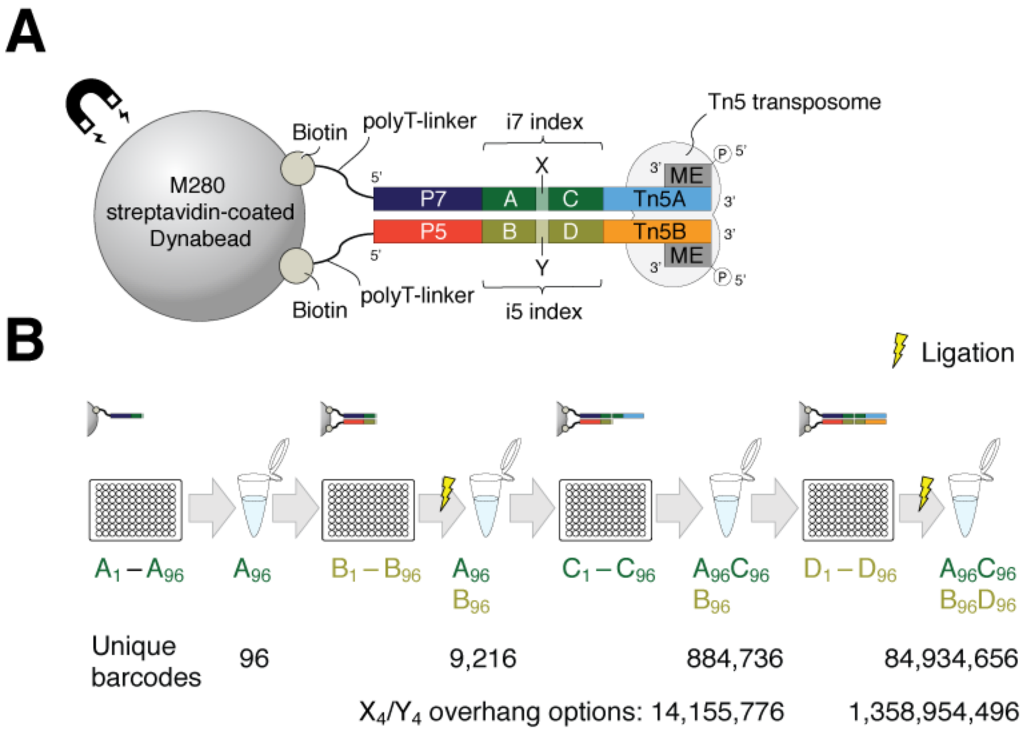

**Fig. S1. Haplotagging bead assembly.** (**A**) The design of a haplotagging bead. Haplotagging beads are microbeads coated with activated Tn5 transposomes that correspond to the Nextera specifications. The key feature is a set of segmental barcodes ("beadTag") that is integrated into the i7 and i5 indexing positions. In the current design, we use two segments each (designated A – C and B – D), linked by a single basepair overhang (X and Y). These oligonucleotides are attached to the bead via the strong streptavidin–biotin binding. An advantage of this design over other similar designs (*8*) is that there is no intervening adaptor sequences (which requires custom sequencing primers), nor is there major presence of splint hybridizing sequences (which would greatly extend the length of the indexing sequence), either of which would prevent the standard TruSeq sequencing protocol to be used on an Illumina sequencer. (**B**) Assembly of the combinatorial beadTag barcode via a split-and-pool procedure. Pre-suspended 96-well plates bearing oligonucleotides are ordered directly from suppliers. Commercial streptavidin-coated dynabeads are aliquoted into each well, pooled, and then re-aliquoted into each well into the next plate. At each step, an individual microbead would be mixed with a single type of barcodes, but as a pool of beads, the entire mixture would feature up to approximately 85 million combinations. If the X and Y overhangs are varied, this can feature up to 1.4 billion combinations.

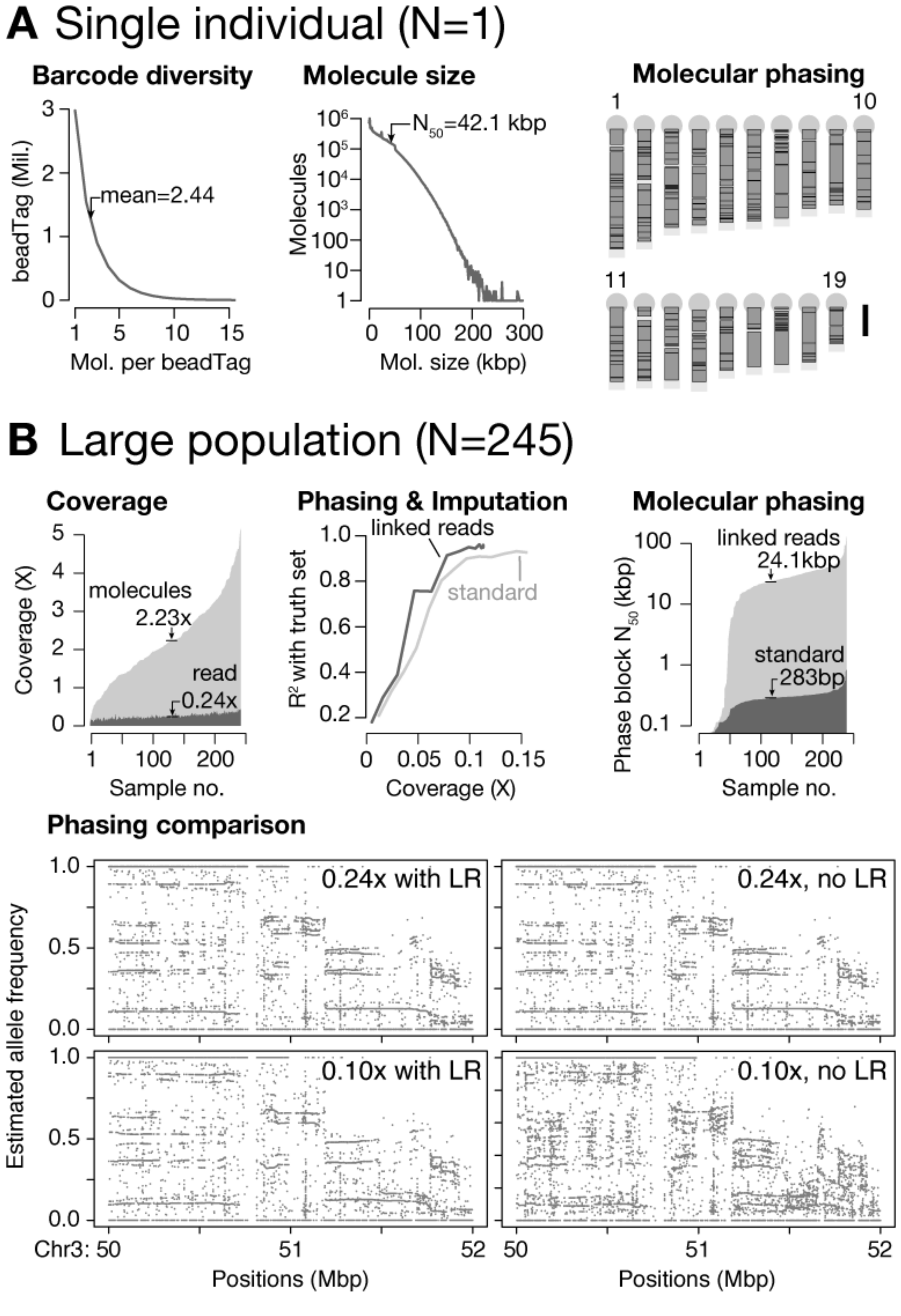

**Fig. S2. Phasing and imputation performance in single individuals and a large population.** (**A**) Barcode diversity, molecule size and phased block sizes from the same F1(BL6xCAST) sample. On average, each barcode is found on only 2 to 3 molecules scattered across the genome. Half of the genome is covered by molecules 42.1 kbp or longer. Phasing was successful across virtually the entire genome. Shown here are the largest phase blocks (dark grey boxes; up to 90% of the total length of all phase blocks, or $N_{90}$) on the 19 autosomes of the mouse. Scale bar: 50 megabase. (**B**) LR sequencing, phasing and imputation results from haplotagging 245 mice from the Longshanks selection experiment (*12*, *13*). LR sequencing allows molecular coverage (median: 2.23x)—as opposed to standard per-base read coverage (median: 0.24x)—to be leveraged across samples to infer and extend haplotype segments. Phasing and imputation while incorporating LR information consistently shows higher correlation with allele frequencies estimated from higher-depth sequencing (*13*) than standard short-read only attempts, even after subsampling to as low as 0.05x coverage per individual mouse. LR data also lead to a 100× increase in phased block lengths. Bottom: representative results from statistical phasing with down-sampled input, with or without LR information. Haplotypes can be visualized by runs of correlated allele frequencies. In this 2 Mbp region, phasing with or without LR information at 0.24x coverage show comparable results. By contrast, at 0.1x coverage, phasing remain robust using LR information (left; the sharper appearances of correlated frequencies suggest possibly improved phasing results) compared to poor phasing results if the input was treated as standard, paired-end reads (right).

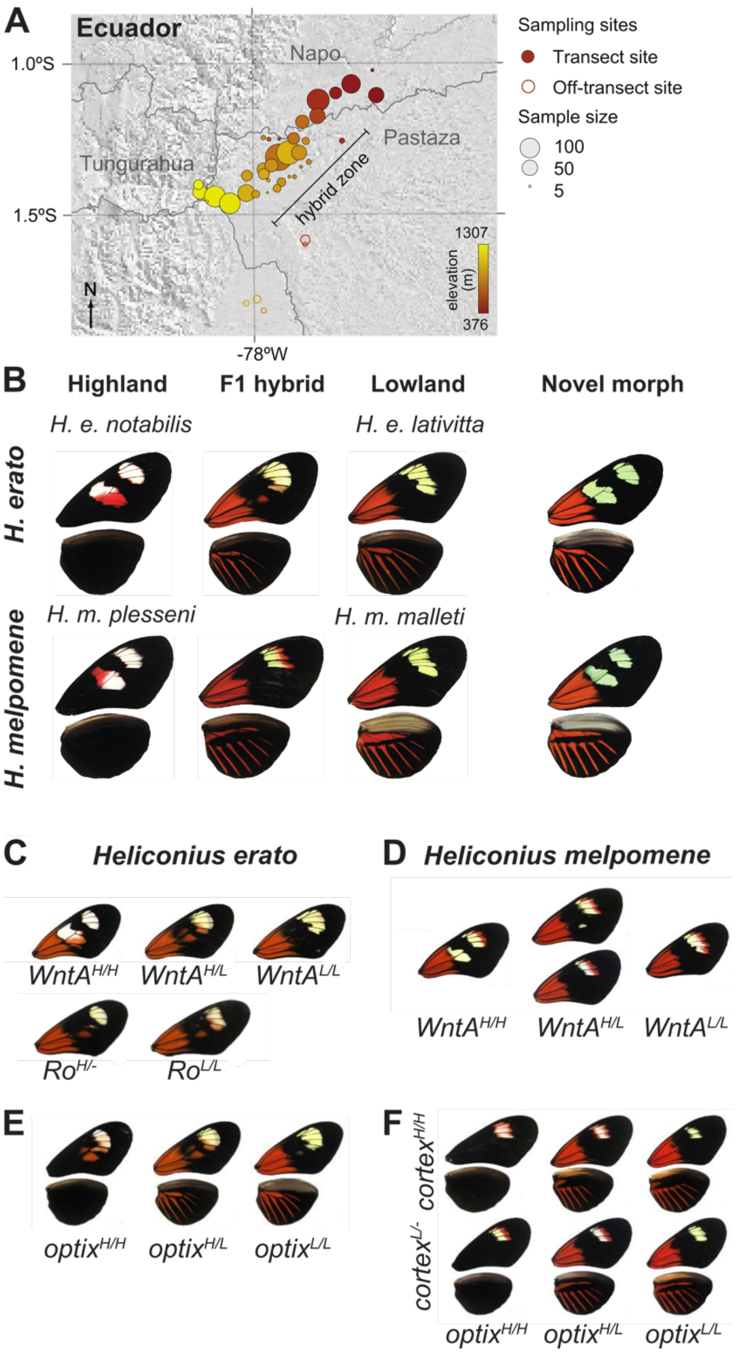

**Fig. S3. Sampling localities and representative morphs.** (**A**) Sampling localities shown in Fig. 2A are shown at higher magnifications here. The color coding corresponds to the same scale as in Fig. 2A. The transect is located along a Southwest–Northeast diagonal. (**B**) Representative individuals of the pure races, F1 and the new hybrid morph in *H. erato* and *H. melpomene*. (**C–F)** Representative individuals visualising how different genotypes at *WntA* (**C–D**) in both species and at *Ro* in *H. erato* affect the forewing band shape (**C–D**), and how genotypes at *optix* in both species and *cortex* in *H. melpomene* affect the distribution of red scales (**E – F**).

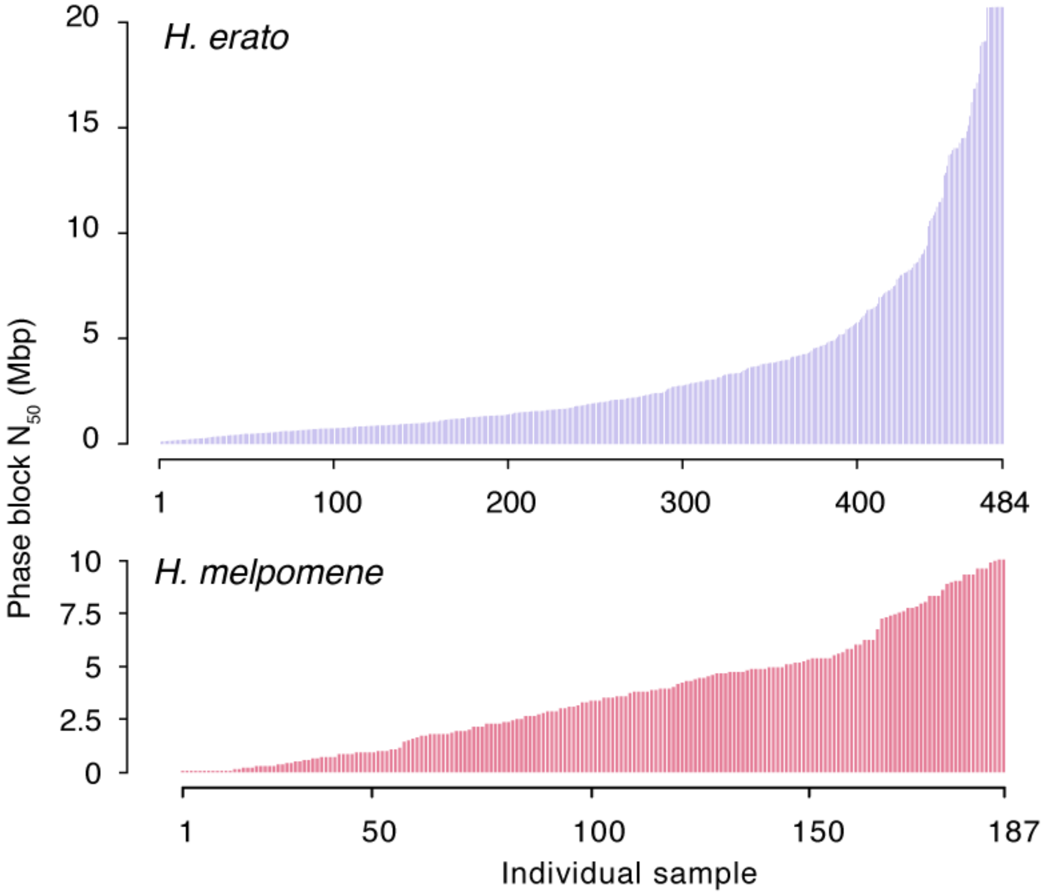

**Fig. S4. Phasing performance in the two butterfly species.** Following statistical phasing, each individual was also phased using molecular information across its imputed heterozygous sites using HAPCUT2. The phased block $N_{50}$ is shown for *H. erato* (top) and *H. melpomene* (bottom). Among *H. erato*, the maximum phased block $N_{50}$ is 20.7 Mbp, which spans the entire Herato1202 scaffold, the third longest scaffold in that assembly.

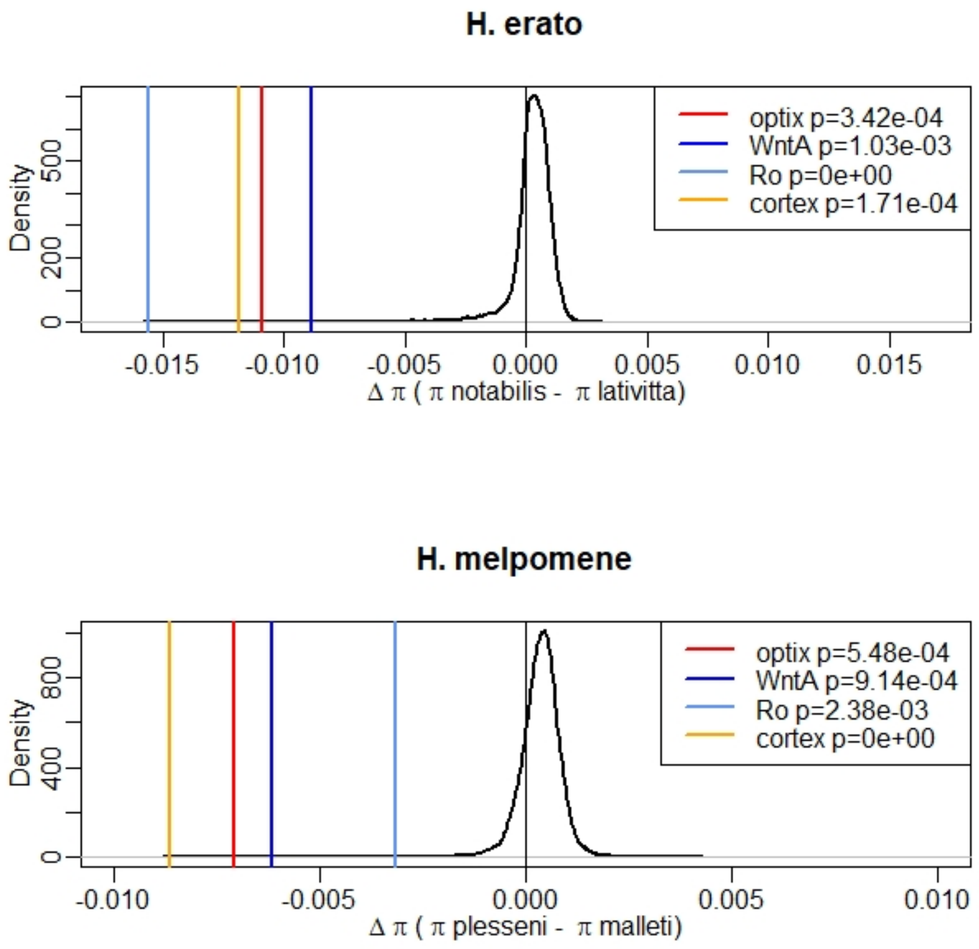

**Fig. S5. Extreme difference in nucleotide diversity at the four major divergent loci.** In both *Heliconius* species, the difference in nucleotide diversity between highland and lowland races (Δπ) was computed in 50 kb windows. The density distribution of Δπ value across the genome is shown in black, with the most extreme 50 kb window at each color pattern locus indicated as vertical colored lines. Color pattern loci show strongly negative Δπ values, indicating stronger reduction in nucleotide diversity in highland races than in lowland races. Empirical one-sided *P*-values are given for each color pattern locus.

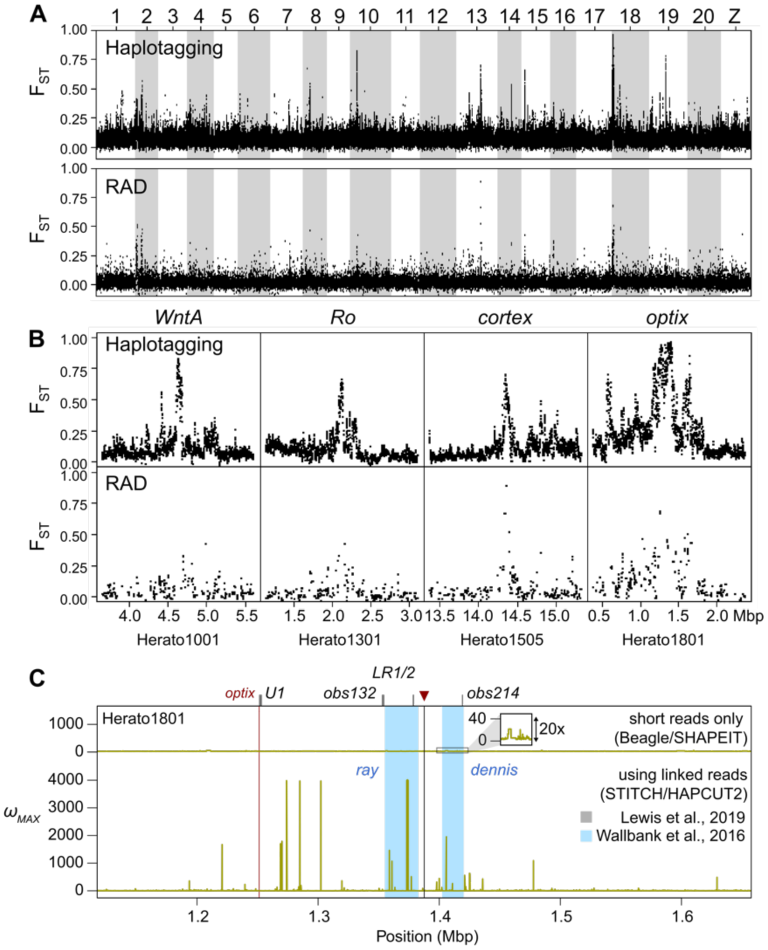

**Fig. S6. Haplotagging data out-perform conventional short-read alternatives.** Patterns of genomic differentiation ($F_{ST}$) across the genome show much higher resolution in haplotagging data (**A**) than in RAD data (**B**) despite the same number of individuals (10 individuals from each population) and comparable number of mapped reads (haplotagging: 127 million vs. RAD: 130 million). $F_{ST}$ values were calculated in sliding windows of 10 kbp with a step size of 2.5 kbp. Windows with less than 10 SNPs were excluded. There are many more regions with marked differentiation using haplotagging data. Most of these regions are validated in the broader, main dataset presented in Fig. 3A. (**B**) The difference in resolution is particularly obvious at the four regions of highest differentiation. (**C**) Comparison the haplotpe-based ω-stastics (*34*), which detects LD signatures associated with genetic hitchhiking with or without LR information. The same data from 32 *H. e. notabilis* individuals were processed using either the STITCH/HAPCUT2 LR pipeline outlined in this paper, or a standard Beagle/SHAPEIT pipeline without using LR information. The ω test searches for increased LD within each flanking area adjacent to the inferred target of selection, but not across it. It is sensitive to accurately constructed haplotypes. The LR pipeline shows a peak $\omega_{MAX}$ of 4014.5 in the region, especially in the area immediately flanking the strongest association with the wing pattern phenotypes (red arrowhead and black vertical bar) that is more than 100 kbp 3' to the coding region of *optix* (red vertical bar). The major signals correspond to regulatory regions (blue shading: *ray* and *dennis* according to (*54*)) and even overlap particular regulatory elements in this region (grey ticks above plot, labelled according to (*25*)). By contrast, the maximum ω-statistics at this locus without using LR information is 29.3 (inset, magnified 20×) and seem to fluctuate.

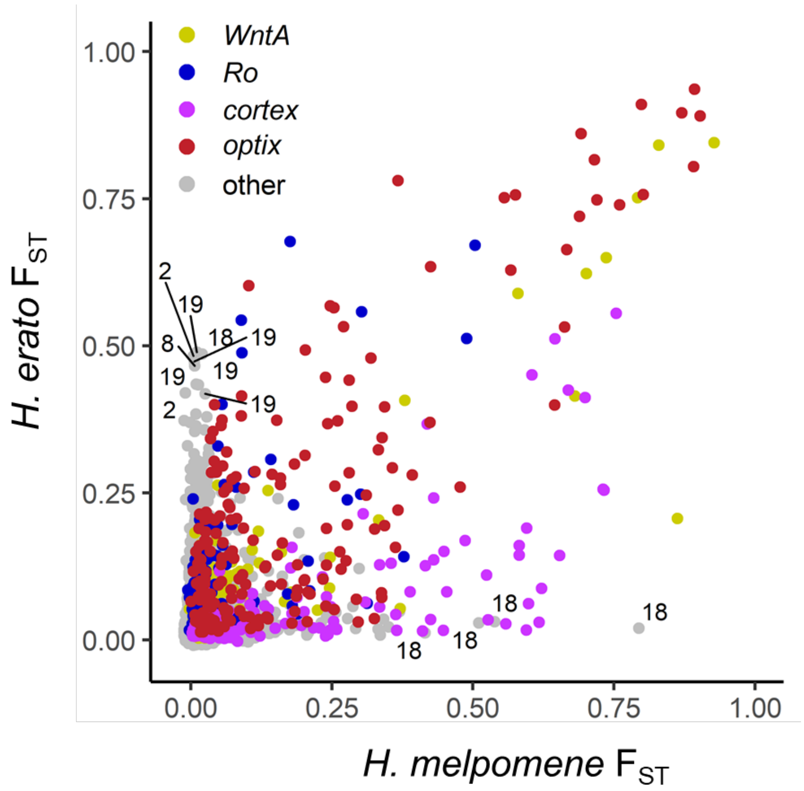

**Fig. S7. All 10kb windows with high genomic differentiation ($F_{ST}$) in both *H. erato* and *H. melpomene* belong to the four major color loci.** $F_{ST}$ computed in 10kb windows between highland and lowland races for each species separately, (*H. melpomene* data were converted to *H. erato* coordinates before averaging). All windows within 0.5 Mbp from the centre of the four major color loci in *H. erato* are shown in color. For windows with an $F_{ST}$ value above 0.4 that are not part of the major color loci the chromosome number is indicated. The four windows that are highly differentiated only in *H. melpomene* are all located on chromosome 18 and are part of a second divergence peak about 2 Mbp away from *optix* unique to this species. This region also shows a steep cline in allele frequency which is coincident with the *optix* cline. Windows that are highly differentiated only in *H. erato* include a region on Chr2 which also shows steep clines shifted towards lower altitudes even compared to the *WntA* cline (fig. S12). This region encompasses six genes of which four are putatively related to diet (fatty acid synthase, trypsin, gustatory receptor for sugar taste, odorant-binding protein).

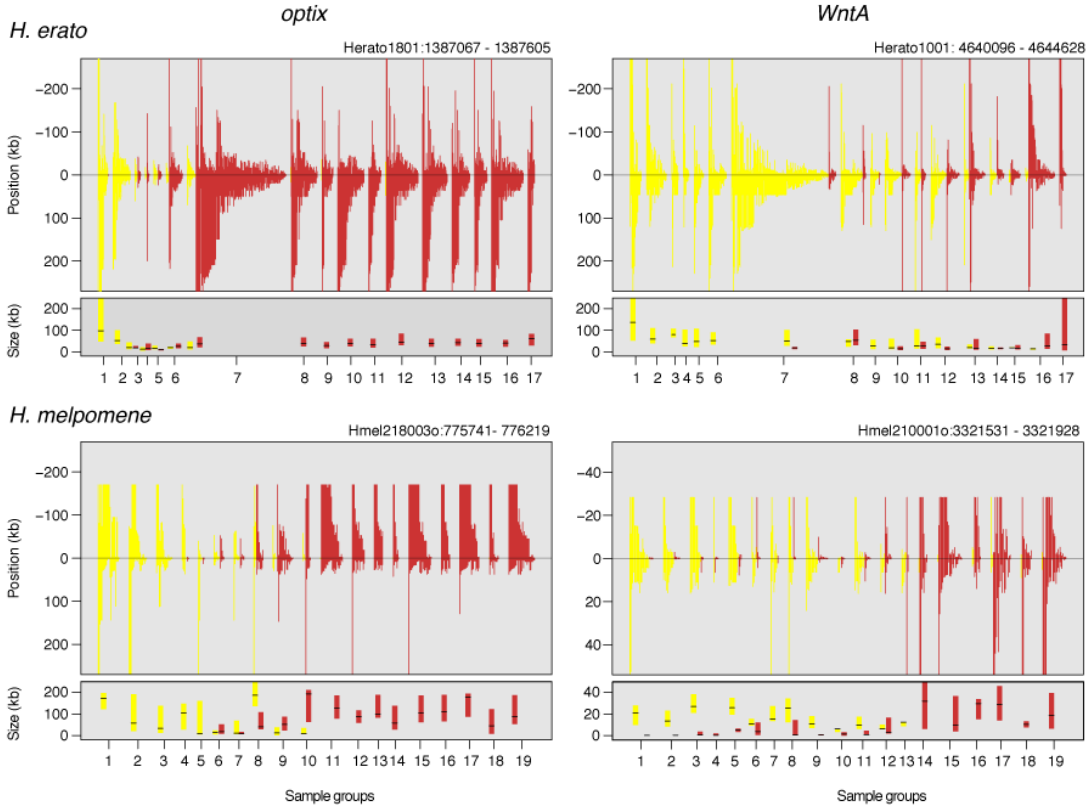

**Fig. S8. Haplotype length distributions at the major color loci *optix* and *WntA*.** Haplotypes from inferred selection targets are plotted in sample groups (Top: *H. erato*; bottom: *H. melpomene*). In each sample group, haplotypes are assigned to into highland (yellow), lowland (red), or ambiguous types (not plotted here), and plotted from the longest to the shortest length, calculated from the closest recombination breakpoints flanking the centre, focal position (see Methods for details). Summarized below each plot is a box plot depicting the median and the interquartile range of the haplotypes in each group (with a minimum of 3 haplotypes). To help visualize the breakdown of the average haplotype in the middle of the hybrid zone, some bars may be truncated at the top. This representation clearly shows the displacement between the *optix* and *WntA* clines. It also shows that haplotype lengths tend to be long at both ends of the hybrid zone, and become broken down through hybridization at the centre of each cline. Within a sample group, comparing the size distribution of haplotypes of each type may also reveal the direction of introgression. Note that the *H. melpomene WntA* locus contains far shorter haplotypes than those at the other depicted loci, and is plotted with a different Y-axis.

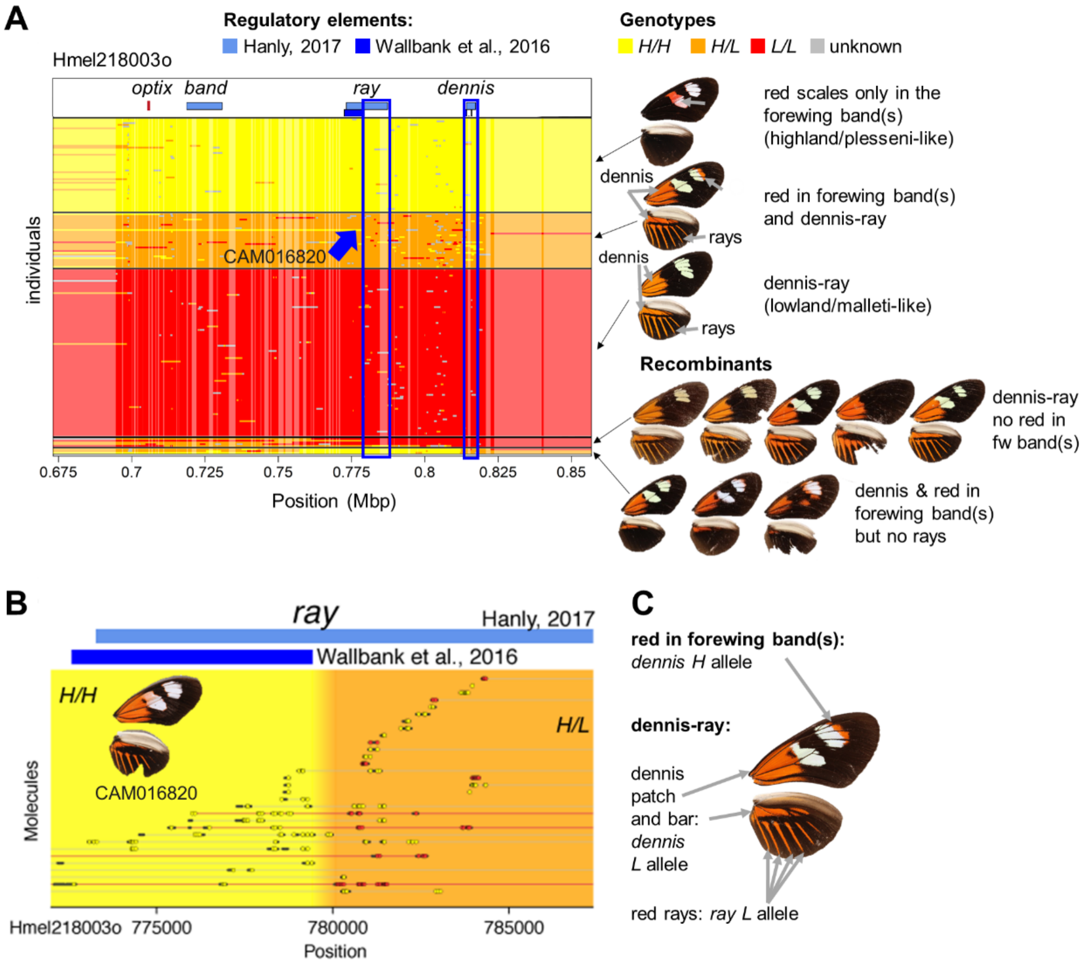

**Fig. S9. Refining the minimal intervals at the *optix* locus in *H. melpomene*.** (**A**) Genotypes at *optix* across all individuals shown as yellow for homozygotes for the highland (*H. m. plesseni*) allele, orange for heterozygotes and red for the lowland (*H. m. malleti*) allele. For clarity, only sites with at least 0.9 homozygote frequency difference between highland and lowland individuals are shown. The previously identified regulatory regions are shown on top (*55*, *85*). Individuals are grouped by the presence and absence of different red elements in the wings (dennis, ray, and red in the forewing bands, see also fig. S3C–F). The first three groups show individuals that are almost entirely homozygous for the lowland allele (yellow), heterozygous (orange) or homozygous for the highland allele (red) across the region, including the *ray* and *dennis* regulatory elements. Accordingly, they show red only in the forewing bands typical for the highland race (highland homozygotes, $optix^{H/H}$, yellow), red scales in the forewing bands and dennis–ray (heterozygotes, $optix^{H/L}$, orange), or only dennis–ray typical for the lowland race (lowland homozygotes, $optix^{L/L}$, red). Recombinant individuals that decouple the red scales, dennis and ray phenotypes are shown in the last two groups. The second last group includes individuals that feature lowland *malleti*-like dennis–ray pattern and lack red scales in the forewing band typical for the highland race. These individuals are homozygous for the lowland allele at the *dennis* and *ray* elements, consistent with their dennis–ray phenotype. However, they are heterozygous at *band*, which should predict red scales in the forewing band, if *band* is the element controlling the trait, as reported elsewhere (*55*, *85*). Instead, these individuals uniformly lack red scales. The only fully concordant region among these individuals is the *dennis* element. This shows that in the Ecuador hybrid zone, *dennis*, rather than *band*, controls the presence of red scales in the forewing bands. The last group consists of individuals that show a dennis patch and bar but no rays. These individuals are homozygous for the highland allele at *ray* (yellow), but heterozygous at *dennis.* Note here how the breakpoints just 5' to the *dennis* elements produces a precise match to the dennis-only phenotype. (**B**) Refinement of the minimal interval through a recombinant, CAM016820. This individual belongs to the middle heterozygous group (blue arrow in **A**, orange group): it has red scales in the forewing band and dennis–ray. It is heterozygous at *dennis* but contains a recombinant breakpoint within *ray* as inferred by (*85*). Since this individual has a normal ray pattern, we can rule out the 6 kbp segment homozygous for the highland allele. Thus, this recombinant individual refines the *ray* element from 14.1 kbp to 8.0 kbp (blue rectangle in **A**). Individual molecules are shown on the left (red lines indicate recombinant haplotypes transitioning from highland, yellow, to lowland, red alleles). Yellow and orange shading indicate the inferred genotypes. (**C**) Inferred regulatory elements controlling the major color pattern elements observed in this hybrid zone.

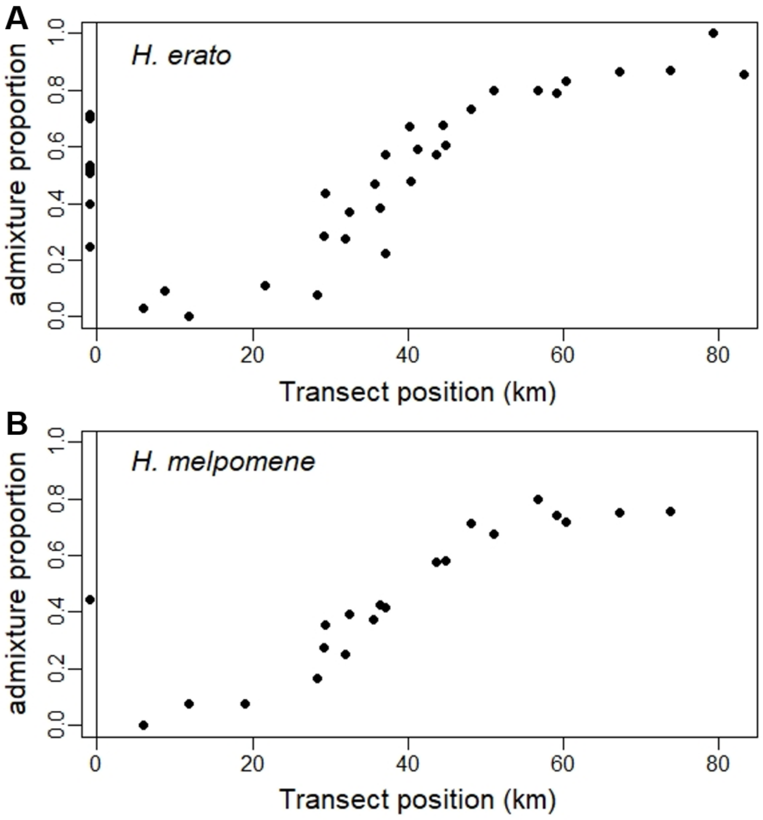

**Fig. S10. Genome-wide clines along the hybrid zone in both species.** Admixture proportions derived from `NGSadmix` (*65*) results with K=2 for *H. erato* and K=3 for *H. melpomene* averaged across all samples of the same collection site and plotted against transect position. Sampling sites outside of the transect are shown below 0. Genomic sites have been randomly subsampled to 10% of the sites to reduce linkage and increase run efficiency. The whole genome shows clinal variation along the transect zone.

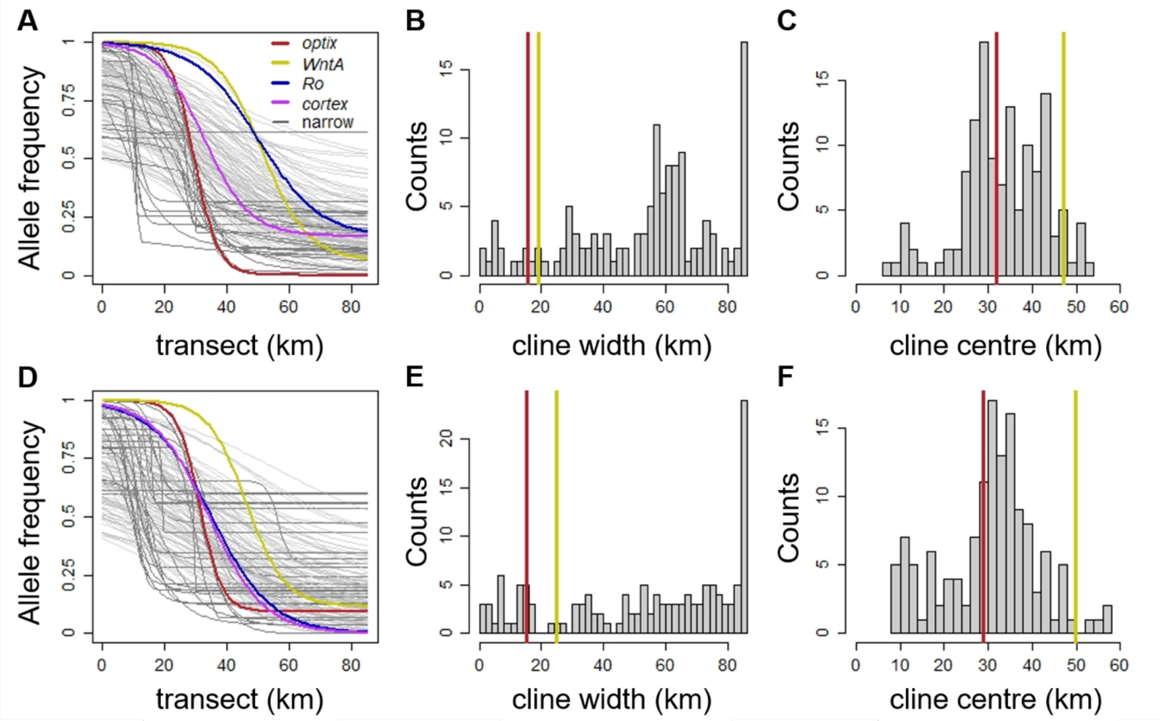

**Fig. S11. Single-site clines along the hybrid zone in both species.** Clines of the most highly differentiated position of each 100 kbp window in *H. erato* (**A-C**) and in *H. melpomene* (**D-F**), excluding sites with $F_{ST}$ below 0.5 in *H. erato* or below 0.4 in *H. melpomene*. Clines of the sites closest to the four color loci are highlighted in color and clines narrower than 30 km in darker grey. A list of the narrow clines is given in Tables S12–13. Histograms show the widths and centres of all clines, whereby the width and centre of the haplotype clines of *optix* and *WntA* are indicated with vertical lines in dark red and yellow, respectively. Note that the high number of narrow clines centred around 10 km represent SNPs where the population at El Topo in Baños (Tungurahua) is distinct from all other populations potentially due to isolation by distance.

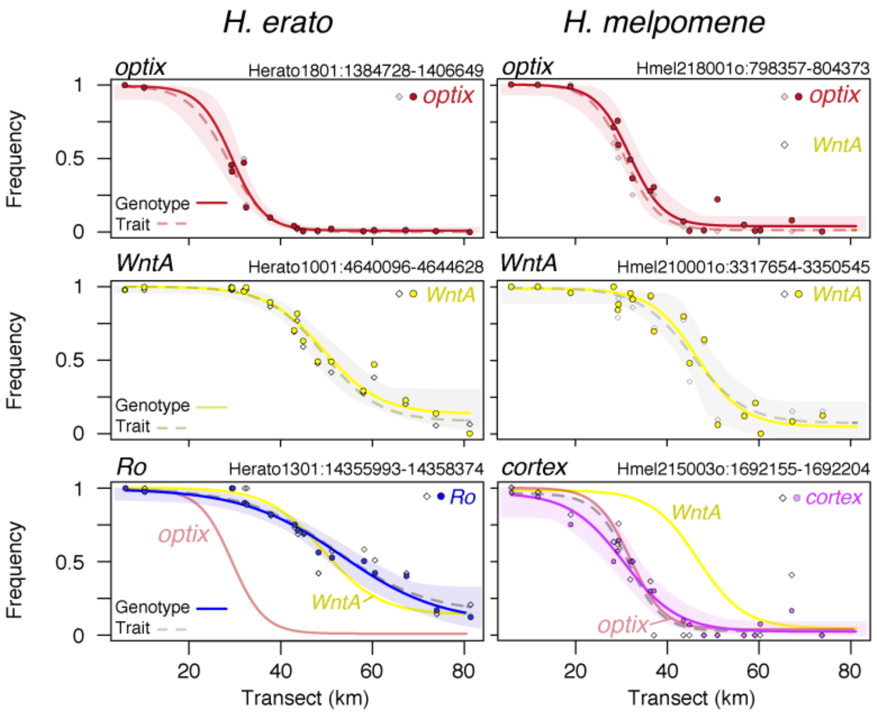

**Fig. S12. Haplotype frequency clines at the major color loci and the frequencies of morphs across the hybrid zones**. Clines at the major color loci *optix*, *WntA, Ro* (in *H. erato*, left column) and *cortex* (in *H. melpomene*, right column) across the hybrid zones are shown for haplotypes (genomic positions indicated above each panel; cline fits: solid lines with shading for confidence intervals; data points: circles) and phenotypes (cline fit: dashed lines; data points: diamonds). For the dominant loci *Ro* and *cortex/N*, the trait-based allele frequencies are estimated by assuming Hardy-Weinberg equilibrium. The haplotype cline fits for *optix* and *WntA* are repeated here to show the remarkable coincidence of these modifier loci with the major loci in each species.

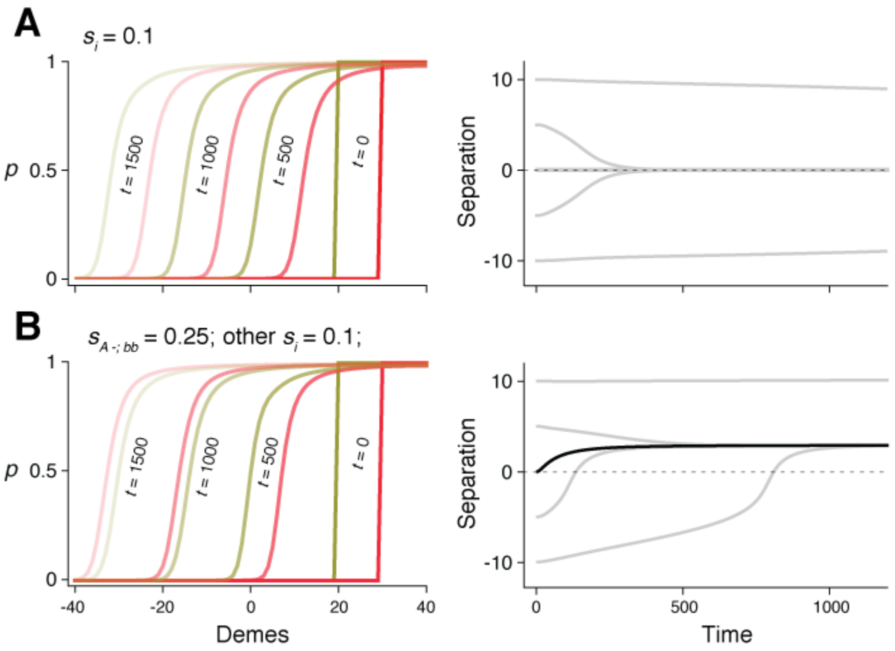

**Fig. S13. Clines at two loci, each with complete dominance for the lowland allele.** Each of the four morphs, labelled $i$, has fitness $1 + s_i (P_i - Q_i)$, where $P_i$ is the frequency of morph $i$. In the top row, selection is symmetric, so that $s_i = 0.1$ for all $i$. (**A**) Initially, there are step clines at each locus (red and yellow), centred at 20, 30. Clines move to the left, due to dominance, but remain ~10 apart; they are shown at 0, 500, 1000, 1500 generations, the two loci indicated by line thickness. Right: separation between clines over time, for different initial separations. If the clines are close enough, swamping and LD pulls them together, but otherwise, they remain separated. A density gradient or extrinsic selection gradient would pin the clines, and force them together even if initially well-separated. (**B**) The same, but with stronger frequency-dependence $s_{A-;bb}$ favoring one of the hybrid phenotypes. If the clines are initially displaced such that the less fit hybrid morph is common, the red cline moves faster, and crosses the yellow cline, reaching an equilibrium shift such that the fitter hybrid is commoner. However, if the clines are far enough apart, the fitter hybrid can be maintained indefinitely (bottom right, lowest line). Right: the black line shows a scenario, where even if the two clines start off coincident, they become displaced due to the fitness advantage of the fitter hybrid morph. The simulation uses nearest-neighbor migration, with $m$=0.5.

| **Platform (reference)** | 10X Genomics, Chromium (*7*) | Haplotagging; this study | TELL-seq (*87*) | Droplet barcode sequencing (*88*) | stLFR (*9*) | CPTv2-seq (*8*) |
|---|---|---|---|---|---|---|
| **Main technique** | Microdroplet compartments | Tn5 tranposase on barcoded beads | MuA transposase + barcoded bead capture | Commercial Tn5-flex + basic emulsion PCR | Tn5-transposase + barcoded bead capture | Tn5-transposase hybridized onto beads |
| **Status** | Commercial, discontinued | **Open protocol** | Commercial, not yet available | **Open protocol; components available** | Commercial, trial basis | Published protocol; not widely adopted |
| **Processing time** | Two days | **< 3 hours** | **< 3 hours** | 8 hours, 2 hours hands on | 5-6 hours | 1 tube method: <3.5 h 384-plate method: slow |
| **Number of steps** | Droplet encapsulation; blunting, A-tailing, adaptor ligation; PCR | **Tn5 transposition; PCR** | MuA transposition; hybridization and ligation; Tn5 tranposition; PCR | Tn5-flex-on-bead-transposition; emulsion PCR; bead enrichment; i7 barcoding | Tn5 tagmentation; hybridization and ligation; optional additional ligation; PCR | 1 tube method: Tn5 tagmentation; PCR |
| **Sequencing platform** | **Illumina TruSeq** | **Illumina TruSeq/Nextera** | **Illumina TruSeq/Nextera** | **Illumina TruSeq/Nextera** | BGI sequencer (Custom); Illumina TruSeq | Illumina (custom sequencing recipe and primers) |

| | | | | | | |
|---|---|---|---|---|---|---|
| **Barcode positioning** | Read 1 inline, 16nt+10nt UMI; i5 8nt for multiplexing | **i5/i7 – 13nt + 13nt with 149nt read 2; can work with 12nt + 13nt with 2x150** | i5/i7 – 18nt + 8nt, with 2x146 | **i5/i7 – 20nt + read 2, i7 8nt for multiplexing** | Custom: 3 x 10nt | Custom i5 and i7 barcodes with long splint sequences |
| **Custom sequencing primer/settings*** | **none** | **none** | none, but details lacking on whether MuA transposon remains in read 2 | **none** | Yes, unsupported | Yes, unsupported |
| **Costs** | high (> 250€ / sample) | **low (< 2€ / sample)** | **low** | medium, USD 19 | USD 28.53 | **1 tube method: low** |
| **Scalability** | - | **+ + + + 96-well plate format** | **+ + + 96-well plate format** | - | + + + ? | + + + ? |
| **Barcode diversity** | 1 – 4 M | **Up to 85 M** | **> 2 B** | >> 1 M | **3.6 B** | 147 K |

* Extended index cycles are fully supported by Illumina and is considered “standard” here
**Bold-faced** cells indicate the best option in terms of cost, handling, or analytical advantage

**Table S1. Linked-read sequencing technique comparisons.** K = thousands; M = millions; B = billions;

| Platform | | Illumina TruSeq (commercial provider) | Illumina Nextera/ Tn5 (in-house) | 10X Chromium (in-house) | Haplotagging |
|---|---|---|---|---|---|
| **Major cost items** | DNA extraction | 2.6€ | 0.53€ | 0.53€ | **0.53€** |
| | DNA normalization and size selection | | 0.26€ | 0.26€ | **0.26€** |
| | Library generation | 13.5€ | 0.73€ | 210.8€ | **0.73€** |
| **Total** | | 16.1€ | 1.52€ | 212€ | **1.52€** |

**Table S2. Example sequencing library preparation costs**

Listed above are representative consumables-only operating costs from a genome core facility for making sequencing-ready libraries from tissue biopsies, excluding one-time costs, which vary greatly across the library types. A key comparison here is that the sequencing library preparation costs for haplotagging is only a fraction of that of 10X Genomics's Chromium, yet it yields comparable results. For haplotagging, the major one-time costs are purchasing the oligonucleotides Data S1 and bead assembly, which for the current experiment cost around 6000€ for 85 million beadTags. However, a single such order will deliver enough oligonucleotides for >20,000 libraries, bringing the per-sample costs down to ~0.3€ per sample.

| Sample | n | Total reads (M) | Mapped Sequence (Gbp) | Fold-coverage (x) | Mean cov./ Sample (x) | Mean molecular coverage (x) |
|---|---|---|---|---|---|---|
| Mouse, F1(CASTxBL6) | 1 | 401.3 | 55.21 | 12.60 | 12.60 | 165.6 |
| Mouse, N2(CASTxBL6)* | 1 | 571.4 | 80.47 | 59.6 | 59.6 | 1232.3 |
| Human, GM12878 | 1 | 283.1 | 37.28 | 12.30 | 12.30 | 106.4 |
| Mouse, Longshanks F11, F16, F17† | 245 | 0.86 | 0.12 | 59.8 | 0.24 | 3.3 |
| *H. erato* | 484 | 3,846 | 431.8 | 952.8 | 1.54 | 11.5 |
| *H. melpomene* | 187 | 1,267 | 153.4 | 1569.0 | 2.77 | 19.6 |

* Only considering heterozygous regions

† Only considering the Chr3: 50–52 Mbp focal region.

**Table S3. Sequencing Summaries**

| Sample | NA12878[a] | GM12878 | F1(BL6xCAST) | F1(BL6xCAST) | N2(BL6xCAST)* |
|---|---|---|---|---|---|
| **Species** | Human | Human | Mouse | Mouse | Mouse |
| **Platform** | CPTv2, one-tube | Haplotagging | 10X Genomics | Haplotagging | Haplotagging |
| **Barcodes (M)** | 0.147 | 1.701 | 1.432 | 1.130 | 2.232 |
| **Read length** | 2 x 76 | 2 x 150 | 2 x 150 | 2 x 150 | 2 x 150 |
| **Number of read-pairs (millions)** | 648 | 141.56 | 61.61 | 200.66 | 285.70 |
| **Mapped bases (Gbp)/Mapped** | 73/75% | 37.28/88% | 16.05/87% | 55.21/92% | 80.47/94% |
| **Uniqueness / Duplicates** | 79%/21% | 41%/59% | 89%/11% | 62%/38% | 75%/25% |
| **Mean coverage (no duplicates)** | 19.2 | 12.30 | 5.14 | 12.60 | 59.64* |
| **Mean DNA per barcode[a]** | 6 | 1.58 | 6.00 | 2.44 | 1.05 |
| **Mean reads per molecule [b]** | 5 | 10.11 | 6.23 | 6.51 | 6.71 |
| **Mean molecular coverage [c] (duplicates removed)** | n.d. | 106.36 | 168.44 | 165.55 | 1232.30 |
| **$N_{50}$[d]/max. molecule size** | 34.9/339 | 63.47/573 | 82.68/1010 | 42.05/415 | 40.87/281 |
| **Informative linked reads[e] $N_{50}$ (kbp)** | 58.5 | 41.79 | 87.72 | 44.63 | 55.98 |
| **heterozygous SNPs phased** | 98% | 98.59% | 99.80% | 99.74% | 99.91% |
| **Phasing block [f] $N_{50}$ (Mbp)** | 1.14 | 1.08 | 29.42 | 20.01 | 14.45 |
| **Longest phasing block (Mbp)** | 3.46 | 6.83 | 87.30 | 61.46 | 58.72* |
| **Short switch error rate [g]** | 0.13% | 0.95% | 0.0011% | 0.18% | 0.075% |
| **Long switch error rate[h]** | 0.0085% | 0.039% | 0.022% | 0.026% | 0.014% |

* Only considering heterozygous regions

a: the number of loci in the genome for a given beadTag

b: molecules are defined as sets of reads sharing the same beadTag within 50 kbp from each other

c: coverage calculated using overlap by molecules

d: $N_{50}$: the shortest fragment among those that make up 50% of the combined length of all fragments

e: LR that overlap heterozygous SNP positions

f: as generated by HAPCUT2 with PHRED-scaled threshold value of 30

g: Incorrectly phased SNPs in a phased block, according to reference-quality data

h: Molecules that show runs of alleles from both haplotypes. These are erroneous molecules arising from actual recombinant molecules or molecular artefacts, or barcode re-use

**Table S4. Phasing performance in single individuals and comparisons with other platforms.**

| Site No. | Site name | Lat (S) | Lon (W) | Elevation (m) | Distance along transect (km) |
|---|---|---|---|---|---|
| | Transect | | | | |
| 1 | Cashaurco | 1° 25.345' | 78° 10.376' | 1252 | 4.98 |
| 2 | Parroquia Cumandá | 1° 27.250' | 78° 08.951' | 1251 | 5.17 |
| 3 | El Topo | 1° 23.881' | 78° 10.688' | 1307 | 6.02 |
| 4 | Mangayacu | 1° 26.227' | 78° 07.373' | 1239 | 8.65 |
| 5 | Pindo-Mirador | 1° 27.598' | 78° 04.368' | 1227 | 11.82 |
| 6 | Fátima 2 | 1° 25.552' | 78° 01.092' | 1058 | 18.97 |
| 7 | Fátima-JatunPaccha km 1.6 | 1° 25.755' | 77° 59.199' | 991 | 21.65 |
| 8 | Colonia SimónBolivar | 1° 22.049' | 78° 00.961' | 1000 | 22.81 |
| 9 | Fátima-JatunPaccha km 7 | 1° 25.422' | 77° 56.863' | 1009 | 25.57 |
| 10 | km 19 Col Llandia km 1.2 | 1° 21.737' | 77° 57.566' | 1041 | 28.34 |
| 11 | Tnt Hugo Ortiz km 2 | 1° 22.981' | 77° 56.674' | 1036 | 28.40 |
| 12 | km 19 Col Llandia 2 | 1° 20.800' | 77° 57.607' | 995 | 29.25 |
| 13 | Fátima-JatunPaccha km 11.7 | 1° 24.596' | 77° 54.921' | 988 | 29.41 |
| 14 | km 18 Com SanPablo | 1° 23.367' | 77° 54.062' | 955 | 32.00 |
| 15 | km 25 Llandia | 1° 19.999' | 77° 56.043' | 953 | 32.48 |
| 16 | Colonia Mariscal I | 1° 22.276' | 77° 52.471' | 961 | 35.57 |
| 17 | Colonia 4 de Agosto | 1° 14.448' | 77° 57.656' | 939 | 35.79 |
| 18 | km 37 San Fco. de Llandia km 3.3 | 1° 18.374' | 77° 54.590' | 839 | 36.39 |
| 19 | Vía Colonia 4 de Agosto km 8.2 | 1° 14.779' | 77° 56.564' | 750 | 37.11 |
| 20 | Colonia Mariscal II | 1° 22.270' | 77° 51.446' | 947 | 37.14 |
| 21 | Vía Colonia 4 de Agosto 1 | 1° 14.736' | 77° 54.514' | 638 | 40.30 |
| 22 | km 35 Antena | 1° 17.462' | 77° 52.642' | 1064 | 40.32 |
| 23 | km 31 Comuna Cajabamba | 1° 20.290' | 77° 50.123' | 954 | 41.23 |
| 24 | km 35 Sendero Pta. San Cristobal | 1° 17.446' | 77° 50.514' | 947 | 43.59 |
| 25 | km 31 bajada río Arajuno | 1° 19.309' | 77° 48.589' | 908 | 44.60 |
| 26 | km 43 Sendero Colonia SanVicente | 1° 14.597' | 77° 51.605' | 845 | 44.89 |
| 27 | San Pedro de Puní | 1° 15.111' | 77° 49.176' | 931 | 48.07 |
| 28 | El Capricho km 5.7 | 1° 11.269' | 77° 49.863' | 824 | 51.02 |
| 29 | LuzDeAmérica km 11.5 | 1° 10.101' | 77° 46.867' | 665 | 56.82 |
| 30 | Vía Arajuno km 58.4 | 1° 15.058' | 77° 41.931' | 606 | 59.22 |
| 31 | Apuya km 2.7 | 1° 06.934' | 77° 46.700' | 579 | 60.38 |

| | | | | | |
|---|---|---|---|---|---|
| 32 | Colonia 20 de Enero | 1º 05.494' | 77º 43.197' | 459 | 67.24 |
| 33 | Y de Misahuallí | 1º 03.683' | 77º 40.106' | 440 | 73.85 |
| 34 | San Pedro de Arajuno | 1º 05.928' | 77º 35.067' | 376 | 79.23 |
| 35 | Río Pusuno | 1º 00.940' | 77º 35.849' | 378 | 83.22 |
| | Not transect | | | | |
| 36 | Río Yanamacas Chico | 1º 47.584' | 78º 01.114' | 1054 | |
| 37 | Sangay | 1º 46.789' | 77º 58.923' | 992 | |
| 38 | Vía Arapicos km 4.5 | 1º 49.057' | 77º 57.608' | 935 | |
| 39 | Antenas del Calvario | 1º 30.697' | 77º 54.684' | 1056 | |
| 40 | Vía Colonia Juan de Velasco km 1 | 1º 27.554' | 77º 53.202' | 994 | |
| 41 | Sendero Col Juan de Velasco | 1º 28.519' | 77º 52.220' | 856 | |
| 42 | Y de Taculín | 1º 30.191' | 77º 50.255' | 612 | |
| 43 | Vía Canelos km 6.3 | 1º 35.812' | 77º 49.330' | 698 | |
| 44 | Río Camagua?/Chontoa | 1º 34.908' | 77º 49.259' | 638 | |
| 45 | Río Llushcayacu | 1º 24.128' | 77º 47.843' | 1034 | |
| 46 | Vía Triunfo-Villano-Parahua | 1º 24.963' | 77º 43.741' | 857 | |
| 47 | Sendero vía Arajuno km 37 | 1º 22.667' | 77º 42.635' | 1114 | |

**Table S5. Sampling locations**

**Table S6.** ***H. erato*** **individuals by site and genotype. Shaded in grey are the two parental forms and the double-banded-hybrid.**

| Site No. | Elevation (m) | *optix*$^{H/H}$ | *optix*$^{H/L}$ | | | | *optix*$^{L/L}$ | | | | | | |
|---|---|---|---|---|---|---|---|---|---|---|---|---|---|
| | | *WntA*$^{H/H}$ | *WntA*$^{H/H}$ | | *WntA*$^{H/L}$ | *WntA*$^{L/L}$ | *WntA*$^{H/H}$ | | *WntA*$^{H/L}$ | | | *WntA*$^{L/L}$ | |
| | | *Ro*$^{H/-}$ | *Ro*$^{H/-}$ | *Ro*$^{L/L}$ | *Ro*$^{H/-}$ | *Ro*$^{H/-}$ | *Ro*$^{H/-}$ | *Ro*$^{L/L}$ | *Ro*$^{H/-}$ | *Ro*$^{L/L}$ | ? | *Ro*$^{H/-}$ | *Ro*$^{L/L}$ |
| 1 | 1252 | 11 | | | | | | | | | | | |
| 2 | 1251 | 3 | | | | | | | | | | | |
| 3 | 1307 | 12 | | | | | | | | | | | |
| 4 | 1239 | 61 | | | | | | | | | | | |
| 5 | 1227 | 41 | 1 | | | | | | | | | | |
| 6 | 1058 | 48 | | | | | | | | | | | |
| 7 | 991 | 19 | 2 | | | | | | | | | | |
| 8 | 1000 | 32 | | | | | | | | | | | |
| 9 | 1009 | | | | | | | | | | | | |
| 10 | 1041 | 29 | 1 | | | | | | | | | | |
| 11 | 1036 | 7 | | | | | | | | | | | |
| 12 | 995 | 31 | 6 | | | | 1 | | | | | | |
| 13 | 988 | 9 | 5 | | | | 1 | | | | | | |
| 14 | 955 | 10 | 10 | | | | | | | | | | |
| 15 | 953 | 15 | 8 | | | | 13 | | | | | | |
| 16 | 961 | | | | | | | | | | | | |
| 17 | 939 | | 3 | | | | 5 | | | | | 1 | |
| 18 | 839 | 2 | 14 | | 1 | | 52 | 1 | 8 | | | | |
| 19 | 750 | | 2 | | | | 4 | | | | | | |
| 20 | 947 | | 1 | | | | 1 | | 1 | | | | |
| 21 | 638 | | | | 1 | | 2 | | | | | | |
| 22 | 1064 | | 3 | 1 | 3 | | 58 | 2 | 17 | | 1 | 2 | |
| 23 | 954 | | | | 1 | | 2 | | 1 | | | | |
| 24 | 947 | | 2 | | | | 20 | 1 | 8 | 1 | | 2 | 1 |
| 25 | 908 | | | | | | 5 | | 5 | 1 | | | |
| 26 | 845 | | | | | | 9 | 2 | 15 | 1 | | 5 | 1 |
| 27 | 931 | | | | | | 2 | 2 | 4 | 3 | | 4 | |
| 28 | 824 | | | | | 1 | 6 | | 12 | 6 | | 9 | 8 |
| 29 | 665 | | | | | | | 1 | 7 | 4 | | 11 | 7 |
| 30 | 606 | | | | | | | | 3 | | | 3 | 1 |
| 31 | 579 | | | | | | 1 | 1 | 10 | 9 | | 17 | 28 |
| 32 | 459 | | | | | | | | 1 | 5 | | 9 | 18 |
| 33 | 440 | | | | | | | | 1 | 3 | | 9 | 24 |
| 34 | 376 | | | | | | | | | 2 | | 10 | 37 |
| 35 | 378 | | | | | | | | | | | | 4 |

| TOTAL | | 330 | 58 | 1 | 6 | 1 | 182 | 10 | 93 | 35 | 1 | 82 | 129 |
|---|---|---|---|---|---|---|---|---|---|---|---|---|---|
| Site No. | Elevation (m) | *optix*$^{H/H}$ *WntA*$^{H/H}$ *Ro*$^{H/-}$ | *optix*$^{H/L}$ *WntA*$^{H/H}$ *Ro*$^{H/-}$ | *Ro*$^{L/L}$ | *WntA*$^{H/L}$ *Ro*$^{H/-}$ | *WntA*$^{L/L}$ *Ro*$^{H/-}$ | *optix*$^{L/L}$ *WntA*$^{H/H}$ *Ro*$^{H/-}$ | *Ro*$^{L/L}$ | *WntA*$^{H/L}$ *Ro*$^{H/-}$ | *Ro*$^{L/L}$ | ? | *WntA*$^{L/L}$ *Ro*$^{H/-}$ | *Ro*$^{L/L}$ |
| *36* | | *1* | | | | | | | | | | | |
| *37* | | *5* | | | | | | | | | | | |
| *38* | | | | | | | | | | | | | |
| *39* | | *3* | *3* | | | | | | *1* | | | | |
| *40* | | *1* | *1* | | | | *3* | | | | | | |
| *41* | | | *2* | | | | *3* | | | | | | |
| *42* | | | *1* | *1* | *1* | | *2* | | *1* | | | | |
| *43* | | | | | | | | | | | | | |
| *44* | | | *2* | | *1* | *1* | | | *2* | | | | |
| *45* | | | | | | | *7* | | | | | | |
| *46* | | | | | | | | | *1* | | | | |
| *47* | | | | | | | | *1* | *1* | *1* | | *1* | |

**Table S7. *H. melpomene* individuals by site and genotype. Shaded in grey are the two parental forms and the double-banded-hybrid.**

| Site No. | Elevation (m) | *optix*$^{H/H}$ *WntA*$^{H/H}$ $N^{H/H}$ | $N^{-/L}$ | *WntA*$^{H/L}$ $N^{H/H}$ | $N^{-/L}$ | *optix*$^{H/L}$ *WntA*$^{H/H}$ $N^{H/H}$ | $N^{-/L}$ | *WntA*$^{H/L}$ $N^{H/H}$ | $N^{-/L}$ | *WntA*$^{L/L}$ $N^{-/L}$ | *optix*$^{-/L}$ *WntA*$^{H/H}$ $N^{-/L}$ | *optix*$^{L/L}$ *WntA*$^{H/H}$ $N^{H/H}$ | $N^{-/L}$ | ? | *WntA*$^{H/L}$ $N^{H/H}$ | $N^{-/L}$ | *WntA*$^{L/L}$ $N^{-/L}$ | ? ? | *optix uncertain* *WntA*$^{H/H}$ $N^{-/L}$ | *WntA*$^{H/L}$ $N^{H/H}$ |
|---|---|---|---|---|---|---|---|---|---|---|---|---|---|---|---|---|---|---|---|---|
| 1 | 1252 | 39 | | | | | | | | | | | | | | | | | | |
| 2 | 1251 | 1 | | | | | | | | | | | | | | | | | | |
| 3 | 1307 | 19 | | | | | | | | | | | | | | | | | | |
| 4 | 1239 | 16 | | | | | | | | | | | | | | | | | | |
| 5 | 1227 | 32 | 1 | 1 | | | | | | | | | | | | | | | | |
| 6 | 1058 | 8 | 4 | 1 | | | | | | | | | | | | | | | | |
| 7 | 991 | 1 | 1 | | | 1 | | | | | | | | | | | | | | |
| 8 | 1000 | 3 | | | | | | | 1 | | | | | | | | | | | |
| 9 | 1009 | 1 | | | | | | | | | | | | | | | | | | |
| 10 | 1041 | 1 | 2 | | | 1 | 1 | | | | | | 1 | | | | | | | |
| 11 | 1036 | | | | | | | | | | | | | | | | | | | |
| 12 | 995 | 2 | | | | 1 | 1 | | 2 | | | | | | | 1 | 1 | | 1 | |
| 13 | 988 | 1 | 1 | | | 2 | 3 | | | | | | 1 | | | | | | | 1 |
| 14 | 955 | | 2 | | | | 4 | 1 | | | | | 1 | | | | | | | |
| 15 | 953 | | 2 | | | | 1 | | | | | 1 | 3 | | 1 | 1 | 2 | | | |
| 16 | 961 | | | | 2 | | | | | | | | 1 | | | | | | 1 | |
| 17 | 939 | | | | | | | | | | | | 1 | | | 1 | 2 | | | |
| 18 | 839 | | 2 | | | 1 | 6 | | | | | 1 | 7 | 2 | | 3 | | | | |
| 19 | 750 | | 1 | | | | 2 | | | | | | | | | | | | | |
| 20 | 947 | | | | | | 3 | | | | | | | | | 3 | 1 | | | |
| 21 | 638 | | | | | | | | | | | | 1 | | | | | | | |
| 22 | 1064 | | | | | | 1 | | | | | | | | | | | | | |
| 23 | 954 | | | | | | | | | | | | 1 | | | | 1 | | | |
| 24 | 947 | | | | | | 1 | | | | | | 10 | | | 4 | 8 | | | |
| 25 | 908 | | | | | | | | | | | | 1 | | | 1 | | | | |
| 26 | 845 | | | | | | | | | | | | 1 | | | 4 | 2 | | | |
| 27 | 931 | | | | | | | | | | | | 3 | | | 2 | 2 | | | |
| 28 | 824 | | | | | | | | | | | | | | | 2 | 29 | | | |
| 29 | 665 | | | | | | | | 1 | | | | | | | 9 | 15 | | | |
| 30 | 606 | | | | | | | | | | | | 1 | | | 1 | 10 | | | |
| 31 | 579 | | | | | | | | | | | | 1 | | | 1 | 47 | | | |
| 32 | 459 | | | | | | | | | | | | | | 1 | 2 | 5 | | | |
| 33 | 440 | | | | | | | | | | | | | | | 8 | 24 | | | |
| 34 | 376 | | | | | | | | | | | | 1 | | | 1 | 19 | 5 | | |
| 35 | 378 | | | | | | | | | | | | | | | | 1 | | | |

| | | | | | | | | | | | | | | | | | | | | |
|---|---|---|---|---|---|---|---|---|---|---|---|---|---|---|---|---|---|---|---|---|
| TOTAL | | 124 | 16 | 2 | 2 | 6 | 23 | 1 | 4 | 0 | 0 | 2 | 35 | 2 | 2 | 44 | 169 | 5 | 2 | 1 |
| Site No. | Elevation (m) | *optix*$^{H/H}$ | | | | *optix*$^{H/L}$ | | | | | *optix*$^{-/L}$ | *optix*$^{L/L}$ | | | | | | | *optix uncertain* | |
| | | *WntA*$^{H/H}$ | | *WntA*$^{H/L}$ | | *WntA*$^{H/H}$ | | *WntA*$^{H/L}$ | | *WntA*$^{L/L}$ | *WntA*$^{H/H}$ | *WntA*$^{H/H}$ | | | *WntA*$^{H/L}$ | | *WntA*$^{L/L}$ | ? | *WntA*$^{H/H}$ | *WntA*$^{H/L}$ |
| | | *N*$^{H/H}$ | *N*$^{-/L}$ | *N*$^{H/H}$ | *N*$^{-/L}$ | *N*$^{H/H}$ | *N*$^{-/L}$ | *N*$^{H/H}$ | *N*$^{-/L}$ | *N*$^{-/L}$ | *N*$^{-/L}$ | *N*$^{H/H}$ | *N*$^{-/L}$ | ? | *N*$^{H/H}$ | *N*$^{-/L}$ | *N*$^{-/L}$ | ? | *N*$^{-/L}$ | *N*$^{H/H}$ |
| *36* | | *1* | *1* | | | *1* | | | | | | | | | | | | | | |
| *37* | | | *3* | | | | | | | | | | | | | | | | | |
| *38* | | | | | | | *2* | | *1* | | | | | | | | *1* | | | |
| *39* | | | *2* | | | | *1* | | | | | | *1* | | | | | | | |
| *40* | | *1* | *1* | | | | *1* | | | | | | *2* | | | *1* | | | | |
| *41* | | *1* | | | | | *3* | | | | | *1* | *8* | | | *1* | | | | |
| *42* | | | | | | | | | | *1* | | | | | | *1* | | | | |
| *43* | | | | | | | | | | | | | | | | *1* | | | | |
| *44* | | | *1* | | | | | | | *1* | *1* | | | | *2* | | *1* | | | |
| *45* | | | | | | | *2* | | | | | | *4* | | | *2* | *3* | | | |
| *46* | | | | | | | | | | | | | | | | | *1* | | | |
| *47* | | | | | | | | | | | | | | | | | | | | |

The *N* locus corresponds to the gene *cortex*.

***H. erato***

| Herato1801: 1399223: | | *optix* - *dennis* & *ray* HH | HL | LL |
|---|---|---|---|---|
| Genotype | 0\|0 | 0 | 9 | 370 |
| | 0\|1 | 0 | 18 | 0 |
| | 1\|0 | 0 | 13 | 1 |
| | 0/1 | 0 | 28 | 2 |
| | 1\|1 | 33 | 4 | 0 |

Sensitivity (high-confidence): **94.0%**
Accuracy (high-confidence): **98.4%**
Accuracy (all): **96.7%**

| Herato1001: 4640866: | | *WntA* - band HH | HL | LL |
|---|---|---|---|---|
| Genotype | 0\|0 | 210 | 8 | 2 |
| | 0\|1 | 3 | 19 | 1 |
| | 1\|0 | 1 | 21 | 3 |
| | 0/1 | 2 | 62 | 7 |
| | 1\|1 | 0 | 6 | 79 |

Sensitivity (high-confidence): **88.2%**
Accuracy (high-confidence): **95.5%**
Accuracy (all): **96.4%**

***H. melpomene***

| Hmel218003o: 800708: | | *optix* - *dennis* & *ray* HH | HL | LL |
|---|---|---|---|---|
| Genotype | **0l0** | 96 | 1 | 0 |
| | **0l1** | 5 | 10 | 0 |
| | **1l0** | 3 | 10 | 0 |
| | **0/1** | 1 | 3 | 0 |
| | **1l1** | 0 | 2 | 53 |

Sensitivity (high-confidence): **94.0%**
Accuracy (high-confidence): **95.4%**
Accuracy (all): **93.5%**

| Hmel210001o: 3333301: | | *WntA* - *band* HH | HL | LL |
|---|---|---|---|---|
| Genotype | **0l0** | 39 | 6 | 0 |
| | **0l1** | 0 | 18 | 0 |
| | **1l0** | 1 | 16 | 1 |
| | **0/1** | 0 | 1 | 0 |
| | **1l1** | 0 | 3 | 102 |

Sensitivity (high-confidence): **98.9%**
Accuracy (high-confidence): **94.1%**
Accuracy (all): **94.1%**

**Table S8. Validation results for STITCH at known color loci**

| *H. e. notabilis*, ω | | | | | |
|---|---|---|---|---|---|
| **Rank** | **Chrom** | **Start** | **End** | **$\omega_{MAX}$** | **Remarks** |
| 1 | Herato1202 | 6484876 | 6485047 | 8129.4 | |
| 2 | Herato1801 | 1373655 | 1374171 | 4014.6 | *Optix* |
| 3 | Herato1301 | 14347538 | 14348248 | 4001.7 | *Ro* |
| 4 | Herato0209 | 138991 | 139670 | 3452.2 | |
| 5 | Herato1411 | 5829303 | 5829391 | 2829.4 | |
| 6 | Herato1505 | 2503334 | 2503860 | 2699.0 | *cortex* |
| 7 | Herato1001 | 4674256 | 4674542 | 2362.7 | *WntA* |
| 8 | Herato1904 | 325896 | 326010 | 1973.0 | |
| 9 | Herato1708 | 339041 | 339341 | 1817.5 | |
| 10 | Herato1005 | 908198 | 908330 | 1785.6 | |
| 11 | Herato1805 | 1019516 | 1019641 | 1744.5 | |
| 12 | Herato0310 | 8188969 | 8188997 | 1587.2 | |
| 13 | Herato0206 | 537017 | 537495 | 1580.7 | Taste receptor |
| 14 | Herato1108 | 5033276 | 5033409 | 1528.4 | |
| 15 | Herato0204 | 454067 | 454293 | 1397.8 | Chr2 inversion |
| 16 | Herato0701 | 5854241 | 5854327 | 1343.2 | |
| 17 | Herato2101 | 870019 | 870184 | 1336.0 | |
| 18 | Herato1807 | 1708149 | 1708271 | 1281.1 | |
| 19 | Herato1007 | 50780 | 50844 | 1244.5 | |
| 20 | Herato0606 | 8021793 | 8021826 | 1204.3 | |
| ***H. e. lativitta*, ω** | | | | | |
| 1 | Herato1003 | 1040018 | 1040285 | 3022.6 | |
| 2 | Herato2101 | 15799095 | 15799327 | 3022.1 | |
| 3 | Herato1701 | 9389072 | 9389597 | 2633.3 | |
| 4 | Herato0601 | 161075 | 161287 | 2117.1 | |
| 5 | Herato1108 | 2531390 | 2531499 | 2116.6 | |
| 6 | Herato0215 | 2311254 | 2311353 | 2066.5 | |
| 7 | Herato1202 | 12880941 | 12881215 | 1929.5 | |
| 8 | Herato0606 | 4570928 | 4571083 | 1864.1 | |
| 9 | Herato1301 | 9903737 | 9903883 | 1513.8 | |
| 10 | Herato0701 | 15271229 | 15271267 | 1451.6 | |
| 11 | Herato1908 | 1150364 | 1150752 | 1436.2 | |
| 12 | Herato0609 | 1403482 | 1403636 | 1416.1 | |
| 13 | Herato0101 | 5372303 | 5372329 | 1388.7 | |
| 14 | Herato2001 | 1896087 | 1896128 | 1252.7 | |
| 15 | Herato1007 | 5579873 | 5580179 | 1121.9 | |
| 16 | Herato1904 | 1936141 | 1936204 | 1117.4 | |
| 17 | Herato1910 | 1116041 | 1116286 | 1111.0 | |
| 18 | Herato0501 | 365815 | 366044 | 1102.5 | |
| 19 | Herato0901 | 11459293 | 11459394 | 1054.6 | |
| 20 | Herato1805 | 3911307 | 3911329 | 976.1 | |

***H. e. notabilis,* SweeD**

| Rank | Chrom | Window | CLR | Alpha | Remarks |
|---|---|---|---|---|---|
| 1 | Herato1301 | 14340000 | 406.9 | 5.50E-05 | *Ro* |
| 2 | Herato1801 | 1410000 | 380.9 | 2.99E-05 | *optix* |
| 3 | Herato1505 | 2100000 | 330.1 | 7.69E-05 | *cortex* |
| 4 | Herato0601 | 1900000 | 218.3 | 1.93E-04 | |
| 5 | Herato0209 | 210000 | 196.0 | 1.12E-04 | |
| 6 | Herato1005 | 4020000 | 153.7 | 1.84E-04 | |
| 7 | Herato2101 | 6570000 | 144.9 | 1.59E-04 | |
| 8 | Herato0701 | 4370000 | 133.0 | 2.20E-04 | |
| 9 | Herato1001 | 4630000 | 129.1 | 1.63E-04 | *WntA* |
| 10 | Herato1904 | 6270000 | 91.1 | 3.70E-04 | |
| 11 | Herato1801 | 3980000 | 88.4 | 6.82E-04 | |
| 12 | Herato0503 | 260000 | 69.6 | 5.19E-04 | |
| 13 | Herato0503 | 7440000 | 69.4 | 1.24E-03 | |
| 14 | Herato1202 | 12050000 | 67.4 | 3.94E-04 | |
| 15 | Herato0310 | 8970000 | 64.7 | 9.03E-04 | |
| 16 | Herato0801 | 4100000 | 64.4 | 2.57E-04 | |
| 17 | Herato0411 | 1830000 | 62.7 | 4.97E-04 | |
| 18 | Herato1003 | 1120000 | 58.3 | 1.07E-03 | |
| 19 | Herato1411 | 1730000 | 52.2 | 4.74E-04 | |
| 20 | Herato1202 | 5640000 | 51.7 | 8.08E-04 | |
| ***H. e. lativitta,* SweeD** | | | | | |
| 1 | Herato1505 | 2470000 | 208.6 | 4.41E-05 | *Cortex* |
| 2 | Herato2101 | 6570000 | 199.5 | 1.61E-04 | |
| 3 | Herato1701 | 9370000 | 166.8 | 2.22E-04 | |
| 4 | Herato1007 | 2910000 | 165.9 | 1.79E-04 | |
| 5 | Herato1001 | 5460000 | 161.9 | 1.57E-04 | *WntA* |
| 6 | Herato2101 | 800000 | 123.0 | 2.80E-04 | |
| 7 | Herato0701 | 4370000 | 114.0 | 2.53E-04 | |
| 8 | Herato1202 | 12050000 | 95.0 | 3.88E-04 | |
| 9 | Herato0801 | 6590000 | 87.3 | 8.30E-04 | |
| 10 | Herato1301 | 10160000 | 79.7 | 3.91E-04 | |
| 11 | Herato0206 | 480000 | 76.8 | 4.53E-04 | |
| 12 | Herato0503 | 2090000 | 70.3 | 2.65E-04 | |
| 13 | Herato1411 | 6410000 | 64.0 | 4.32E-04 | |
| 14 | Herato0310 | 8970000 | 60.3 | 1.08E-03 | |
| 15 | Herato2101 | 13530000 | 58.7 | 4.71E-04 | |
| 16 | Herato1108 | 4550000 | 53.6 | 5.39E-04 | |
| 17 | Herato0411 | 1830000 | 53.1 | 5.27E-04 | |
| 18 | Herato1708 | 2250000 | 52.4 | 6.44E-04 | |
| 19 | Herato2001 | 10610000 | 51.6 | 8.00E-04 | |
| 20 | Herato1801 | 1370000 | 51.5 | 2.38E-04 | *optix* |

***H. e. notabilis,* |iHS|**

| Rank | Chrom | Window | norm. iHS | Remarks |
|---|---|---|---|---|
| 1 | Herato1505 | 2480000 | 0.973684 | *Cortex* |
| 2 | Herato1801 | 580000 | 0.969136 | *optix* |
| 3 | Herato1910 | 5240000 | 0.968354 | |
| 4 | Herato1411 | 1690000 | 0.909871 | |
| 5 | Herato0101 | 15690000 | 0.880478 | |
| 6 | Herato0206 | 510000 | 0.878412 | |
| 7 | Herato0701 | 11640000 | 0.873118 | |
| 8 | Herato2101 | 10210000 | 0.862319 | |
| 9 | Herato1003 | 1080000 | 0.847291 | |
| 10 | Herato1605 | 2980000 | 0.847222 | |
| 11 | Herato1301 | 14820000 | 0.814126 | *Ro* |
| 12 | Herato1904 | 6790000 | 0.803957 | |
| 13 | Herato1701 | 9400000 | 0.790614 | |
| 14 | Herato0701 | 18890000 | 0.788095 | |
| 15 | Herato1708 | 1970000 | 0.785714 | |
| 16 | Herato1905 | 140000 | 0.75 | |
| 17 | Herato0209 | 150000 | 0.72549 | |
| 18 | Herato2101 | 11560000 | 0.70297 | |
| 19 | Herato1904 | 2830000 | 0.689482 | |
| 20 | Herato1002 | 110000 | 0.687386 | |

***H. e. lativitta,* |iHS|**

| | | | | |
|---|---|---|---|---|
| 1 | Herato1505 | 2480000 | 0.969697 | *Cortex* |
| 2 | Herato0206 | 510000 | 0.930591 | |
| 3 | Herato1805 | 3220000 | 0.819767 | |
| 4 | Herato0701 | 3790000 | 0.792982 | |
| 5 | Herato1701 | 9400000 | 0.786477 | |
| 6 | Herato1605 | 2980000 | 0.77381 | |
| 7 | Herato1003 | 1030000 | 0.762443 | |
| 8 | Herato1005 | 4020000 | 0.680488 | |
| 9 | Herato2101 | 6610000 | 0.675 | |
| 10 | Herato1007 | 2920000 | 0.668142 | |
| 11 | Herato1708 | 1950000 | 0.666667 | |
| 12 | Herato1901 | 2640000 | 0.631579 | |
| 13 | Herato2101 | 9560000 | 0.616541 | |
| 14 | Herato2101 | 800000 | 0.60217 | |
| 15 | Herato1701 | 11420000 | 0.588816 | |
| 16 | Herato2101 | 2610000 | 0.571429 | |
| 17 | Herato1301 | 13020000 | 0.56314 | |
| 18 | Herato0701 | 1240000 | 0.561497 | |
| 19 | Herato1001 | 4650000 | 0.547672 | *optix* |
| 20 | Herato0503 | 7570000 | 0.546816 | |

**Table S9. Top loci based on ω, SweeD and iHS in *H. erato.***

The top 20 loci found across the genome for the various selection statistics. Windows that are within 10 kbp of scaffold boundaries are excluded to avoid edge artifact.

***H. m. plesseni*, ω**

| Rank | Chrom | Start | End | $\omega_{MAX}$ | Remarks |
|---|---|---|---|---|---|
| 1 | Hmel210001o | 3342425 | 3342590 | 3723.7 | *WntA* |
| 2 | Hmel218003o | 783780 | 784041 | 3723.2 | *optix* |
| 3 | Hmel209001o | 1025345 | 1025407 | 3719.6 | |
| 4 | Hmel212001o | 12834990 | 12835662 | 3719.6 | |
| 5 | Hmel220003o | 8951615 | 8951717 | 3719.2 | |
| 6 | Hmel217001o | 4641742 | 4641788 | 3718.7 | |
| 7 | Hmel215003o | 1551143 | 1551242 | 3717.0 | *cortex* |
| 8 | Hmel208001o | 5625952 | 5626052 | 3715.9 | |
| 9 | Hmel201001o | 12103949 | 12104458 | 3451.7 | |
| 10 | Hmel211001o | 4378073 | 4378573 | 3124.6 | |
| 11 | Hmel207001o | 3725393 | 3725456 | 3121.5 | |
| 12 | Hmel213001o | 2053637 | 2053722 | 2975.1 | |
| 13 | Hmel202001o | 4028410 | 4028800 | 2701.1 | |
| 14 | Hmel203003o | 3301855 | 3301931 | 2613.3 | |
| 15 | Hmel206001o | 11682020 | 11682247 | 2515.4 | |
| 16 | Hmel219001o | 16075252 | 16075617 | 2094.3 | |
| 17 | Hmel205001o | 9861849 | 9862146 | 1937.9 | |
| 18 | Hmel216002o | 4791484 | 4791708 | 1692.4 | |
| 19 | Hmel214004o | 8389306 | 8389455 | 1390.7 | |
| 20 | Hmel221001o | 5515125 | 5515215 | 1378.2 | |

***H. m. malleti*, ω**

| | | | | | |
|---|---|---|---|---|---|
| 1 | Hmel210001o | 11548093 | 11548334 | 3481.9 | |
| 2 | Hmel207001o | 8238419 | 8238700 | 3035.0 | |
| 3 | Hmel219001o | 12155812 | 12156047 | 3032.1 | |
| 4 | Hmel213001o | 302788 | 303293 | 3031.5 | |
| 5 | Hmel206001o | 5104806 | 5104982 | 2783.8 | |
| 6 | Hmel215003o | 7885379 | 7885404 | 2783.4 | |
| 7 | Hmel208001o | 4463888 | 4464081 | 2556.7 | |
| 8 | Hmel211001o | 4509794 | 4510695 | 2483.8 | |
| 9 | Hmel201001o | 12939641 | 12939872 | 2470.7 | |
| 10 | Hmel220003o | 9616953 | 9617222 | 2375.1 | |
| 11 | Hmel218003o | 811264 | 811463 | 2262.0 | *optix* |
| 12 | Hmel212001o | 6192598 | 6193015 | 2262.0 | |
| 13 | Hmel205001o | 6491516 | 6491749 | 2261.4 | |
| 14 | Hmel214004o | 439046 | 439088 | 2261.1 | |
| 15 | Hmel220001o | 43827 | 44282 | 2076.9 | |
| 16 | Hmel217001o | 11779651 | 11779804 | 1914.4 | |
| 17 | Hmel204001o | 2900029 | 2900271 | 1845.6 | |
| 18 | Hmel216002o | 3928738 | 3928823 | 1779.0 | |
| 19 | Hmel203003o | 7591363 | 7591454 | 1730.0 | |
| 20 | Hmel209001o | 2459661 | 2459696 | 1567.1 | |

| ***H. m. plesseni,* SweeD** | | | | | |
|---|---|---|---|---|---|
| **Rank** | **Chrom** | **Window** | **CLR** | **Alpha** | **Remarks** |
| 1 | Hmel204001o | 2890000 | 433.36 | 5.75E-05 | |
| 2 | Hmel212001o | 12880000 | 418.13 | 2.97E-05 | |
| 3 | Hmel215003o | 1470000 | 411.35 | 4.08E-05 | *Cortex* |
| 4 | Hmel201001o | 17130000 | 381.6 | 5.14E-05 | |
| 5 | Hmel202001o | 3850000 | 374.52 | 9.77E-05 | |
| 6 | Hmel213001o | 20000 | 345.23 | 7.07E-05 | |
| 7 | Hmel210001o | 11750000 | 267.09 | 6.04E-05 | |
| 8 | Hmel207001o | 7670000 | 210 | 1.14E-04 | |
| 9 | Hmel211001o | 3830000 | 207.28 | 6.67E-05 | |
| 10 | Hmel218003o | 15250000 | 206.87 | 1.51E-04 | |
| 11 | Hmel208001o | 6190000 | 205.36 | 7.02E-05 | |
| 12 | Hmel219001o | 8760000 | 134.17 | 1.71E-04 | |
| 13 | Hmel217001o | 12170000 | 126.76 | 1.29E-04 | |
| 14 | Hmel216002o | 3300000 | 126.15 | 2.53E-04 | |
| 15 | Hmel220003o | 9800000 | 124.2 | 1.40E-04 | |
| 16 | Hmel212001o | 2870000 | 119.11 | 3.83E-04 | |
| 17 | Hmel206001o | 3700000 | 114.59 | 4.76E-04 | |
| 18 | Hmel209001o | 4030000 | 87.13 | 2.53E-04 | |
| 19 | Hmel214004o | 5960000 | 85.81 | 4.16E-04 | |
| 20 | Hmel203003o | 9460001 | 68.14 | 3.59E-04 | |
| ***H. m. malleti,* SweeD** | | | | | |
| 1 | Hmel210001o | 11750000 | 348.43 | 5.60E-05 | |
| 2 | Hmel204001o | 2890000 | 332.46 | 6.73E-05 | |
| 3 | Hmel217001o | 5790000 | 281.7 | 7.48E-05 | |
| 4 | Hmel202001o | 3850000 | 271.03 | 1.11E-04 | |
| 5 | Hmel218003o | 15830000 | 262.91 | 8.09E-05 | |
| 6 | Hmel208001o | 6180000 | 241.13 | 9.66E-05 | |
| 7 | Hmel213001o | 7140000 | 234.26 | 4.54E-05 | |
| 8 | Hmel201001o | 8780000 | 219.35 | 6.14E-05 | |
| 9 | Hmel212001o | 13270000 | 215.57 | 1.61E-04 | |
| 10 | Hmel220003o | 2650000 | 204.56 | 6.10E-05 | |
| 11 | Hmel207001o | 8210000 | 195.55 | 1.47E-04 | |
| 12 | Hmel206001o | 8010000 | 193.65 | 4.22E-05 | |
| 13 | Hmel215003o | 2070000 | 173.58 | 1.50E-04 | |
| 14 | Hmel203003o | 1660000 | 162.25 | 2.91E-04 | |
| 15 | Hmel214004o | 1850000 | 161.15 | 1.40E-04 | |
| 16 | Hmel216002o | 830000 | 158.72 | 9.71E-05 | |
| 17 | Hmel211001o | 5830000 | 148.92 | 1.52E-04 | |
| 18 | Hmel219001o | 3950000 | 111.69 | 1.75E-04 | |
| 19 | Hmel209001o | 4070000 | 103.25 | 2.35E-04 | |
| 20 | Hmel216002o | 3300001 | 100.97 | 2.86E-04 | |

| ***H. m. plesseni,* |iHS|** | | |
|---|---|---|---|---|
| **Rank** | **Chrom** | **Window** | **norm. iHS** | **Remarks** |
| 1 | Hmel218003o | 1230000 | 1 | *optix* |
| 2 | Hmel218003o | 15130000 | 0.942761 | |
| 3 | Hmel215003o | 1250000 | 0.889474 | *Cortex* |
| 4 | Hmel213001o | 10320000 | 0.835897 | *vvl* |
| 5 | Hmel218003o | 5380000 | 0.798742 | |
| 6 | Hmel201001o | 11860000 | 0.786517 | |
| 7 | Hmel210001o | 260000 | 0.762712 | |
| 8 | Hmel220003o | 11850000 | 0.752604 | |
| 9 | Hmel206001o | 8020000 | 0.725664 | |
| 10 | Hmel201001o | 10450000 | 0.713235 | |
| 11 | Hmel218003o | 2650000 | 0.6875 | |
| 12 | Hmel211001o | 5520000 | 0.629108 | |
| 13 | Hmel210001o | 3370000 | 0.627273 | *WntA* |
| 14 | Hmel212001o | 12790000 | 0.597315 | |
| 15 | Hmel206001o | 11040000 | 0.593023 | |
| 16 | Hmel210001o | 16130000 | 0.584 | |
| 17 | Hmel220003o | 5430000 | 0.56962 | |
| 18 | Hmel203003o | 6080000 | 0.564103 | |
| 19 | Hmel210001o | 10400000 | 0.53125 | |
| 20 | Hmel219001o | 11560000 | 0.503311 | |
| ***H. m. malleti,* |iHS|** | | |
| 1 | Hmel218003o | 15110000 | 0.997135 | |
| 2 | Hmel202001o | 1820000 | 0.918728 | |
| 3 | Hmel220003o | 4860000 | 0.813253 | |
| 4 | Hmel206001o | 8030000 | 0.79798 | |
| 5 | Hmel213001o | 15760000 | 0.785408 | |
| 6 | Hmel201001o | 12540000 | 0.77193 | |
| 7 | Hmel207001o | 9050000 | 0.712329 | |
| 8 | Hmel212001o | 16140000 | 0.688259 | |
| 9 | Hmel212001o | 10600000 | 0.657778 | |
| 10 | Hmel201001o | 10450000 | 0.61324 | |
| 11 | Hmel201001o | 8590000 | 0.595745 | |
| 12 | Hmel208001o | 6190000 | 0.578947 | |
| 13 | Hmel206001o | 13110000 | 0.564453 | |
| 14 | Hmel217001o | 5790000 | 0.551913 | |
| 15 | Hmel205001o | 4460000 | 0.550926 | |
| 16 | Hmel220003o | 12100000 | 0.545455 | |
| 17 | Hmel206001o | 11050000 | 0.526042 | |
| 18 | Hmel212001o | 6280000 | 0.507653 | |
| 19 | Hmel218002o | 140000 | 0.507463 | |

**Table S10. Top loci based on ω, sweeD and *iHS* in *H. melpomene.***

The top 20 loci found across the genome for the various selection statistics (only 19 loci rise above the threshold value of 0.5 in the case of iHS in *H. m. malleti*). Windows that are within 10 kbp of scaffold boundaries are excluded to avoid edge artifact.

| Species | Locus | Centre (SL) | | Width (SL) in km | |
|---|---|---|---|---|---|
| | | Phenotype | Genotype | Phenotype | Genotype |
| *H. erato* | *optix* | 30.80 (28.3–33.1) | 31.87 (29.3–34.3) | 15.59 (9.7–23.3) | 15.70 (8.7–25.0) |
| | *WntA* | 46.29 (41.8–51.5) | 47.15 (43.0–51.7) | 20.93 (6.8–37.8) | 18.95 (6.5–34.5) |
| | *Ro* | 52.43 (44.9–59.2) | 52.71 (46.2–58.8) | 39.43 (23.0–55.9) | 39.72 (24.7–55.5) |
| | LG2 *Inv* | n.a. | 46.63 (41.24 – 53.43) | n.a. | 53.44 (24.67 – 102.63) |
| *H. melpomene* | *optix* | 28.55 (25.1–31.2) | 28.91 (25.6–31.4) | 15.85 (10.6–22.4) | 15.12 (9.7–22.1) |
| | *WntA* | 49.38 (46.0–52.8) | 49.78 (45.7–54.4) | 23.59 (15.4–32.5) | 24.82 (15.0–35.9) |
| | *N* | 31.37 (28.2–34.2) | 31.15 (27.0–35.1) | 15.76 (8.4–25.7) | 22.79 (12.0–35.8) |

**Table S11. Cline analysis at five different loci.**

Maximum likelihood estimates of cline centre and width, with two log-likelihood support limits (SL), for the allele frequency and phenotypic clines of the major color pattern loci and the polymorphic inversion.

| Chr | Scaffold | Position | Best model | Cen (km) | Width (km) | $F_{ST}$ (site) | $F_{ST}$ (win) | HMM state | Description |
|---|---|---|---|---|---|---|---|---|---|
| 1 | Herato0101 | 7538046 | IV | 29.3 | 0.5 | 0.55 | 0.08 | 1 | no peak |
| 1 | Herato0101 | 12588853 | IV | 10.5 | 5.4 | 0.51 | 0.07 | 1 | no peak |
| 10 | Herato1001 | 4673183 | I | 51.4 | 29.3 | 1 | 0.65 | 2 | *WntA* |
| 12 | Herato1202 | 10753259 | III | 27.2 | 12.2 | 0.51 | 0.06 | 1 | no peak |
| 15 | Herato1505 | 2096799 | I | 32.7 | 29.7 | 0.78 | 0.45 | 2 | *cortex* |
| 15 | Herato1507 | 3016730 | IV | 10.2 | 0.9 | 0.5 | 0.05 | 1 | no peak |
| 17 | Herato1701 | 1663642 | IV | 27.7 | 19 | 0.52 | 0.12 | 1 | no peak |
| 18 | Herato1801 | 387738 | IV | 10.3 | 2.7 | 0.54 | 0.08 | 1 | no peak |
| 18 | Herato1801 | 632087 | III | 24.5 | 30 | 0.81 | 0.23 | 2 | *optix* |
| 18 | Herato1801 | 892383 | IV | 24.8 | 24.5 | 0.77 | 0.18 | 2 | *optix* |
| 18 | Herato1801 | 958379 | I | 26.5 | 28 | 0.92 | 0.36 | 2 | *optix* |
| 18 | Herato1801 | 1055518 | IV | 18.2 | 29.6 | 0.82 | 0.25 | 2 | *optix* |
| 18 | Herato1801 | 1182167 | IV | 28 | 18.1 | 0.99 | 0.56 | 2 | *optix* |
| 18 | Herato1801 | 1261271 | III | 29.1 | 17.1 | 1 | 0.75 | 2 | *optix* |
| 18 | Herato1801 | 1389883 | I | 29.5 | 15.4 | 1 | 0.94 | 2 | *optix* |
| 18 | Herato1801 | 1419325 | I | 28.5 | 16 | 1 | 0.9 | 2 | *optix* |
| 18 | Herato1801 | 1586723 | IV | 23.2 | 26.8 | 0.85 | 0.32 | 2 | *optix* |
| 18 | Herato1805 | 4375044 | III | 9.6 | 7 | 0.54 | 0.08 | 1 | no peak |
| 19 | Herato1901 | 2003842 | III | 26.6 | 11.1 | 0.59 | 0.08 | 1 | no peak |
| 19 | Herato1904 | 2840926 | III | 30.4 | 21.9 | 0.84 | 0.32 | 2 | small peak |
| 20 | Herato2001 | 3935391 | IV | 12.4 | 5.3 | 0.51 | 0.04 | 1 | no peak |
| 20 | Herato2001 | 10912020 | III | 29.5 | 4.5 | 0.55 | 0.06 | 1 | no peak |

| | | | | | | | | | |
|---|---|---|---|---|---|---|---|---|---|
| 20 | Herato2001 | 15219605 | III | 26.8 | 5.8 | 0.71 | 0.3 | 2 | small peak |
| 21 | Herato2101 | 1007837 | IV | 6.9 | 29.7 | 0.5 | 0.04 | 1 | no peak |
| 21 | Herato2101 | 8501906 | IV | 10.3 | 7.1 | 0.52 | 0.06 | 1 | no peak |

**Table S12. Loci with narrow clines in *H. erato.***

| | |
|---|---|
| Chr | Chromosome |
| Best model | I: fixed maximum and minimum allele frequencies, no exponential tails |
| | III: fixed maximum and minimum allele frequencies; two exponential tails mirrored about the cline centre |
| | IV: estimated maximum and minimum allele frequencies, no exponential tails |
| Cen | Centre |
| HMM state: | 1: background; 2: high differentiation |

| Chr | Scaffold | Position | Best model | Cen (km) | Width (km) | $F_{ST}$ (site) | $F_{ST}$ (win) | HMM state | Description |
|---|---|---|---|---|---|---|---|---|---|
| 1 | Hmel201001o | 11097620 | IV | 14.9 | 9.2 | 0.51 | 0.07 | 1 | no peak |
| 2 | Hmel202001o | 2274430 | IV | 8.7 | 7.7 | 0.44 | 0.1 | 1 | tiny peak |
| 4 | Hmel204001o | 4135399 | IV | 11.1 | 27.6 | 0.51 | 0.06 | 1 | no peak |
| 7 | Hmel207001o | 1573526 | IV | 10.5 | 14.1 | 0.4 | 0.05 | 1 | no peak |
| 9 | Hmel209001o | 8354789 | IV | 13.7 | 2.8 | 0.45 | 0.02 | 1 | no peak |
| 10 | Hmel210001o | 1872141 | IV | 16.9 | 13.1 | 0.44 | 0.04 | 1 | no peak |
| 10 | Hmel210001o | 1922772 | IV | 56.5 | 7.3 | 0.41 | 0.03 | 1 | no peak |
| 10 | Hmel210001o | 3321549 | I | 46.3 | 24.7 | 1 | 0.88 | 2 | *WntA* |
| 11 | Hmel211001o | 8781462 | IV | 17.1 | 7.6 | 0.44 | 0.02 | 1 | no peak |
| 11 | Hmel211001o | 9047961 | IV | 11.5 | 4.4 | 0.4 | 0.03 | 1 | no peak |
| 11 | Hmel211001o | 9912550 | IV | 11.8 | 13.2 | 0.45 | 0.04 | 1 | no peak |
| 11 | Hmel211001o | 10070575 | IV | 12.4 | 2.9 | 0.43 | 0.03 | 1 | no peak |
| 13 | Hmel213001o | 10126057 | IV | 34.4 | 14.9 | 0.65 | 0.17 | 2 | *vvl* |
| 13 | Hmel213001o | 10562270 | IV | 16.4 | 7.2 | 0.41 | 0.04 | 1 | *vvl* |
| 14 | Hmel214004o | 1337697 | IV | 32.8 | 17.1 | 0.58 | 0.12 | 1 | no peak |
| 14 | Hmel214004o | 1537776 | IV | 8.2 | 15.1 | 0.4 | 0.03 | 1 | no peak |
| 16 | Hmel216002o | 1173257 | IV | 20.7 | 23.1 | 0.54 | 0.03 | 1 | no peak |
| 16 | Hmel216002o | 5376532 | IV | 8.9 | 12.8 | 0.46 | 0.03 | 1 | no peak |
| 17 | Hmel217001o | 10244627 | IV | 13.6 | 10.4 | 0.42 | 0.04 | 1 | no peak |
| 18 | Hmel218003o | 781492 | IV | 31.7 | 14.2 | 1 | 0.08 | 1 | *optix* |
| 18 | Hmel218003o | 812325 | I | 31.5 | 17.7 | 1 | 0.52 | 2 | *optix* |
| 18 | Hmel218003o | 923741 | IV | 31.3 | 14.5 | 0.94 | 0.8 | 2 | *optix* |
| 18 | Hmel218003o | 1028407 | III | 29.6 | 12.7 | 0.77 | 0.93 | 2 | *optix* |
| 18 | Hmel218003o | 2661551 | IV | 30.9 | 16.9 | 1 | 0.3 | 2 | second big peak |

| | | | | | | | | | |
|---|---|---|---|---|---|---|---|---|---|
| 18 | Hmel218003o | 9311544 | IV | 16.0 | 0.5 | 0.41 | 0.03 | 1 | no peak |
| 18 | Hmel218003o | 13038669 | IV | 13.4 | 3.7 | 0.47 | 0.03 | 1 | no peak |
| 19 | Hmel219001o | 5975768 | IV | 8.4 | 7.4 | 0.41 | 0.08 | 1 | no peak |
| 21 | Hmel221001o | 1140770 | IV | 18.8 | 0.5 | 0.44 | 0.04 | 1 | no peak |
| 21 | Hmel221001o | 1271592 | IV | 29.3 | 0.1 | 0.46 | 0.03 | 1 | no peak |
| 21 | Hmel221001o | 4674519 | IV | 12.3 | 13.2 | 0.41 | 0.06 | 1 | no peak |
| 21 | Hmel221001o | 7581777 | IV | 10.4 | 6.8 | 0.42 | 0.3 | 2 | tiny peak |

**Table S13. Loci with narrow clines in *H. melpomene*.**

Chr Chromosome
Best model I: fixed maximum and minimum allele frequencies, no exponential tails
III: fixed maximum and minimum allele frequencies; two exponential tails mirrored about the cline centre
IV: estimated maximum and minimum allele frequencies, no exponential tails
Cen Centre
HMM state: 1: background; 2: high differentiation

| *H. erato* | | | | *H. melpomene* | | | |
|---|---|---|---|---|---|---|---|
| **Locus** | **n** | $\widehat{F}_{IS}$ | **ΔLog(*L*)** | **Locus** | **n** | $\widehat{F}_{IS}$ | **ΔLog(*L*)** |
| *WntA* | 11 | 0.058 | 0.15 | *WntA* | 16 | 0.045 | 0.1 |
| *Ro* | 14 | 0.006 | 0 | Homologue of *Ro* | 14 | 0.006 | 0 |
| cortex | 17 | 0.281 | 3.54 | *cortex* | 14 | 0.314 | 4.65 |
| *optix* | 9 | 0 | 0 | *optix* | 14 | 0 | 0 |

**Table S14. Maximum likelihood estimates for heterozygote deficit, $\widehat{F}_{IS}$.**

Fitting the same value for all polymorphic samples. There is no significant deviation, except at *cortex*, in both species. Assuming an asymptotic $\chi_1^2$ distribution for 2ΔLog(L), this corresponds to *P* = 0.78%, 0.23% in erato and in melpomene.

| Species | Type | Cline pairs | $\hat{R}$ | ΔLog($L$) | limits |
|---|---|---|---|---|---|
| *H. erato* | coincident | 2 | -0.025 | 0.21 | -0.099 – 0.054 |
| | displaced | 4 | -0.016 | 0.16 | -0.075 – 0.049 |
| *H. melpomene* | coincident | 3 | 0.042 | 0.28 | -0.068 – 0.154 |
| | displaced | 3 | 0.007 | 0.01 | -0.083 – 0.098 |

**Table S15. Maximum likelihood estimates for the correlation between loci ( $R = D/\sqrt{p_1 q_1 p_2 q_2}$ ) , together with the difference in log(likelihood) relative to $R = 0$.**

The support limits correspond to a drop in log(likelihood) by 2 units, asymptotically equivalent to 95% confidence limits. Including all polymorphic populations. Including only the highly polymorphic samples ( $\sqrt{p_1 q_1 p_2 q_2} > 0.1$) makes little difference.

| | *H. erato* | | | | *H. melpomene* | | | |
|---|---|---|---|---|---|---|---|---|
| | *WntA* | *Ro* | *cortex* | *optix* | *WntA* | *vvl* | *cortex* | *optix* |
| width | 29.9 | 47.8 | 38.0 | 17.0 | 32.4 | 36.8 | 31.3 | 14.9 |
| position | 47.3 | 55.7 | 26.2 | 28.2 | 43.5 | 34.8 | 30.3 | 29.8 |
| dominance | +4.38 | +0.40 | +6.13 | -9.28 | +2.28 | -3.09 | -4.56 | +4.13 |

**Table S16. Testing for asymmetry of single-locus clines.**

Linear frequency dependence, with no dominance, maintains a symmetric cline, whereas linear positive frequency dependence, with full dominance of the lowland alleles, maintains asymmetric clines, with introgression into the lowland population. The table shows the estimated position and width for each locus, in each species, for the best-fitting model. The last row shows the difference in log likelihood between the best-fitting asymmetric vs. symmetric models; a positive value favors asymmetry, and a value greater than 2 conventionally indicates significance. None of the best-fitting models gave evidence for residual variation (i.e., we estimate $F_{ST} = 0$). Likelihoods are calculated using a beta-binomial model. Note that *Ro* in *H. melpomene* refers to the homologous locus *vvl*.
