## Supplementary material for "Haplotype tagging reveals parallel formation of hybrid races in two butterfly species": Data S4

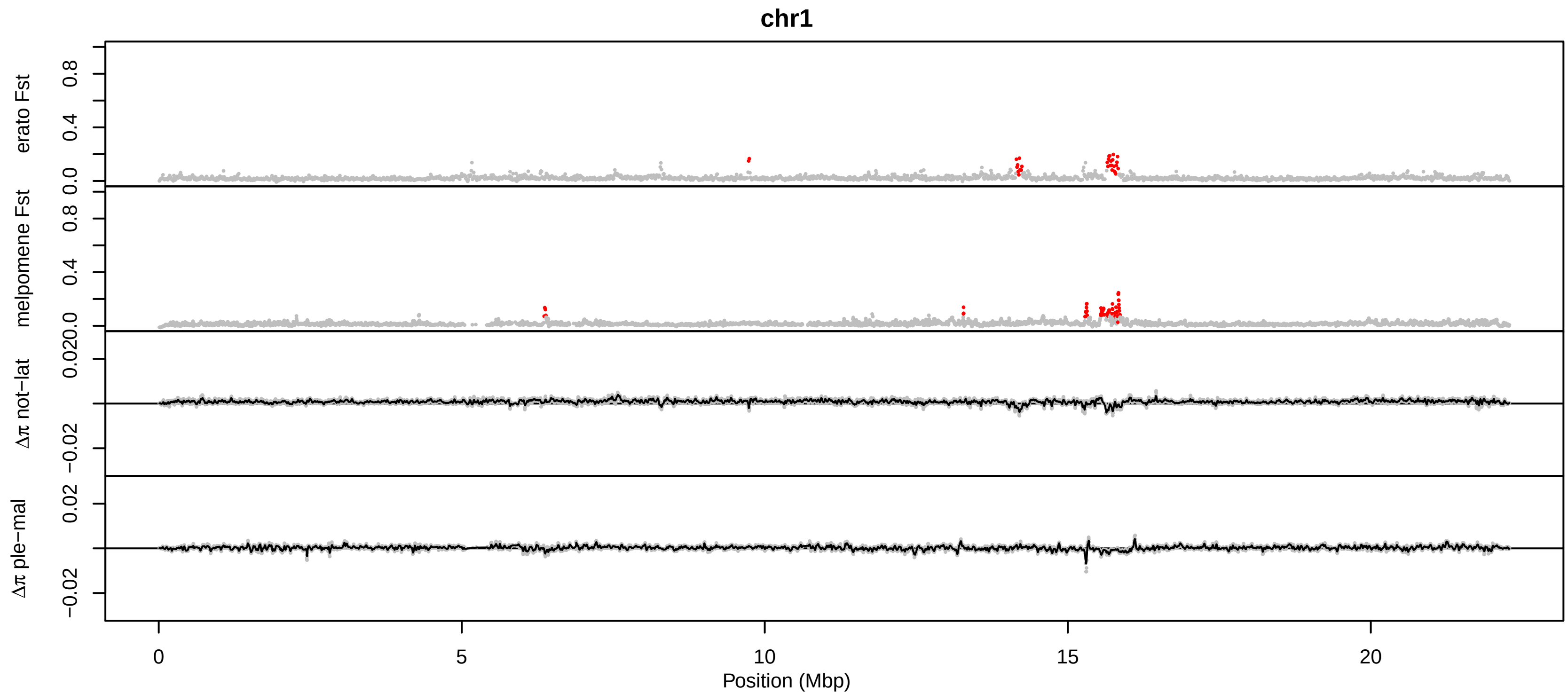
chr1
erato Fst
melpomene Fst
Δπ not−lat
Δπ ple−mal
0.0
0.4
0.8
0.020
−0.02
0.02
0
5
10
15
20
Position (Mbp)

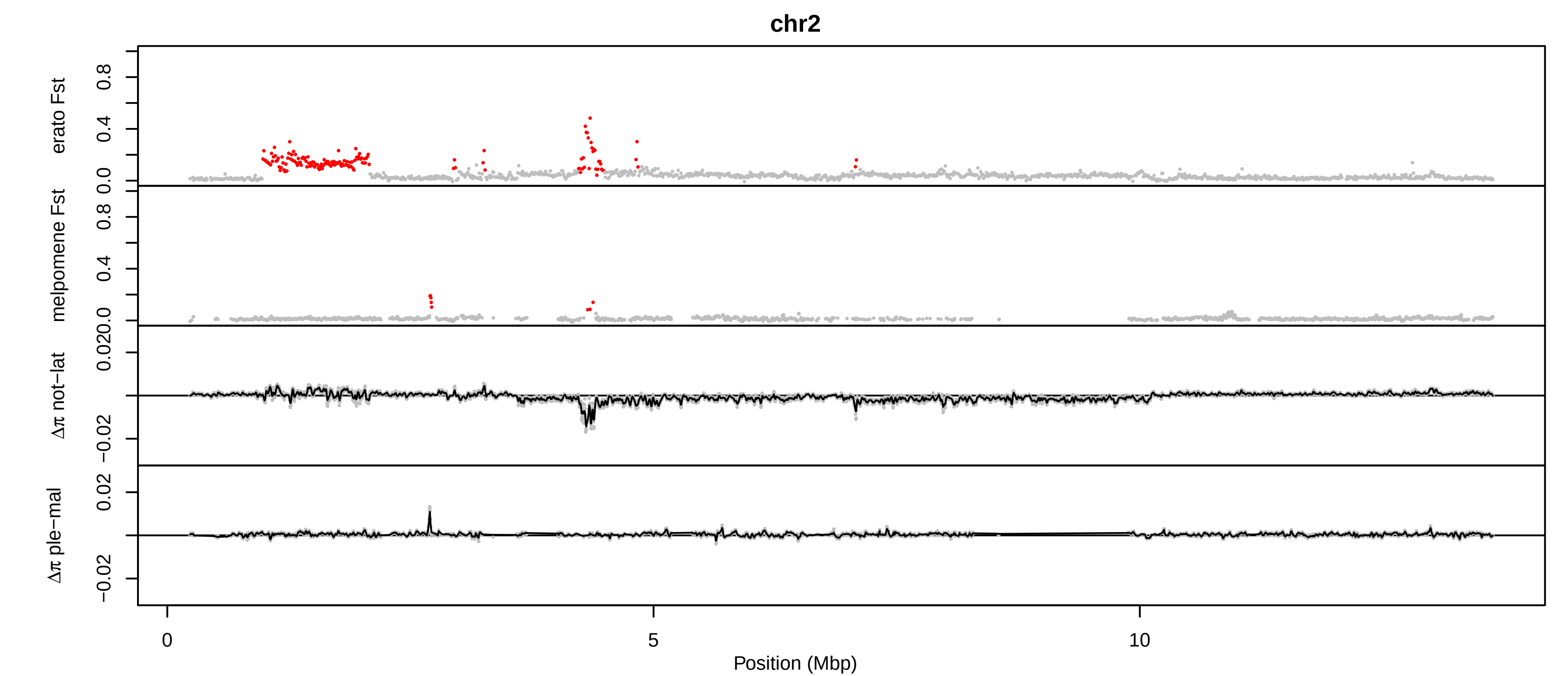

chr2
erato Fst
melpomene Fst
Δπ not−lat
Δπ ple−mal
0.0
0.4
0.8
0.020
−0.02
0.02
0
5
10
Position (Mbp)

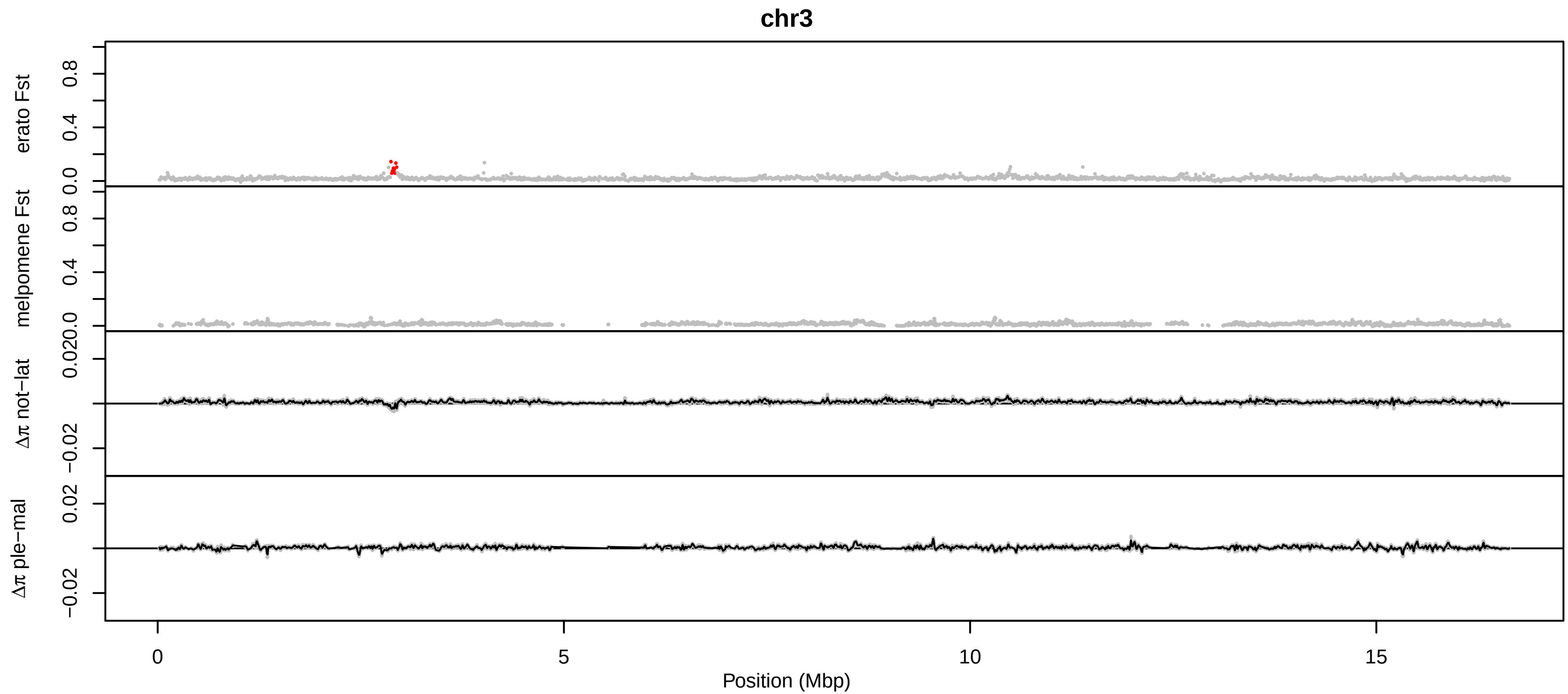

chr3
erato Fst
melpomene Fst
Δπ not−lat
Δπ ple−mal
0.0
0.4
0.8
0.020
0.02
−0.02
0
5
10
15
Position (Mbp)

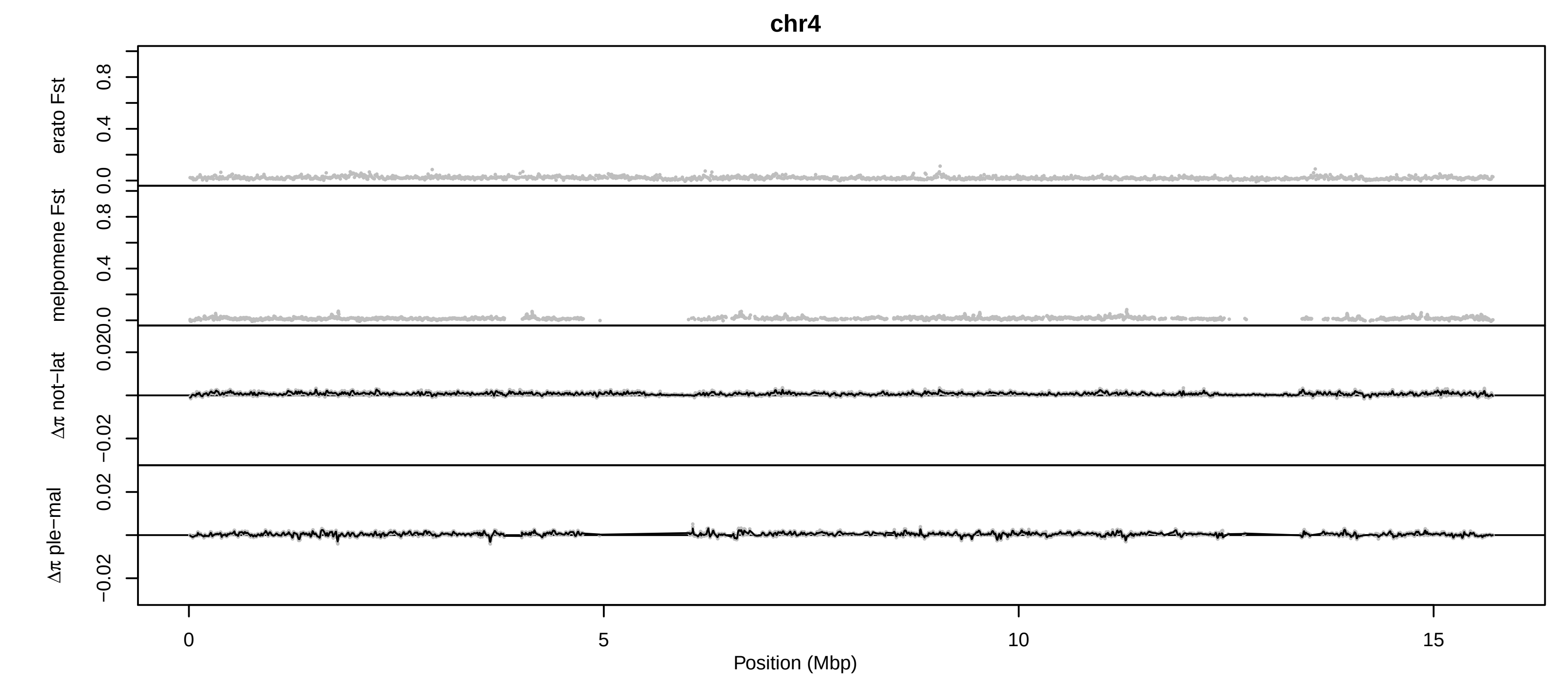

chr4
erato Fst
melpomene Fst
Δπ not−lat
Δπ ple−mal
0.0
0.4
0.8
0.020
−0.02
0.02
0
5
10
15
Position (Mbp)

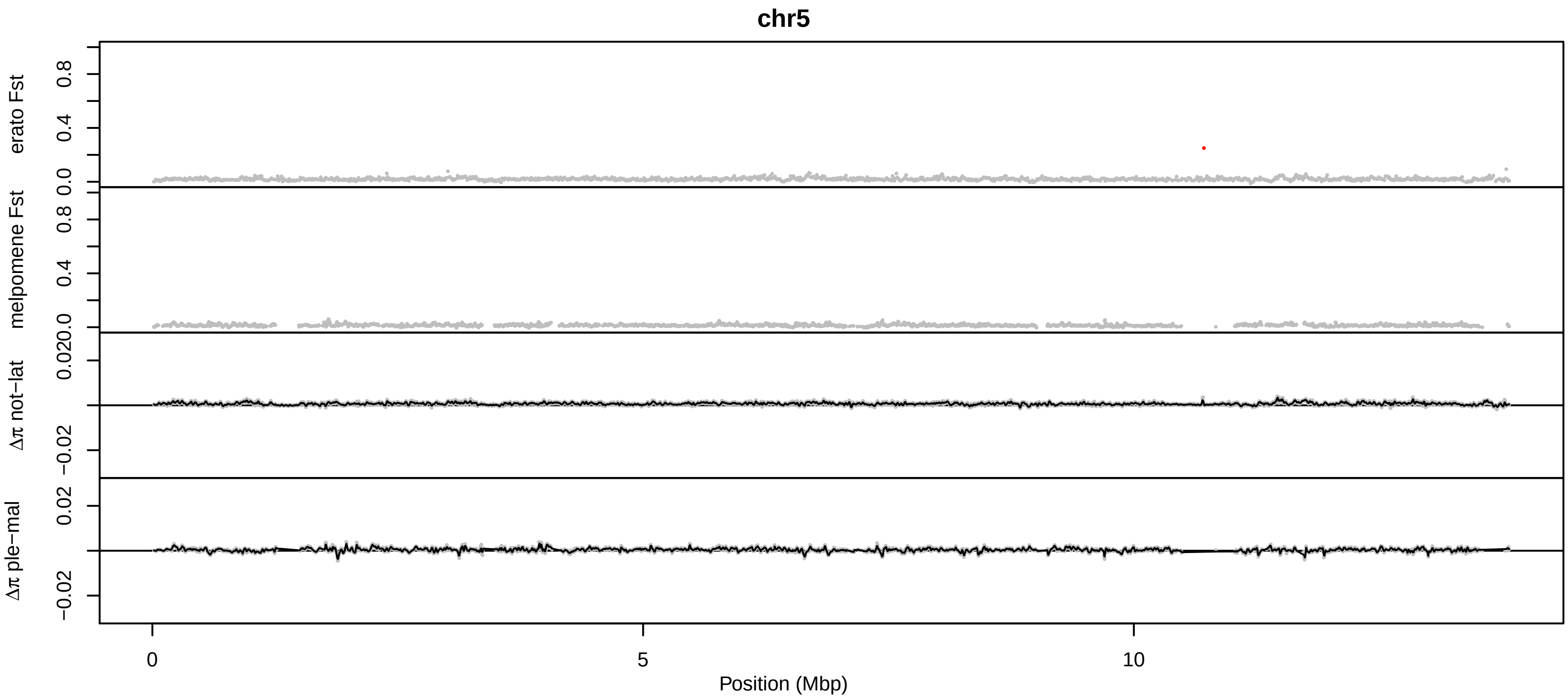
chr5
erato Fst
melpomene Fst
Δπ not−lat
Δπ ple−mal
0.0
0.4
0.8
0.020
−0.02
0.02
0
5
10
Position (Mbp)

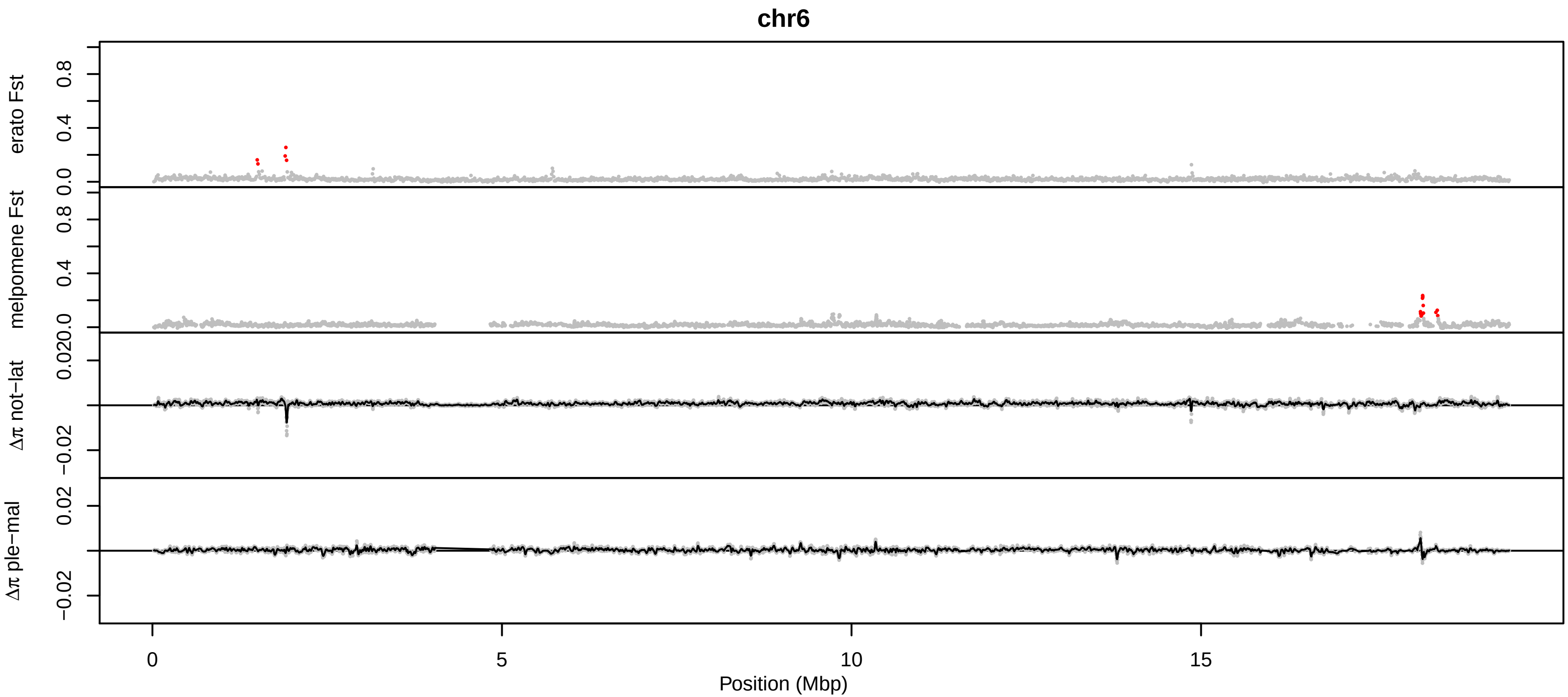
chr6
erato Fst
melpomene Fst
Δπ not−lat
Δπ ple−mal
0.0
0.4
0.8
0.020
−0.02
0.02
0
5
10
15
Position (Mbp)

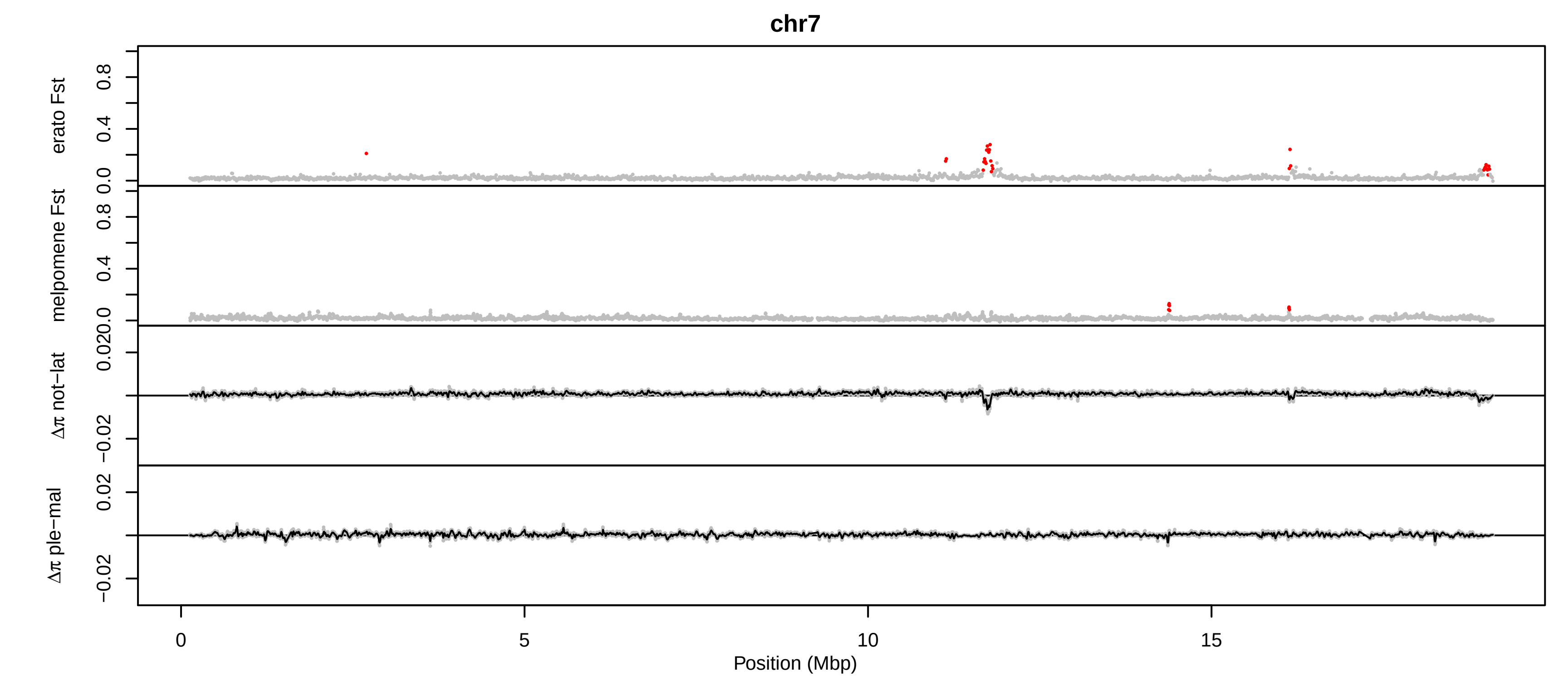
chr7
erato Fst
melpomene Fst
Δπ not−lat
Δπ ple−mal
0.0
0.4
0.8
0.020
−0.02
0.02
0
5
10
15
Position (Mbp)

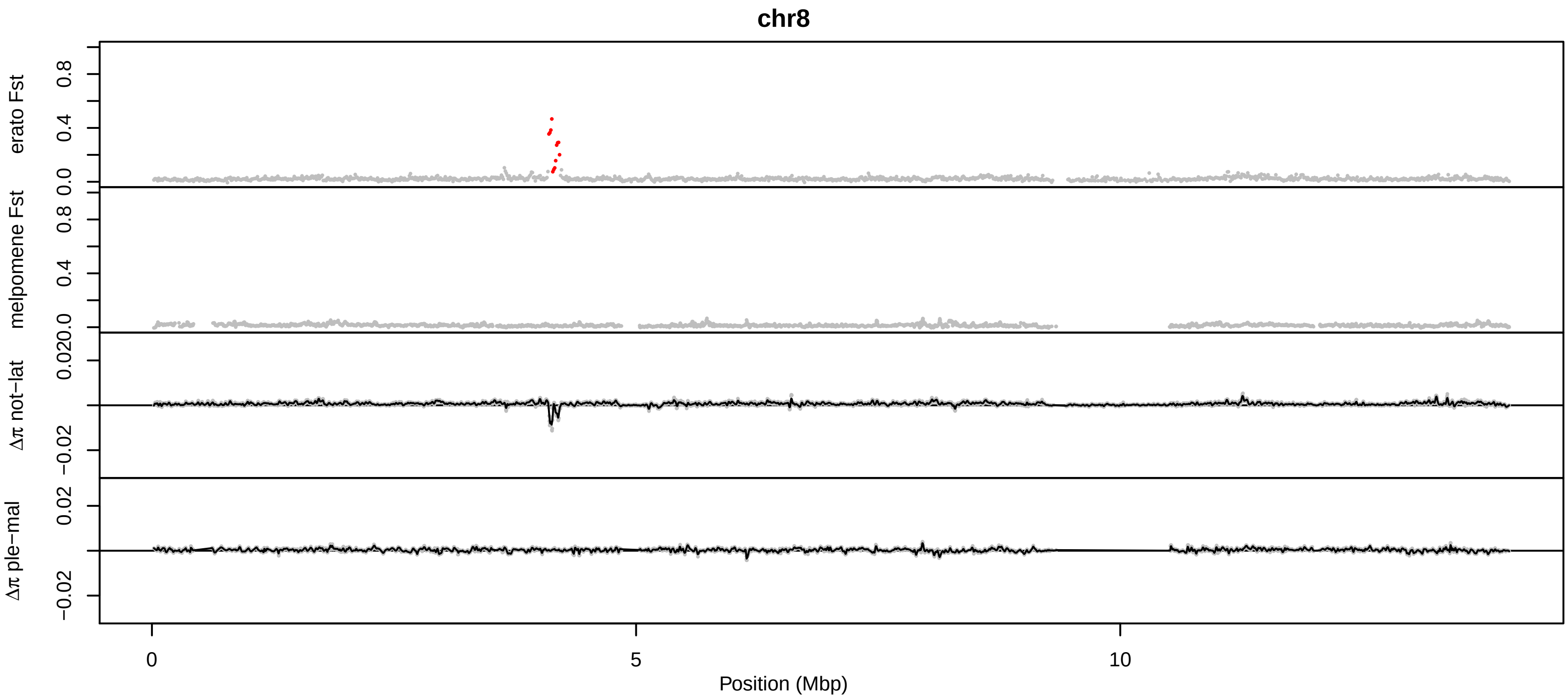
chr8
erato Fst
melpomene Fst
Δπ not−lat
Δπ ple−mal
0.0
0.4
0.8
0.0
0.4
0.8
0.020
−0.02
0.02
−0.02
0
5
10
Position (Mbp)

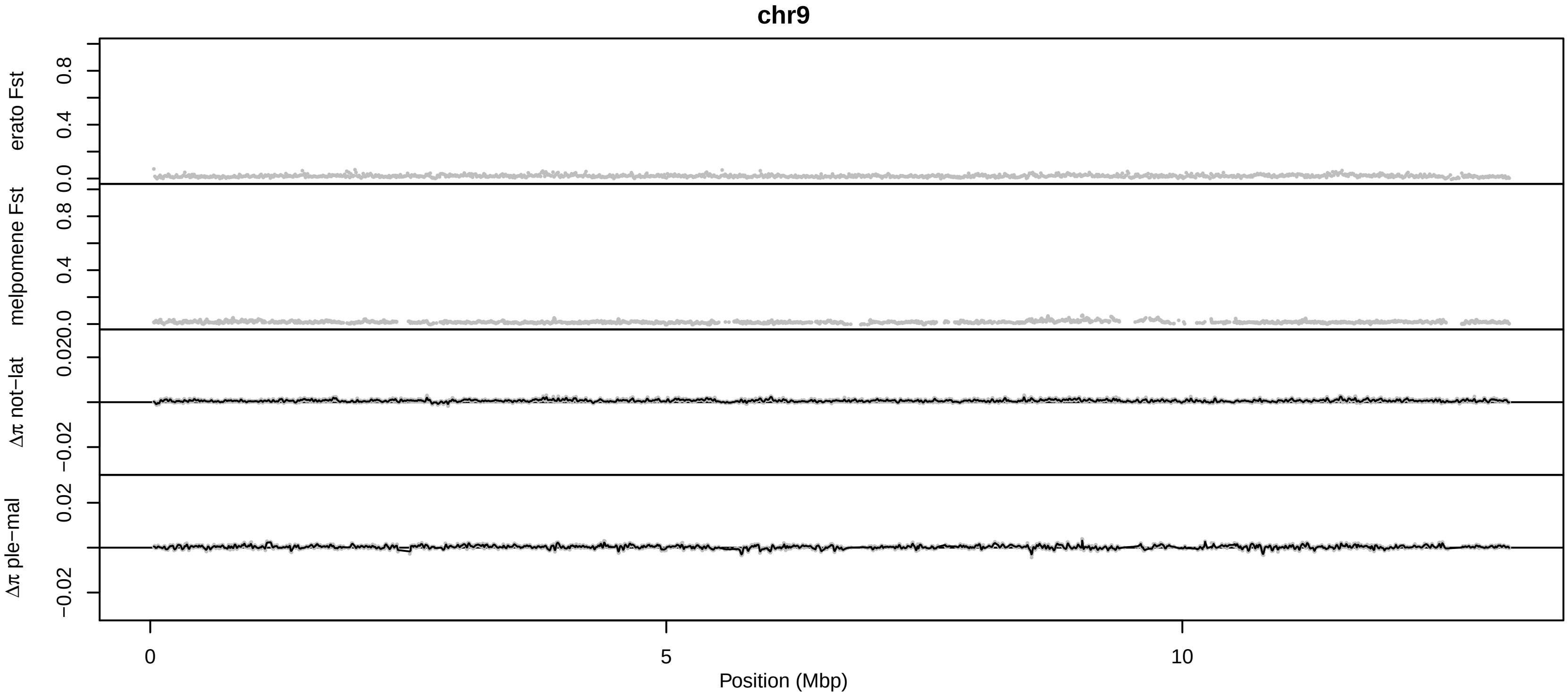

chr9
erato Fst
melpomene Fst
Δπ not−lat
Δπ ple−mal
0.0
0.4
0.8
0.020
−0.02
0.02
0
5
10
Position (Mbp)

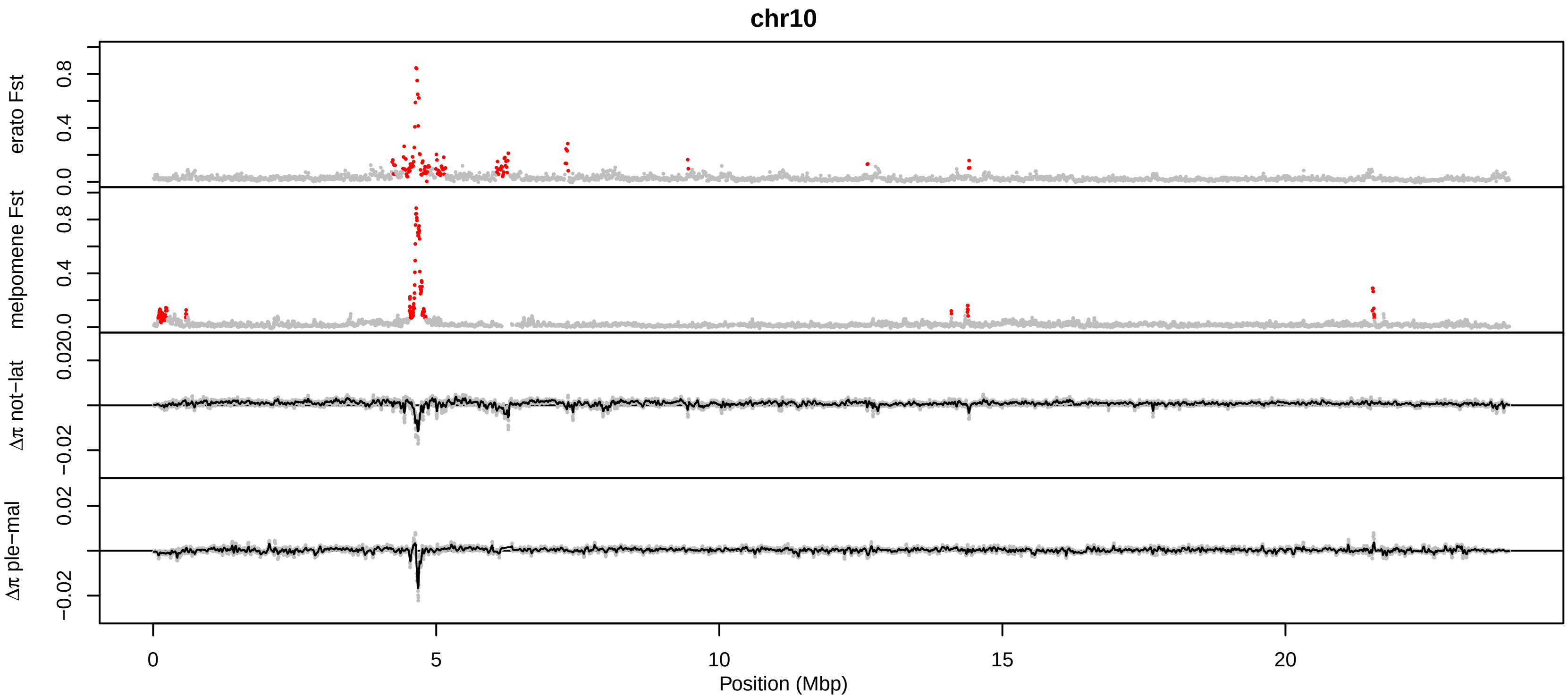
chr10
erato Fst
melpomene Fst
Δπ not−lat
Δπ ple−mal
0.0
0.4
0.8
0.020
−0.02
0.02
0
5
10
15
20
Position (Mbp)

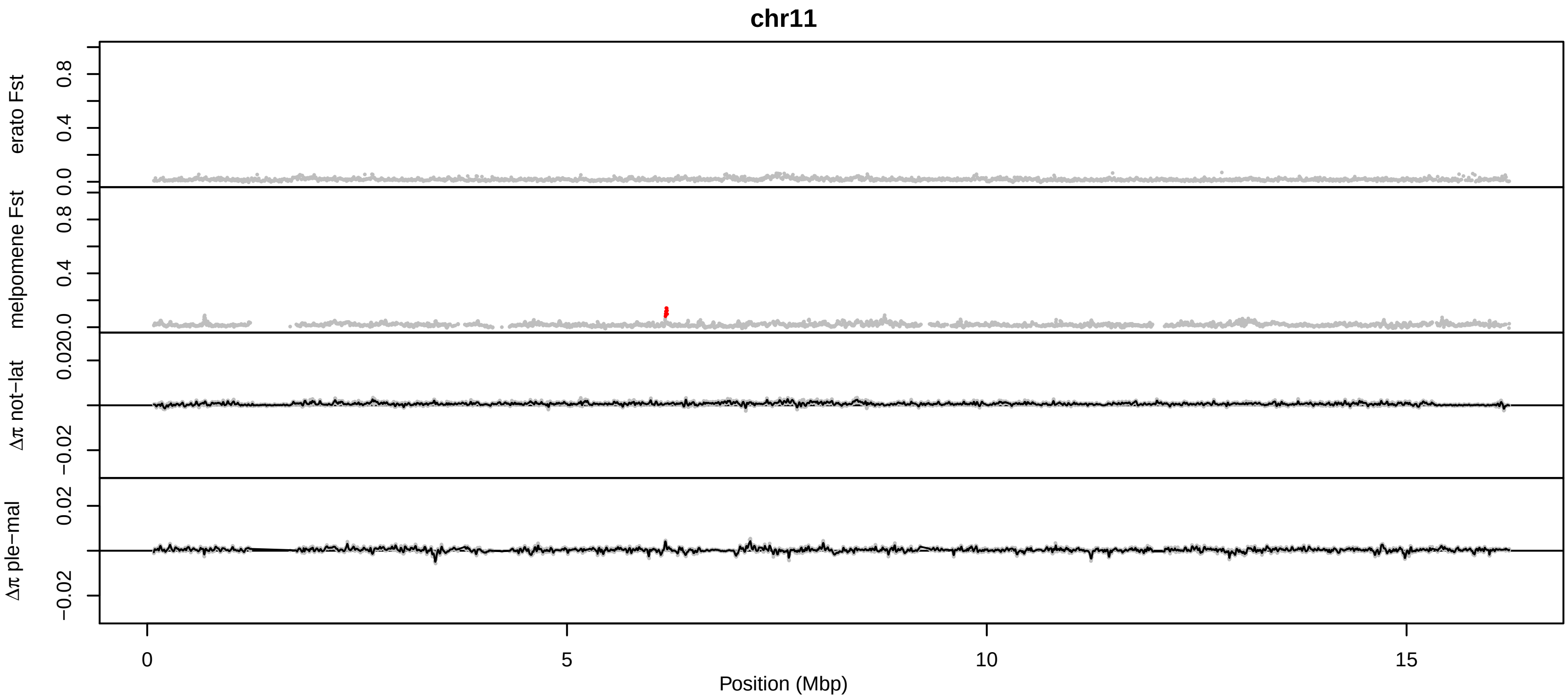
chr11
erato Fst
melpomene Fst
Δπ not−lat
Δπ ple−mal
0.0
0.4
0.8
0.020
−0.02
0.02
0
5
10
15
Position (Mbp)

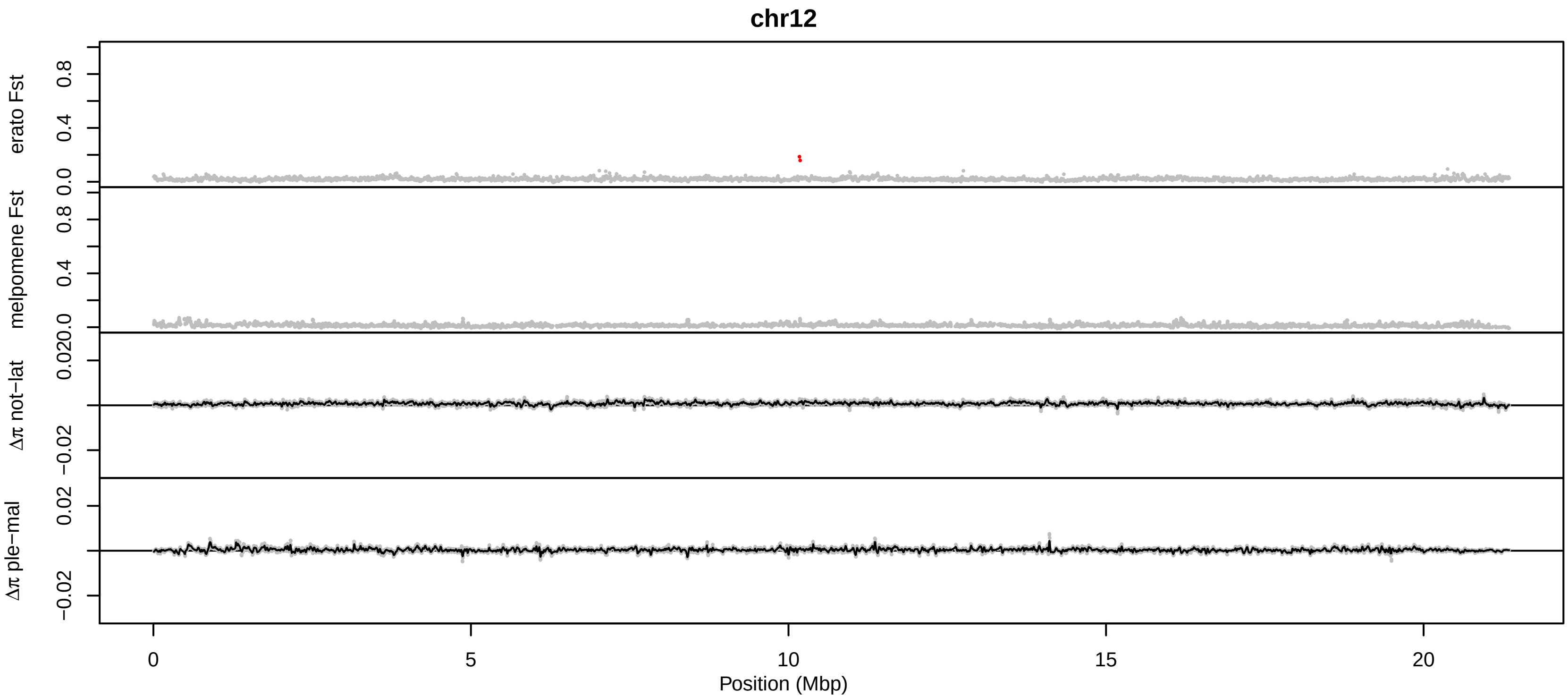

chr12
erato Fst
melpomene Fst
Δπ not−lat
Δπ ple−mal
0.0
0.4
0.8
0.02
0.00
−0.02
0
5
10
15
20
Position (Mbp)

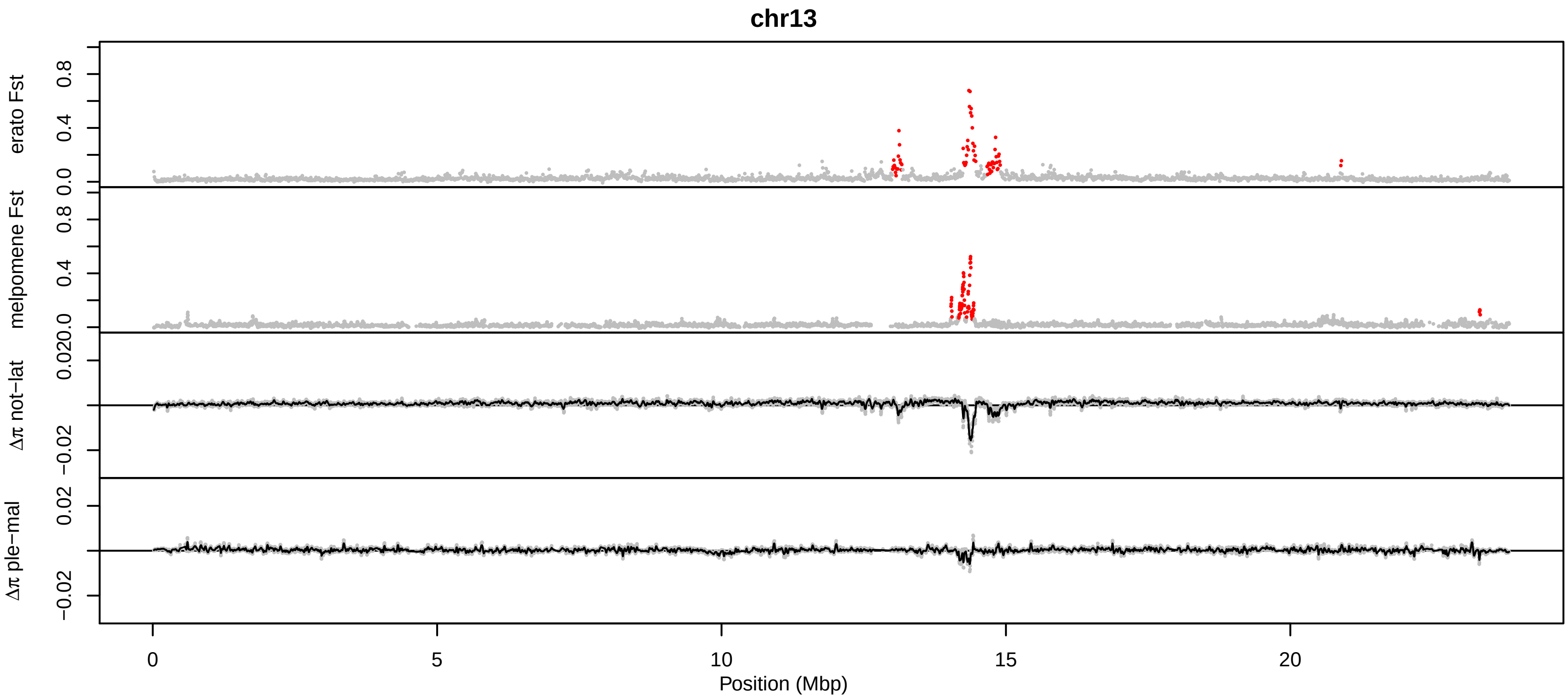
chr13
erato Fst
melpomene Fst
Δπ not−lat
Δπ ple−mal
0.0
0.4
0.8
0.020
−0.02
0.02
0
5
10
15
20
Position (Mbp)

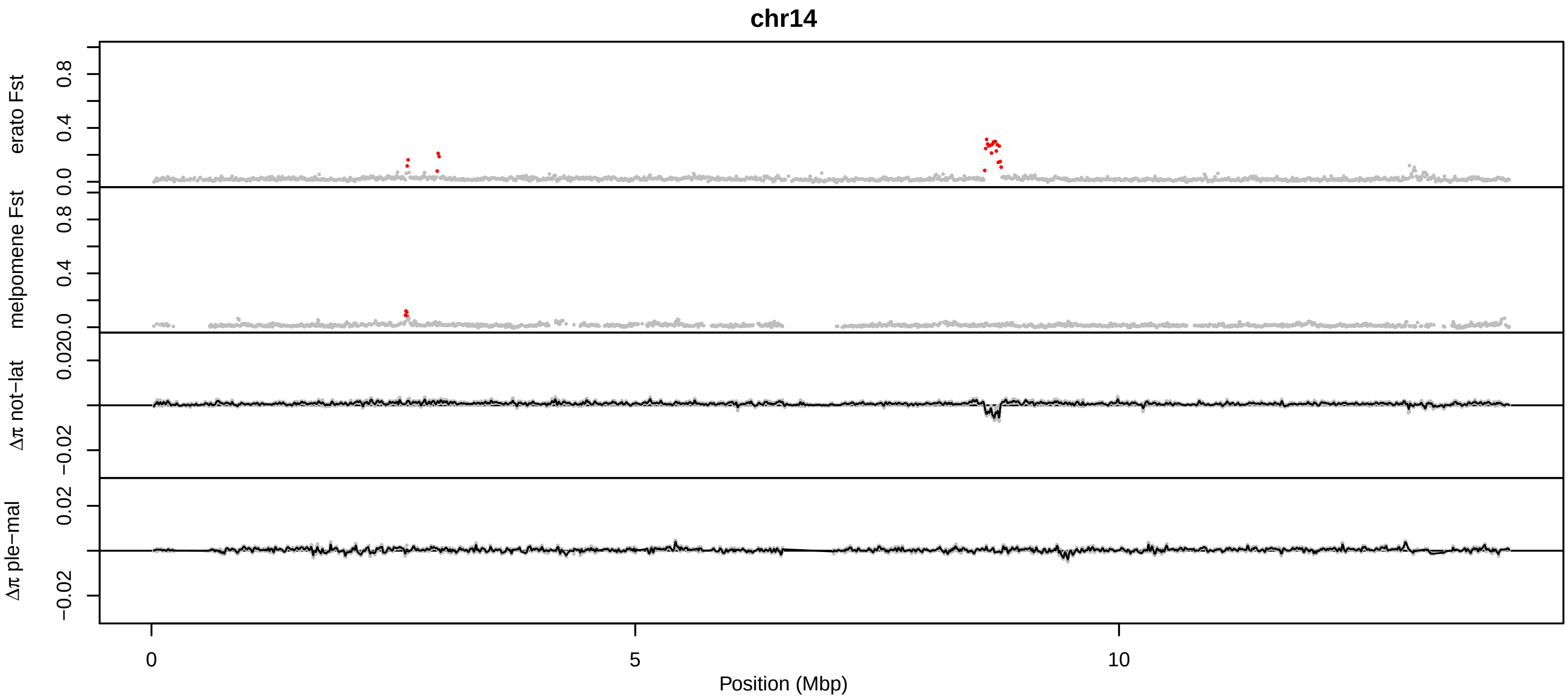

chr14
erato Fst
melpomene Fst
Δπ not−lat
Δπ ple−mal
0.0
0.4
0.8
0.020
−0.02
0.02
0
5
10
Position (Mbp)

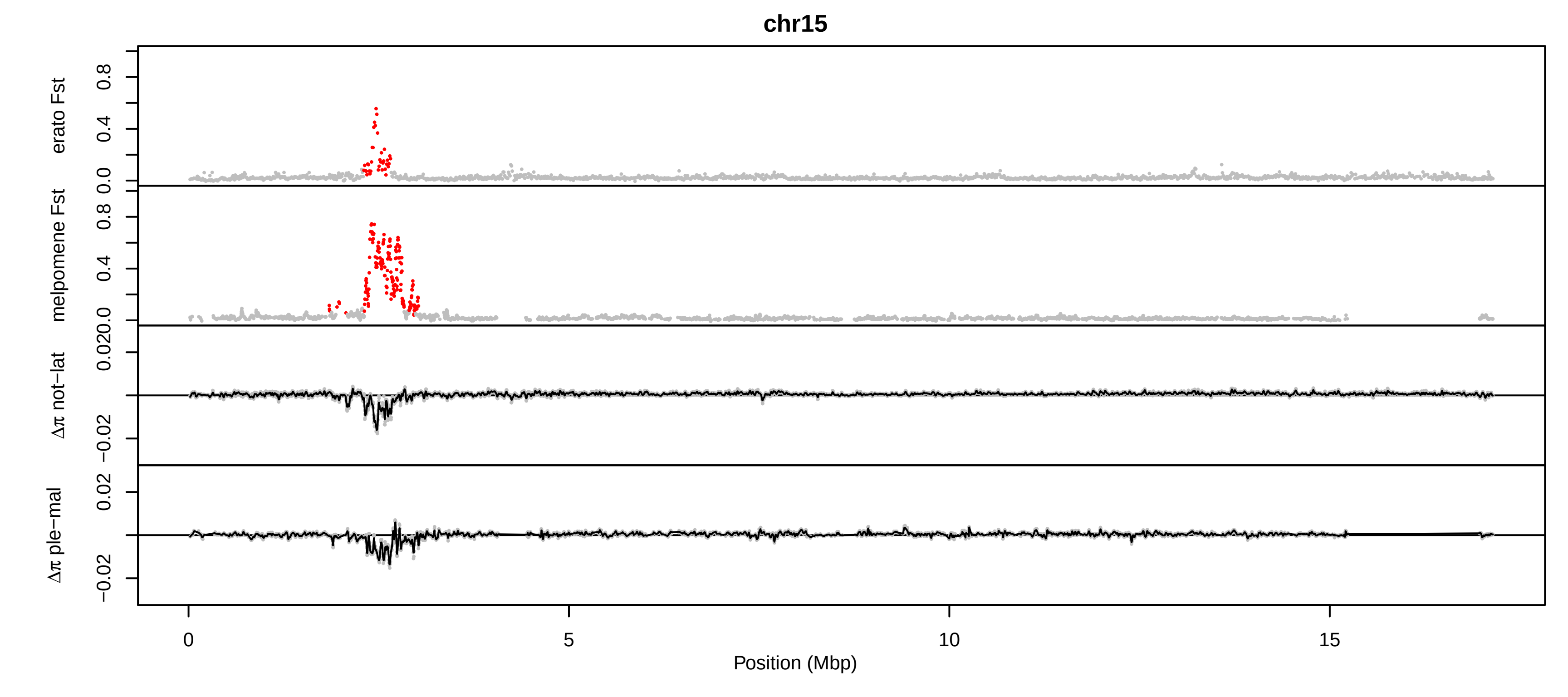
chr15
erato Fst
melpomene Fst
Δπ not−lat
Δπ ple−mal
0.0
0.4
0.8
0.020
−0.02
0.02
0
5
10
15
Position (Mbp)

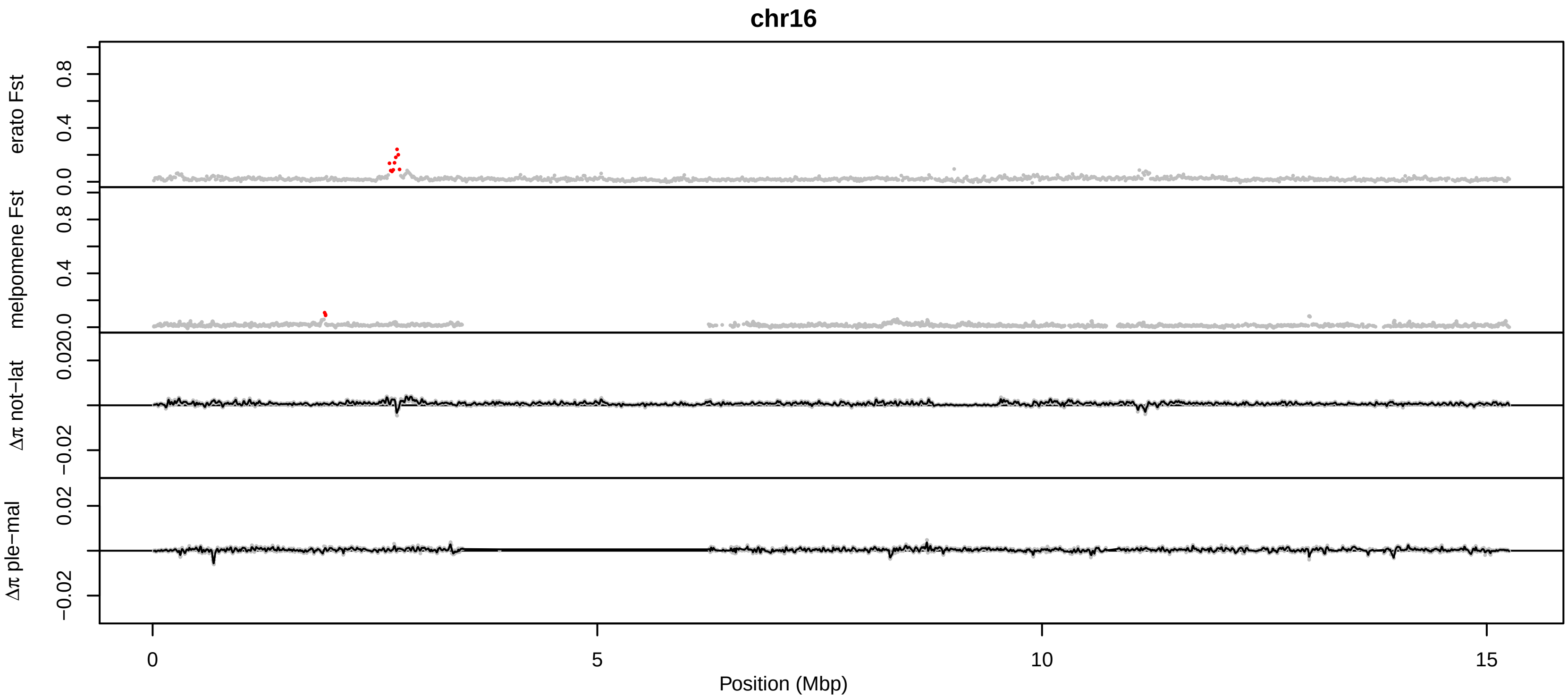
chr16
erato Fst
melpomene Fst
Δπ not−lat
Δπ ple−mal
0.0
0.4
0.8
0.020
−0.02
0.02
0
5
10
15
Position (Mbp)

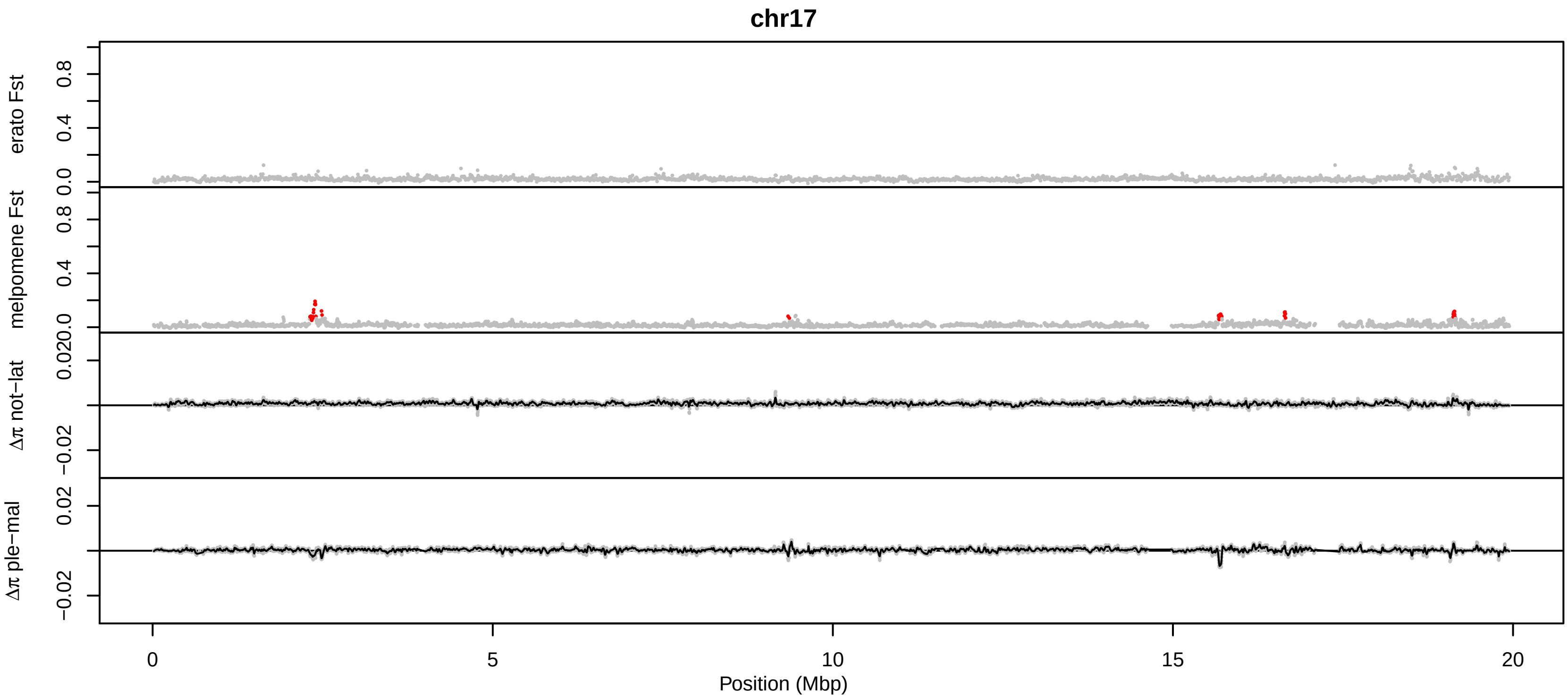

chr17
erato Fst
melpomene Fst
Δπ not−lat
Δπ ple−mal
0.0
0.4
0.8
0.020
−0.02
0.02
0
5
10
15
20
Position (Mbp)

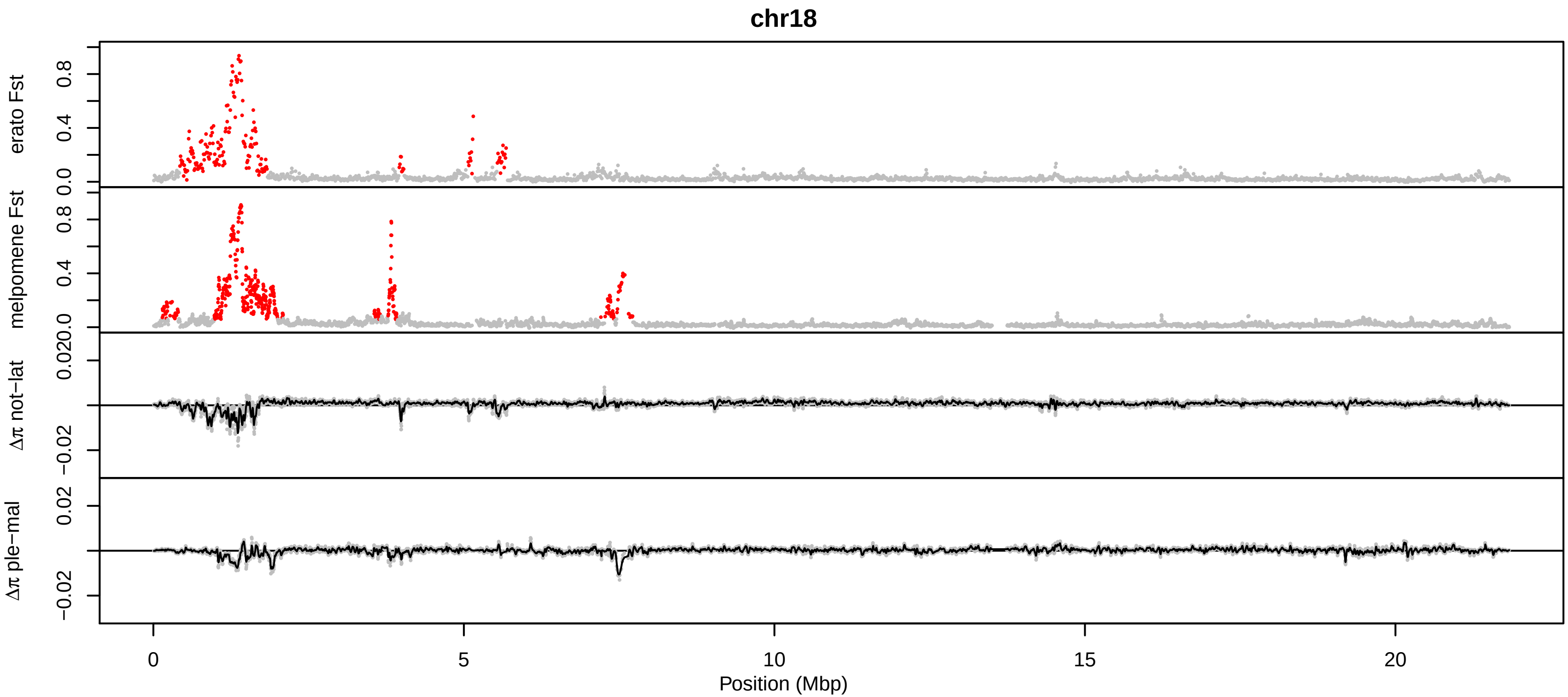
chr18
erato Fst
melpomene Fst
Δπ not−lat
Δπ ple−mal
0.0
0.4
0.8
0.020
−0.02
0.02
0
5
10
15
20
Position (Mbp)

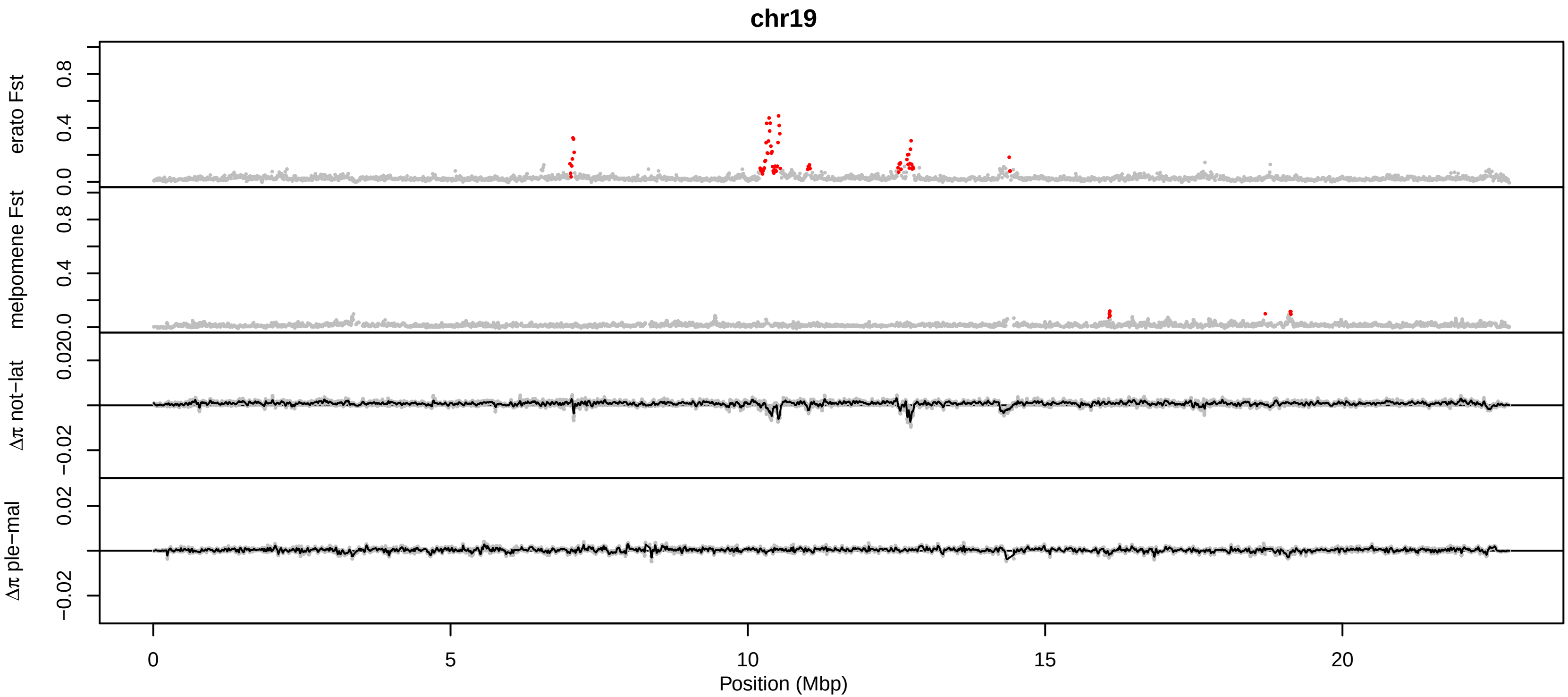

chr19
erato Fst
melpomene Fst
Δπ not−lat
Δπ ple−mal
0.0
0.4
0.8
0.020
−0.02
0.02
0
5
10
15
20
Position (Mbp)

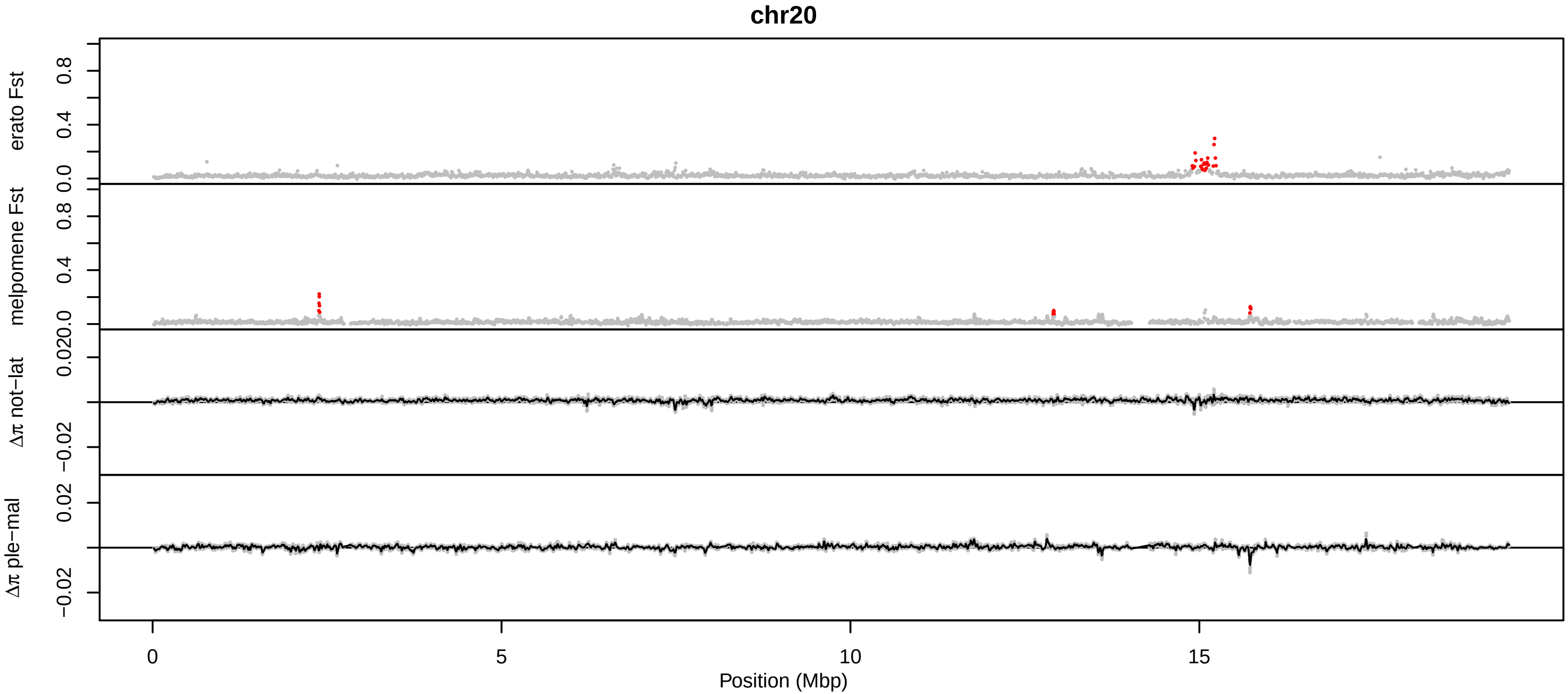

chr20
erato Fst
melpomene Fst
Δπ not−lat
Δπ ple−mal
0.0
0.4
0.8
0.020
−0.02
0.02
0
5
10
15
Position (Mbp)

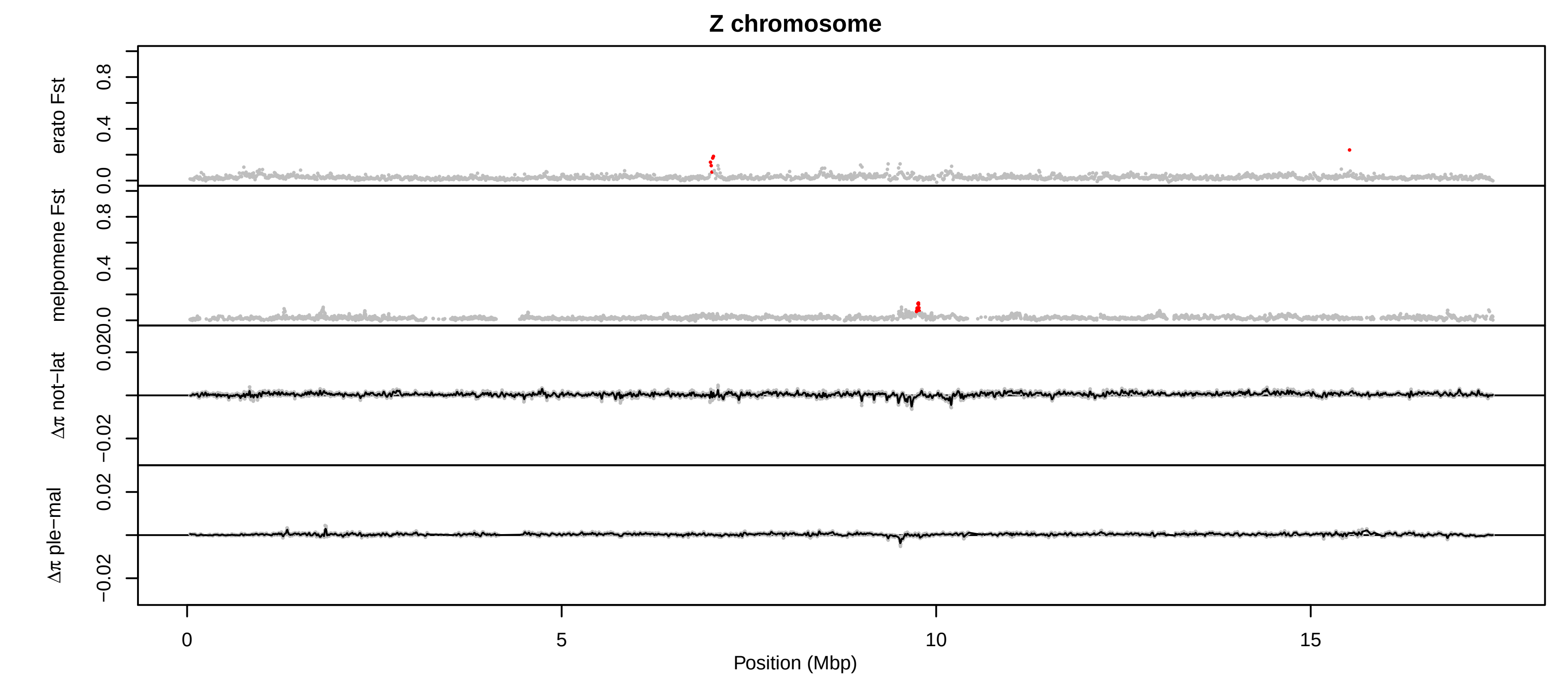

Z chromosome
erato Fst
melpomene Fst
Δπ not−lat
Δπ ple−mal
0.0
0.4
0.8
0.020
−0.02
0.02
0
5
10
15
Position (Mbp)
